# Consistent trade-offs in ecosystem services between land covers with different production intensities

**DOI:** 10.1101/621706

**Authors:** Carla Gómez-Creutzberg, Malgorzata Lagisz, Shinichi Nakagawa, Eckehard G. Brockerhoff, Jason M. Tylianakis

## Abstract

Sustaining multiple ecosystem services across a landscape requires an understanding of how consistently services are shaped by different categories of land uses. Yet, this understanding is generally constrained by the availability of fine-resolution data for multiple services across large areas and the spatial variability of land-use effects on services. We systematically surveyed published literature for New Zealand (1970 – 2015) to quantify the supply of 17 services across 25 land covers (as a proxy for land use). We found a consistent trade-off in the services supplied by anthropogenic land covers with a high production intensity (e.g., cropping) versus those with extensive or no production. In contrast, forest cover was not associated with any distinct patterns of service supply. By drawing on existing research findings we reveal complementarity and redundancy (potentially influencing resilience) in service supply from different land covers. This can guide practitioners in shaping land systems that sustainably support human well-being.

## I. INTRODUCTION

Human transformation of the Earth’s surface through land-use activities has reached an unprecedented magnitude, and constitutes a major driver of global environmental change (Turner, Lambin, & Reenberg, 2008; Steffen *et al*., 2015). Humans rely on resources appropriated through land use, however most of these practices affect the Earth’s ecosystems in ways that undermine human well-being (Foley *et al*., 2005). Continued population growth and increased per capita consumption of resources (Godfray *et al*., 2010) make it critical to find the ways to reconcile production and sustainability in land systems.

Ecosystem services (ES) offer a framework for addressing these complex issues by explicitly accounting for the benefits that ecosystems bring to society. Central to this framework is the idea that human well-being is underpinned by a diverse constellation of ES (MEA, 2005). Most of these ES are not accounted for in conventional land use planning and management decisions which, instead, tend to focus on the production of a single ES (e.g., provision of food or timber) (Robertson & Swinton, 2005; Rodríguez *et al*., 2006). By highlighting the importance of multiple over individual services, the ES framework encourages decision makers to prioritize long-term well-being over immediate economic reward (Guerry *et al*., 2015; Costanza *et al*., 2014).

Developing strategies that optimize ES across different land uses, or enhance multiple ES within a single type of land use (Lambin & Meyfroidt, 2011), relies on understanding the occurrence and interactions between different ES and their responses to management interventions. To this end, important efforts have been made to map and quantify ES supply (see Crossman *et al*., 2013; Groot *et al*., 2012; and Martínez-Harms & Balvanera, 2012 for reviews) and, more specifically, assess how different ES are enhanced synergistically or traded-off against each other (Nelson *et al*., 2009; Bateman *et al*., 2013). More recently, research on ES trade-offs and synergies has come together under the concept of ES bundles: groups of ES that repeatedly appear together in space and/or time (Raudsepp-Hearne, Peterson, & Bennett, 2010; Saidi & Spray, 2018). ES bundles can be examined in terms of the supply (Queiroz *et al*., 2015) and the demand (Ament *et al*., 2017) of ES. In either case, ES bundles can be used subsequently to identify any common processes or external factors driving different ES (Mouchet *et al*., 2014).

A systematic review of 51 studies on ES bundles revealed multiple approaches to bundling ES and the consequent difficulties in obtaining cross-site comparisons and generalizations of bundles and their drivers (Saidi & Spray, 2018). Moreover, even when the same methods, datasets and groups of ES were used to identify ES bundles and their relation to social-ecological variables in two regions, the results were highly inconsistent between regions (Spake *et al*., 2017) and, therefore, not generalizable to other locations. This inconsistency may result from the choice of ES indicators, socio-ecological variables and spatial units of analysis (Spake *et al*., 2017). Often, studies that examine ES bundles use administrative units (e.g., municipalities) as the scale at which ES are quantified (Saidi & Spray, 2018). However, administrative units can mask ES associations because they: 1) are variable in size (within the same hierarchical level), 2) occur at scales that are too coarse to capture the fine-scale processes linked to some ES, 3) encompass heterogeneous sets of land covers / land uses and 4) have boundaries that may cut across ecologically relevant units (Spake *et al*., 2017). Therefore, identifying consistent rules regarding ES bundles and their drivers requires tailored analyses that focus on finer scales, such as ES measured in individual plots within land cover types (Spake *et al*., 2017).

Here we directly test whether there are any general rules for the effect of land use on ES bundles by assessing the supply of multiple ES across land covers (as a proxy for land use) at a national scale. We systematically surveyed the published literature for New Zealand (1970 - 2015) to collate studies with quantitative evidence of how different land covers compare against each other in processes relating to the supply of one or more ES. For each study, we calculated standardized pairwise comparisons (expressed as log response ratios) of land covers in their supply of individual services. We used these ratios to conduct network meta-analysis for individual services and obtained, for each service, quantitative estimates of service supply from individual land covers.

With this comprehensive evidence base, we first discuss land cover effects on individual ES and then examine associations between ES to delineate any potential synergies and trade-offs arising from services that are best supplied by similar or different land covers. Similarly, we also examine associations between land covers based on the different ES they supply. We use this to detect: 1) any land covers that may be operating as “generalists” (i.e. supplying many ES) or “specialists” (i.e. supplying just a few ES) and 2) groups of land covers that supply similar profiles of ES (i.e. ES bundles sensu Raudsepp-Hearne *et al*., 2010). The latter includes services that are typically traded off against each other.

Subsequently, we test whether there are generalities regarding how categories of land cover influence ES bundles (i.e. sets of ES supplied consistently across more than one land cover) by testing for systematic differences between forested and non-forested habitats and between exotic-species-dominated production and native non-production land covers (note that we use the term production to refer to economic activity rather than primary production). If they exist, these differences would suggest that production/no production, forest/non-forest cover and native/exotic vegetation are attributes that drive changes in ES supply across multiple land covers. Previous research has shown that attributes of single land cover types can drive the value of multiple ES (Sutherland, Gergel, & Bennett, 2016) and trade-offs and synergies between ES (Felipe-Lucia *et al*., 2018). Duarte and colleagues (2018) also present evidence that landscape composition metrics (e.g., percentage of natural areas and of non-crop areas) affect some ES (water quality, pest regulation, pollination and disease mitigation); however, their analysis did not identify specific attributes of natural or non-crop areas that could shape ES supply. Our analysis extends these perspectives to include attributes shared by multiple land covers, which can potentially inform management decisions at broader scales and allow generalities across land covers. We conclude with an example of how our findings can be used to examine the effects of land cover trajectories or contrasting management decisions on landscape-scale ES trade-offs.

## II. METHODS

Unlike existing reviews and meta-analyses on ES (e.g., Howe *et al*., 2014; Malinga *et al*., 2015; Lee & Lautenbach, 2016; Nieto-Romero *et al*., 2014), our work does not collate existing ES assessments. Rather, we synthesize primary biophysical research that compares land covers in relation to a large variety of measures (which we term ‘ES indicators’) that indicate the supply of an ES, regardless of whether ES terminology was used. Despite the growing literature on ES (Chaudhary *et al*., 2015), our understanding of ES bundles, trade-offs and synergies has traditionally been impaired by the lack of, and costliness of obtaining, detailed spatial data on multiple ES from multiple land uses across landscapes (Andrew *et al*., 2015). This has led to the widespread approach of using expert or model estimates of ES per land use or land cover class as input for ES assessments (see Jacobs *et al*., 2015 for a review; Aldana Domínguez *et al*., 2019; and Chen, Chi, & Li, 2019 provide recent examples). Here, we propose an alternative approach that makes it possible to use primary data to study land cover and ES relations by capitalizing upon existing research across multiple disciplines. We use New Zealand as a case study because the high levels of endemic flora and fauna and relatively recent introduction of large-scale intensive agriculture make conservation-production tensions particularly acute, and necessitate conservation strategies that go beyond protected areas (Craig *et al*., 2000).

Our systematic review was structured according to the “Guidelines for Systematic Review in Environmental Management” developed by the Collaboration for Environmental Evidence (CEE, 2013). We searched the literature for quantitative comparisons of two or more land covers in the supply of one or more ES within New Zealand. Our ES definitions were adapted from the Millennium Ecosystem Assessment (MEA, 2005), with a total of 35 ES spanning across the provisioning, regulating, cultural and supporting categories (Supplementary Dataset 1). Despite the debates on whether the Millennium Ecosystem Assessment classification of ES leads to double counting of some services (Wallace, 2007; Fisher, Turner, & Morling, 2009), we have adopted it in this study because of its wide use and because our main interest was not to render a final valuation of ES (where double counting would be an issue), but instead to provide a comprehensive overview of the complete spectrum of direct and indirect benefits from ecosystems. Land uses, formally defined as the purposes to which humans put land into use (Dale *et al*., 2000), were captured in our research as land covers (Supplementary Dataset 2), since these include units that are not directly used by humans and, consequently, correspond more closely with the actual experimental or sampling units of many of the documents in our search.

### (1) Data collection, aggregation and calculation of effect sizes

Full details of the search and screening process are described in Supplementary Methods 1; here we present a brief outline. We searched the Scopus database for titles, abstracts and keywords with at least one match in each of the 3 components that structured our search: 1) “New Zealand”, 2) land cover and land use terms and 3) ES terms (see Supplementary Methods 2 for the full search phrase). Land cover terms included all possible variations of “land use” and “land cover” as well as the names of specific land use and land cover types (both generic and specific to New Zealand). The ES component drew upon the names of each service (and possible variations of these) but also included vocabulary describing processes and conditions that could reflect their supply at the site scale akin to individual land cover units. The search was finalized in December 2014, and was constrained to include documents published from 1970 onward, to be comparable with current land use regimes in New Zealand (MacLeod & Moller, 2006).

Our keyword search yielded 9,741 references. An initial automated screening process reduced these to 4,373 publications by removing references that only mentioned a single type of land cover or land use in their title, abstract and keywords. We excluded these studies because measures of ES supply from single land covers could not be standardized in a way that would make them comparable across studies *and* compatible with the standardized land cover comparisons of ES supply that informed the rest of our meta-analysis.

Publications with 2 or more land cover terms were scanned using Abstrackr, an interactive machine learning system for semi-automated abstract screening, often used in medical meta-analyses (Wallace *et al*., 2012). By learning from the abstracts or words that a user identifies as relevant during the screening process, Abstrackr can predict the likely relevance of unscreened abstracts and effectively assist in the exclusion of irrelevant ones (more details in Supplementary Methods 1).

Abstract screening yielded 914 relevant papers, which were passed on to a team of four reviewers for full-text assessment and data extraction. Studies that did not have replicated observations (as defined in Supplementary Methods 1) for any land covers were discarded, whereas studies that contained replication on some, but not all, of the land covers were kept and only data on the replicated land covers were extracted. Although we only included terrestrial land covers, ES supplied by land but linked to a water body were included in our analysis. Full details of how the full-text selection criteria were applied can be found in Supplementary Methods 3. In total, we extracted data from 133 studies that met all inclusion criteria (see Supplementary Dataset 4 for bibliographic details of each study).

Information on the land covers, quantitative measures of ES supply, experimental design and bibliographic details for each study was collated in a database. To allow for comparability across studies, individual land covers described in each study were matched to the nearest category in New Zealand’s Land Cover Database - LCDB (Thompson, Grüner, & Gapare, 2003). This classification system includes forest, shrubland and grassland areas of either predominantly native or exotic vegetation, as well as cropland and more artificial surfaces such as built-up surfaces and mining areas (Supplementary Dataset 2).

Often, the same quantitative measure of ES supply obtained from a study (indicators, presented in Supplementary Dataset 3) would be relevant to more than one ES. This reflects the overlaps that exist between different ES (e.g., soil structure plays a role in both soil formation and regulation of water timing and flows), and the multiple values that humans can receive from a given ecosystem process. We therefore decided to assign each indicator to as many ES as it was relevant to, and use this allocation in our main analysis. However, to understand the influence on our results of sharing indicators between ES, we also conducted the same analysis with each indicator assigned to only one ES. In Supplementary Results 5 we present the results of this analysis.

For each indicator - ES combination we defined the general direction of the relationship by determining whether larger values of the indicator would generally reflect an increase or decrease in ES supply. This was done because the majority of the studies in our meta-analysis did not explicitly use ‘ecosystem services’ terminology. Instead, they measured environmental or ecological variables that could be used as indicators of ES supply, provided a conceptual link could be defined between the indicator (e.g., annual water discharge of a catchment) and the corresponding ES (provision of freshwater). When we could not readily assign indicators to ES or determine the direction of the indicator - ES relationship we consulted with experts with specialized knowledge of the field related to each indicator (see Acknowledgements). Although we recognize that the relationship between an indicator and a ES may be non-linear (e.g., pollination services may saturate with large numbers of pollinators), in most cases it was not possible to establish a clearly defined non-linear function, so we assumed a linear relationship for all indicators. Supplementary Dataset 3 provides an overview of the relations we defined between each indicator and ES.

Unique identifiers allowed us to define individual studies, regardless of whether they were within a publication that included more than one study or across different publications (Supplementary Methods 1). Multiple measures from within the same replicate site were aggregated into a single value per replicate (see Supplementary Methods 1 for details). Methods for standardizing measures of variance are presented in Supplementary Methods 4.

We obtained a final database with information on 457 ES indicators among 2,943 pairwise comparisons of land covers from 133 studies. A log response ratio was used as the effect measure for comparing pairs of land covers within each study, and was standardized such that larger values always represented greater ES supply in the numerator land cover relative to the denominator one (see Supplementary Methods 1 for this standardization and log response ratio variance calculations).

Studies with more than one indicator of a given ES were aggregated to have the same weight as studies with only a single indicator (this was based on either the mean log response ratio across multiple indicators or the single indicator represented in all land covers of a study, details in Supplementary Methods 1). Subsequently, the total number of land cover comparisons in our final dataset of 133 studies was reduced from 2,943 to 920 comparisons for individual ES within single studies (See Supplementary Dataset 5 for an overview of the final data).

### (2) Data analysis

Data analysis was conducted as a two stage process: we first examined the supply of each ES by different land covers, and then assessed the relationships among land covers in terms of multiple ES. For the first stage, we conducted a separate network meta-analysis (Salanti, 2012) for each ES. While conventional meta-analysis compares 2 treatments at a time (using direct comparisons from each study), a network meta-analysis can compare multiple (i.e. 3 or more) treatments simultaneously. This is achieved by using both direct evidence (studies comparing pairs of treatments) and indirect evidence derived from linking common treatments across different studies in a network of evidence (Salanti, 2012). For example, if some studies show that land cover A is better than B in supplying an ES, and others provide direct evidence that B is better than C, then a network meta-analysis allows us to make the inference that A will also be better than C. We therefore used network meta-analysis to compare, for each ES, a wide array of land covers across different studies, even though we did not have data for direct comparisons among all combinations of land covers.

We conducted our network meta-analyses with the R package Netmeta (Schwarzer *et al*., 2019), which offers a frequentist approach to calculate point estimates (and their corresponding 95% confidence intervals) of the effect of the different land covers on the supply of individual ES. Estimates were expressed as the log response ratio of each land cover relative to a reference land cover: high producing exotic grassland. We selected this land cover as our reference, because it was the only land cover that was represented across all ES in our dataset (and would therefore allow us to compare our results across ES at a later stage).

In *Netmeta*, we used a random effects meta-analytic model to generate estimates and confidence intervals from which we then calculated probability scores (*P*-scores; Rücker & Schwarzer, 2015) on how different land covers ranked in the supply of each ES. Estimates, confidence intervals and *P*-scores then allowed us to construct, for each ES, a so-called forest plot or blobbogram (sensu Lewis & Clarke, 2001) to compare different land covers in their ES supply.

Bundles, trade-offs and synergies in land cover effects across the whole suite of ES were then examined using hierarchical clustering of the network meta-analytic estimates. For this, we constructed a land cover by ES matrix (Fig. S44, Supplementary Results 3) using the estimated log response ratios of each land cover (relative to the high producing exotic grassland reference) in each ES, as determined with the individual network meta-analyses. Missing values in this matrix resulted from sets of land covers for which we had no information on a given ES or could not infer the corresponding ratios.

For analysis, we selected subsets of this matrix with no gaps and the largest possible number of total cells. This resulted in two data subsets: a matrix of nine ES by eight land covers and another matrix with nine land covers by eight ES. The matrix with nine ES was rotated to have ES as rows (land covers as columns) and used to compare ES in terms of the land covers that supply them. This allowed us to identify ES bundles (sets of ES supplied similarly across multiple land covers), synergies in ES supply, and ES that would likely be traded off with one another in land-use decisions. The matrix with nine land covers was used to compare land covers (to identify redundancy) in the supply of eight ES. This allowed us to explore how land-cover differences influence ES bundles.

We calculated a dissimilarity matrix from each of these matrices using the *daisy function* of the *cluster* package for R (Maechler *et al*., 2019) with Euclidean distances. For the rotated matrix with nine ES, distances were based on ES observations for each land cover, while for the matrix with nine land covers, distances were based on land cover observations for each ES. We applied hierarchical clustering (using the R *hclust* function; R Core Team, 2019) to each of the distance matrices and constructed dendrograms on how different land covers or ES compared against each other. Following Raudsepp-Hearne *et al*. (2010), we also used these distance matrices to conduct k-means cluster analysis (with the *kmeans* function in R; R Core Team, 2019) and identify groups of land covers and ES exhibiting similar behavior. In each case, the number of clusters was determined using a scree plot (Figs. S3 and S4, Supplementary Methods 5).

Finally, we used our distance matrices with nine land covers to test hypotheses on whether broad categories of land covers explained the trends observed in the corresponding clustering. Specifically, land covers were grouped under two categorical variables, one denoting the presence/absence of forest cover and another separating production land covers, dominated by exotic vegetation cover, from those with no production activities. Originally, we expected to compare land covers with a native vs. exotic vegetation cover separately from production vs. no production. However, we omitted the former category because, except for one, all land covers with exotic vegetation were production and all native covers had little or no production. We used a permutational multivariate analysis of variance (PERMANOVA) to test whether these variables or their interaction explained between-land-cover differences in the supply of multiple ES.

PERMANOVA analyses were conducted using the *adonis* function of the vegan package in R (Oksanen *et al*., 2019). Variables are added sequentially in the *adonis* algorithm. To be conservative, we performed the PERMANOVA twice and swapped the order of the variables in the second iteration, so that each variable was tested second, after controlling for any collinearity with the other predictor (i.e. adjusted sums of squares). The *betadisper* function of the *vegan* package was used to test the assumption of multivariate homogeneity of group dispersions, and all tests met this assumption. Table S4 (Supplementary Methods 5) presents the land cover categories used in these analyses.

## III. RESULTS

### (1) Data coverage

From our systematic survey, we identified a total of 133 studies that were relevant to our analysis and matched our selection criteria. Overall, these studies contributed data on 17 different ES, 25 land cover types and 457 measures (which we term ‘ES indicators’) on ES supply. All four of the Millennium Ecosystem Assessment ES categories (supporting, provisioning, regulating and cultural services; MEA, 2005) were represented within our dataset. However, most studies examined supporting and regulating services, with 115 and 110 studies, respectively. Only 44 studies presented data on provisioning services and four on cultural ones. All of the ES in the supporting category (habitat provision, nutrient cycling, soil formation, water cycling and primary production) are represented in our database. Only four land cover comparisons had more than 20 studies (high producing exotic grassland vs. exotic forest, indigenous forest vs. high producing exotic grassland, short-rotation cropland vs. high producing exotic grassland and exotic forest vs. indigenous forest); whereas the remaining land cover pairs were represented by 10 or fewer studies each. Further details on the number of studies per land cover comparison and per combination of ES and land cover are available in Supplementary Results 1.

### (2) Land cover effects on individual ES

There were consistent trends in the supply of multiple services by specific land cover types, but also great variability in the supply of some services. An overview of the evidence base (number of studies, types of ES indicators and network of land cover comparisons) and the outcomes of the individual network meta-analyses for each of the 17 ES in our database is presented in Supplementary Results 2. In this supplement, we use forest plots (sensu Lewis & Clarke, 2001), see Fig. S8, Supplementary Results 2 for an example) to show the main results of the meta-analysis, i.e. how different land covers compare against each other in their supply individual ES. Specifically, the values in these plots are given as log response ratios which express the overall estimates of service supply by individual land covers relative to a reference land cover (high-producing exotic grassland).

For several ES, the positive log response ratio estimate and narrow confidence intervals in the forest plots (Figs. S8, S17, S19, S38, Supplementary Results 2) reveal that land covers with native vegetation cover (i.e. broadleaved indigenous hardwoods, indigenous forest, manuka/kanuka, matagouri or grey scrub and, in many cases, tall tussock grassland) tended to rank higher in ES supply than the more intensive high-value production land covers (particularly short-rotation cropland and high-producing exotic grassland). Regulation of water timing and flows, water purification, freshwater provision and disease mitigation conformed to this general pattern. In these services, low producing grasslands (which comprise a mix of exotic and native vegetation) and exotic forests also perform relatively well and always rank within the top half of all land covers.

For habitat provision (Fig. S13, Supplementary Results 2) the difference between land covers with native vegetation and production systems was less important than the presence of forest vegetation cover. For this service, most land covers with forest vegetation (exotic forest, broadleaved indigenous hardwoods and indigenous forest) ranked higher in their estimates of ES supply than those with open covers (short-rotation cropland, tussock, low and high producing grasslands) or deciduous hardwoods.

Meanwhile, primary production tended to be highest under production systems (e.g., croplands, exotic forest, and high-intensity grassland) and lower in land covers with low or no production (e.g., low producing and tussock grasslands, indigenous forest), rather than differing between forested and open covers. However, these trends were not statistically significant due to the wide and overlapping confidence intervals (Fig. S23, Supplementary Results 2).

Importantly, these results indicate that no single land cover supplies all ES at a maximal level. Indigenous forests ranked high in the supply of many ES (particularly habitat provision, freshwater provision, disease mitigation and global climate regulation - Supplementary Results 2). However, in some ES they were outperformed by other land covers such as tall tussock grasslands (which were well suited to water purification; Fig. S19, Supplementary Results 2) and advanced successional forest (broadleaved indigenous hardwoods, which ranked high in regulation of water timing and flows, nutrient cycling and habitat provision; Figs. S8, S11 and S13, Supplementary Results 2). Therefore, multiple land covers will be required within the landscape to ensure the supply of multiple ES.

The forest plots in Supplementary Results 2 for primary production (Fig. S23), erosion control (Fig. S27), pest regulation (Fig. S30), waste treatment (Fig. S32), capture fisheries (Fig. S34), ethical & spiritual values (Fig. S36), pollination (Fig. S41) and regional & local climate regulation (Fig. S43) all present wide, overlapping confidence intervals for all or most of their estimates. This suggests statistically non-significant differences in the supply of these services among land covers. For some services, this could be due to small evidence bases, either in terms of few studies or few comparisons for specific land cover pairs within the network of land cover comparisons that inform the meta-analysis. In the case of erosion control, where the evidence base is formed by 22 studies (Supplementary Results 2 - Erosion control), overlapping confidence intervals in the land covers with the greatest number of comparisons (which would therefore be expected to have lower variance) still expressed high variability in ES supply, suggesting that other factors besides land cover (e.g., slope, soil type) likely account for the differences in erosion control across the sites in all 22 studies.

### (3) Land cover effects across multiple ES

We explored how the above trends in the supply of individual services translate into bundles, synergies and trade-offs among ES. For this we conducted multivariate analyses to simultaneously explore differences in the supply of multiple services across land covers (see Methods - Data analysis). These analyses allowed us to examine whether groups of ES responded similarly to differences in land cover and, conversely, whether groups of land covers played a similar role in the supply of multiple ES.

#### (a) Differences among ES from the land covers that supply them

For this analysis we used a matrix of eight land covers by nine ES to identify clusters of ES based on how they are supplied by different land covers. We identified a total of five clusters, three of which were formed by only one ES while the remaining two had two and four ES each (Fig. 1). This suggests that more than half of the nine ES in this analysis are supplied in a distinct way by different land covers, and reinforces the notion that multiple land covers are required to supply a range of ES. Moreover, the separation of services into clusters of one to two also suggests that their supply is traded-off across land covers. This trade-off is acute for water-related services; most of these tend to occupy distinct spaces within the dendrogram, with water cycling standing apart from all other ES, water purification and freshwater provision in a separate cluster, and regulation of water timing and flows in a single branch close to global climate regulation and nutrient cycling (Fig. 1). The trade-off between water cycling and regulation of water timing and flows is probably because land covers that allow for increased runoff and present low water retention (such as harvested forests, croplands and built-up areas) deliver more of the water cycling service than land covers that promote soil water storage and, consequently, perform better in regulating water timing and flows (e.g., broadleaved indigenous hardwoods, indigenous forests and low producing grasslands). Freshwater provision and water purification form a cluster because the water quality aspect of their supply was assessed with the same indicators for both services (Supplementary Dataset 3) and, in both cases, greater service supply came from land covers contributing to enhanced water quality (such as tall tussock grassland and indigenous forest; Figs. S17 and S19, Supplementary Results 2).

**Fig. 1:**
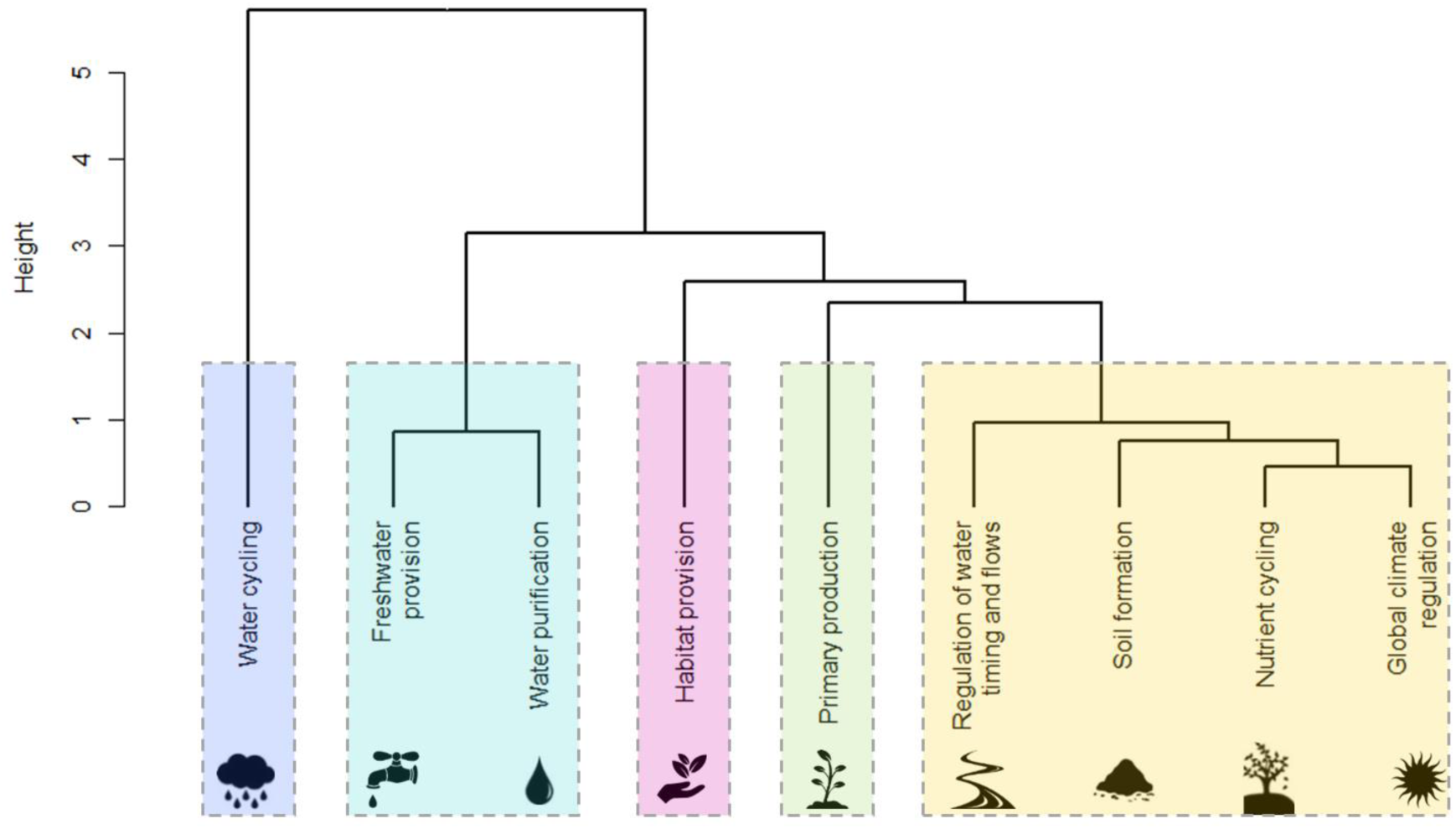
Hierarchical clustering of ES. Services within the same box form a cluster (as determined by k-means cluster analysis) and are therefore supplied similarly across eight land covers (low producing grassland, tall tussock grassland, high producing exotic grassland, short - rotation cropland, indigenous forest, exotic forest, harvested forest and orchard, vineyard and other perennial crops). A greater separation between the branching points for clusters along the height axis indicates greater dissimilarity among clusters in the extent to which they are supplied by the eight land covers included in the analysis.

In contrast to the water-related ES, those more closely linked to the soil system (nutrient cycling and soil formation) are found closer to each other in Fig. 1, and appear to be delivered similarly across land covers (Figs. S11 and S15, Supplementary Results 2). In our analysis, global climate regulation falls under this broad group of services and is closely linked to nutrient cycling (Fig. 1). This is likely due to the indicators shared by both (Supplementary Dataset 3) and a gap in our database with respect to the contribution of vegetation and livestock in greenhouse gas fluxes. In New Zealand, these contributions are well studied within a given land cover, but the lack of comparisons across land covers and uses prevented us from making a more comprehensive quantification of how this service is supplied.

### (b) Differences among land covers in their supply of services

Our analysis of how land covers compared against each other in their supply of ES was based on a matrix of nine land covers by eight ES. We found a gradient of land covers that separates those with lower production from the high value production systems (Fig. 2).

**Fig. 2:**
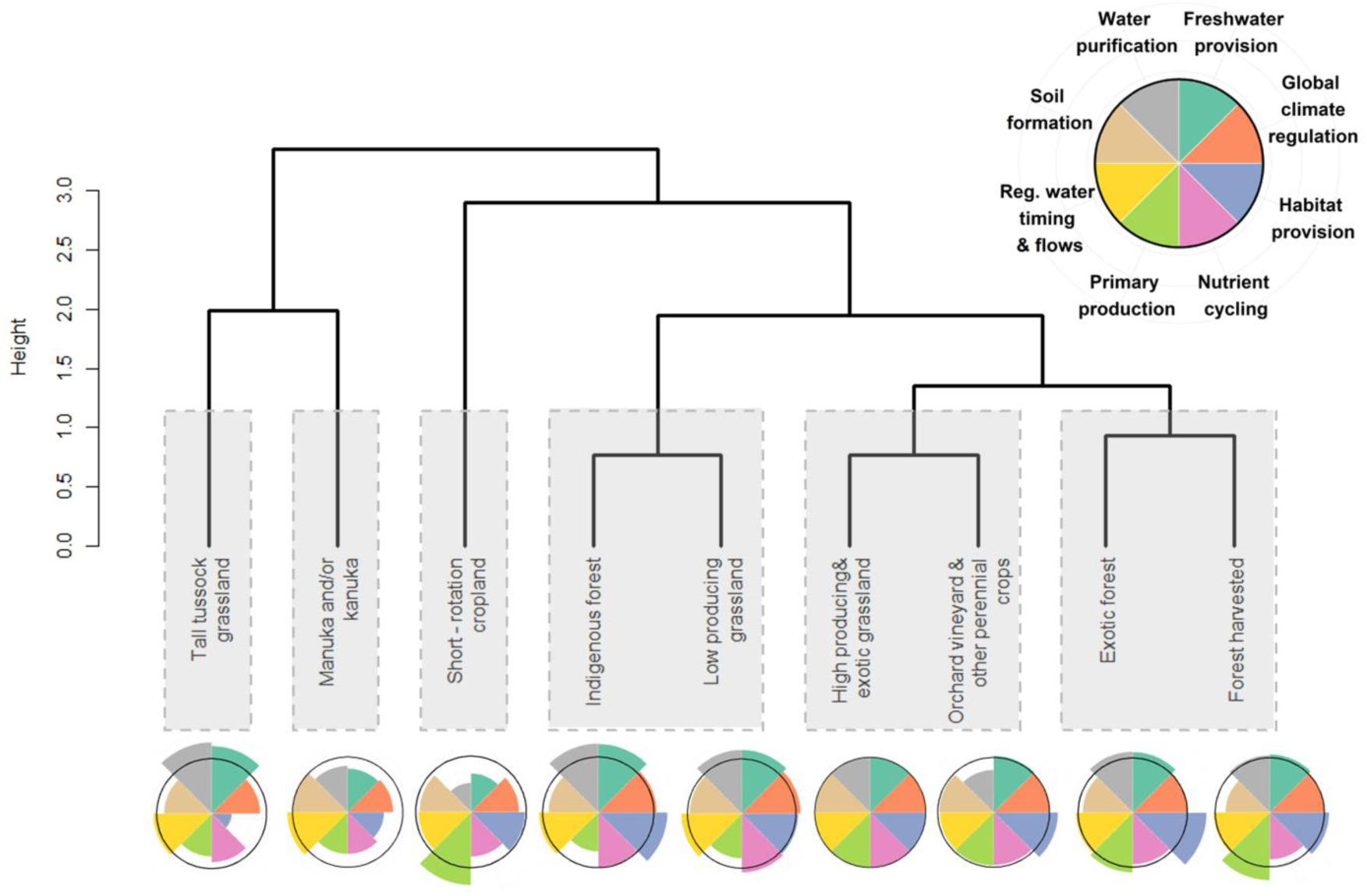
Hierarchical clustering of land covers. Boxes enclose land covers that exhibit a greater similarity in their supply of eight ecosystem services (habitat provision, primary production, freshwater provision, soil formation, nutrient cycling, water purification, global climate regulation and regulation of water timing and flows). In contrast, land covers that merge at a greater height have a greater dissimilarity in their service supply. The flower diagrams at the bottom illustrate how each land cover supplies each of the eight ES, with longer petals indicating a greater supply of an ES. For comparison, the black ring around each flower diagram marks the supply from high producing exotic grassland, the land cover used as reference in our meta-analysis.

Land covers with high production value and dominated by exotic vegetation cover (croplands, high producing exotic grassland, exotic and harvested forests) occupied separate clusters from those with low or no production and primarily native components in their vegetation cover (tall tussock and low producing grassland, manuka and/or kanuka and indigenous forest). Likewise, with the exception of low producing grassland, land covers with forest vegetation cover occupied separate clusters from those with a more open vegetation cover.

The clusters with single land covers in Fig. 2 appear to specialize in supplying high levels of only one to three of the nine ES used in the analysis. Tall tussock grassland supplies high levels of water purification and freshwater provision, while manuka and/or kanuka (a successional land cover) is noted for soil formation and regulation of water and timing of flows; short-rotation cropland ranks high in supplying primary production. In contrast, the three clusters with pairs of land covers in Fig. 2 exhibit a more uniform supply of the different ES. Nevertheless, each of these three clusters also appears to supply a distinct ES bundle. The cluster formed by exotic and harvested forests supplies a bundle with high biomass production and habitat provision while the cluster formed by indigenous forest and low producing grassland supplies a bundle specializing in purifying, providing and regulating the flow of water. Lastly, the cluster formed by high producing exotic grassland and orchard, vineyard and other perennial crops appears to supply even (yet not necessarily high) levels of all ES.

Greater differences in ES supply can be inferred from the larger differences in the height at which clusters separate from each other (Fig. 2). Consequently, in Fig. 2, the clusters with two production land covers (harvested and exotic forest plus high producing exotic grassland and orchard, vineyard & other perennial crops) are similar in their supply of ES but differ from the cluster with indigenous forest and low producing grassland. In turn, these three clusters with pairs of land covers are more similar to each other (indicated by the lower branch point) than they are to the clusters with single land covers. The clusters with pairs of land covers are also more close to the short-rotation cropland than to tall tussock grassland and manuka and/or kanuka, which are more similar to each other than they are to the rest of the land covers.

The trade-off in service supply between production and non-production land covers was statistically significant (PERMANOVA, Pseudo *F*_1,6_ = 3.064, partial *R*^2^= 0.312, *p* < 0.05; detailed results in Supplementary Results 4). The assumption of homogeneous dispersion between both groups was met (*F*_1,8_ = 0.718, *p* > 0.05), suggesting that neither supplies a greater range of ES among its different land covers. Conversely, the separation between forested and non-forested land covers did not significantly explain the distribution of land covers in ES space (Pseudo *F*_1,6_ = 0.536, partial *R*^2^= 0.055, *p* > 0.05; see also Supplementary Results 4) nor did the interaction between forested/non-forested and production/non-production (Pseudo *F*_1,6_ = 1.159, partial *R*^2^= 0.118, *p* > 0.05; Supplementary Results 4).

## IV. DISCUSSION

We have synthesized over 40 years of quantitative primary evidence on the ES supplied by different land cover types at a national scale, and used this to identify bundles and trade-offs among ES, as well as general land cover characteristics driving these associations. Overall, we found strong evidence that high-value production land covers supplied a different set of non-market services than all the land covers with low or no production and native elements in their vegetation cover. Together, land covers with low or no production outperformed the production ones in supplying several supporting and regulating ES (e.g., freshwater provision, disease mitigation and regulation of water timing and flows). In contrast, most production land covers specialized in supplying primary production.

Interestingly, forest cover (native or exotic) was not associated with significant differences in the suite of services supplied. Instead, we observed a close affinity between land covers with contrasting forest covers (e.g., between low producing grassland and indigenous forest and between exotic forests and high producing exotic grasslands) in their supply of several ES including water purification and regulation of water and timing of flows. Only for habitat provision did we observe that land covers with a forest cover (indigenous forest and exotic forest - harvested and unharvested) performed better than those without a forest cover in service supply.

In New Zealand, production land covers are dominant, with exotic forests, high producing exotic grasslands, croplands, and orchards/vineyards occupying 42% of the country’s terrestrial area in 2012 (Landcare Research, 2015). Our assessment, like other ES assessments elsewhere (Costanza *et al*., 2014), shows that decisions on ecosystem management (such as those leading to the dominance of production land covers) reflect preferences for a set of ES over others. Specifically, the trade-offs we find between production and low or no production land covers illustrate how the preference for ES with a high market value and short-term returns occurs at the expense of ES that have a non-market value but are essential for sustained, long-term human well-being (Rodríguez-Loinaz, Alday, & Onaindia, 2015).

The above findings resonate with the recommendations of Foley and colleagues (2011) with respect to halting indiscriminate expansion of agriculture into sensitive ecosystems. However, our findings also suggest that, at the landscape scale, the trade-offs between the ES supplied by production and non-production land covers are not solved with a single land cover. Even for the ES that were best delivered by land covers with no production, we did not find evidence of a single land cover consistently performing better than the rest in the supply of all ES. Therefore, a landscape with a mosaic of these land covers is more likely to offer a broader suite of ES than one dominated by large extents of any single low or no production land cover (Fischer, Lindenmayer, & Manning, 2006; Law *et al*., 2015).

Thus, we support earlier recommendations to extend beyond the dichotomy of conservation vs. production land into a more a comprehensive management (Grau, Kuemmerle, & Macchi, 2013; Tscharntke *et al*., 2005). Such management could, for example, contemplate the extension or restoration of under-represented native land uses at strategic sites where intensive use is not matched by increased production yield, to promote the supply of critical ES or broaden the existing suite. To this end, management will need to be informed by a comprehensive understanding of how ES can scale up from individual land use units and how the relative sizes of different land use units within a landscape can affect ES supply.

Our analysis shows that low-intensity production land covers that retain some native vegetation (i.e. the low producing grasslands in our dataset) can approach native land covers (indigenous forests) in terms of overall ES supply. These low-intensity production land covers demonstrate that production and a suite of other ES can be jointly delivered, providing empirical support to the notion of managed ecosystems with “restored” ES proposed by Foley *et al*. (2005). Importantly, we identified great variability in how land covers supplied certain ES, despite there being high replication in our evidence base for these effects (e.g., erosion control by high producing exotic grasslands, indigenous and exotic forests). This suggests that local environmental conditions (e.g., slope) and management practices can significantly alter how a given land use affects ES supply (Felipe-Lucia *et al*., 2018). In turn, this implies some potential to improve ES supply by adjusting management practices within specific land uses (Guerra & Pinto-Correia, 2016; Pang *et al*., 2017) or better incorporating local environmental conditions into land-use decisions. Within individual land uses, decisions on which practices to adopt will require detailed research on the effects of different management regimes on ES supply (Guerra & Pinto-Correia, 2016; Maseyk, Dominati, & Mackay, 2018), as well as an understanding of the extent to which the plasticity in ES supply is constrained (or favored) by environmental factors.

A critical challenge in applying the ES framework to spatial and environmental planning is understanding the extent to which different land uses affect ES supply (Braat & Groot, 2012). The uneven coverage of different ES that we observed in the literature reflects both the variable difficulty of quantifying the supply for different ES and the likely relevance of comparing the supply of certain ES among land uses. Within our dataset, supporting and regulating ES are best represented. In the global literature, regulating ES are also the most commonly quantified and mapped category, however, they are usually followed by provisioning ES, while the evidence on supporting ES is scarce (Howe *et al*., 2014; Malinga *et al*., 2015; Martínez-Harms & Balvanera, 2012; Crossman *et al*., 2013). The limited representation of provisioning ES in our dataset possibly occurred because most provisioning ES (e.g., milk, timber) are linked to single or few land covers and, consequently, are unlikely to be compared across land covers. Such services, however, enter the market directly and can be more readily quantified in monetary terms. In contrast, the supporting and regulating ES that predominate in our dataset usually translate to externalities in the context of production systems, and are likely more readily quantified through biophysical indicators than monetary units (Howe *et al*., 2014; Czúcz *et al*., 2018).

Cultural ES are poorly represented in our database, with the few indicators for this category all being shared with the capture fisheries provisioning service, because they pertain to eels, which are of cultural significance to Māori in New Zealand. Cultural ES have non-material and ideological dimensions that are not readily quantified and, thus, are not well represented even within the emerging body of specialized literature on ES supply assessment (Hernández-Morcillo, Plieninger, & Bieling, 2013). Moreover, it has been suggested that cultural ES escape the instrumental value domain present in the ES framework. Instead, they fall under the relational domain, whereby value is not solely defined in terms of the direct benefits derived from an ecosystem, but also in terms of the social webs of desired and actual relationships constructed around that ecosystem or its components (Chan *et al*., 2016). Consequently, for these ES, a quantitative approach like ours should be complemented with assessments that address the relational dimensions of the values people hold for the natural elements in different land uses to better represent their importance in a cultural context (Lyver *et al*., 2017).

Individual ES are defined to encompass distinct processes and values, but these are often quantified by overlapping sets of indicators (Czúcz *et al*., 2018). For example, in our dataset indicators from water and soil pertained to more than one ES (e.g., water purification and provision of freshwater both share indicators of water quality, while erosion control and soil formation share indicators on soil stability). Ecosystem service indicators can also occupy different positions in the spectrum connecting the supply and demand end of ES (Villamagna, Angermeier, & Bennett, 2013). Here we have focused exclusively on the supply end and, more specifically, on the capacity of land covers to provide ES rather than on their actual flow or delivery as benefits perceived by a specific group of individuals.

Since the Millennium Ecosystem Assessment was released, there have been initiatives to redefine ES and their categories (TEEB, 2010; CICES, 2018). Here we argue that future work in determining how to best quantify ES, their potential and realized delivery, and their spatio-temporal variation, will be at least as important as refining their taxonomy. Furthermore, if a focus on quantifying ES should reveal aspects of services that are best left unquantified (such as the relational domain of cultural ES), this could also lead to the development of alternative ways of assessing those ES, which could then be applied in combination with quantitative approaches like the one we have developed here. Recent developments, like the concept of nature’s contributions to people and the framework for their assessment proposed by Díaz and colleagues (2018), provide an opportunity for reconciling these issues.

Our work suggests that there is great potential in using existing data for assessing ES bundles and interactions more cost-efficiently than through direct field observation. Yet, an important caveat to our approach stems from underlying factors that are correlated with land use and impact the supply of certain ES. For example, since land uses such as forestry and natural habitats are frequently found on steep slopes, this physical characteristic will likely influence erosion control in a way that co-varies with land cover. At the most extreme end, some ES may not be related to land cover, but rather respond to other spatially variable factors (e.g., aesthetic values from housing location on hillsides). These factors were beyond the scope of our work, as we did not separate the effects of spatial factors from those of land cover. In fact, one could argue that land use is not selected independently from the local environment, so these factors are a frequent (though not universal) component of any land use and its influence on ES. Nevertheless, future approaches may benefit from examining how these factors affect the between- or within-land-use differences in ES supply. This distinction would allow a shift from comparisons across locations (as we examined here), which allow comparisons of existing landscapes, to the predicted impacts of land use change on ES at any location. However, such predictions would also need to incorporate legacy effects of past land uses, as these can have enduring consequences on ecosystem functioning (Dallimer *et al*., 2015; Perring *et al*., 2016).

Our method for using existing data to assess bundles, trade-offs and synergies in ES supply across land covers can facilitate the comparison of entire landscapes, for example, by projecting land covers or land uses into multidimensional ES-supply space (Fig. 3). This mapping could reveal two key characteristics for land-use planning: 1) land covers/uses that cluster together, and thus exhibit redundancy (and potentially resilience) in ES supply, or 2) land covers/uses that occur at opposite extremes of ES-supply space, and are therefore likely to exhibit complementary roles in their service supply (as ES are traded off between them). In addition, the total hyper-volume occupied by all land covers/uses in this multidimensional ES-supply space (ordination plots in Fig. 3) can indicate the diversity of ES supplied by all land covers/uses within a given landscape (analogous to interpretations of species in trait space; Laliberte & Legendre, 2010), which could be used in comparisons of existing landscapes or future scenarios.

**Fig. 3:**
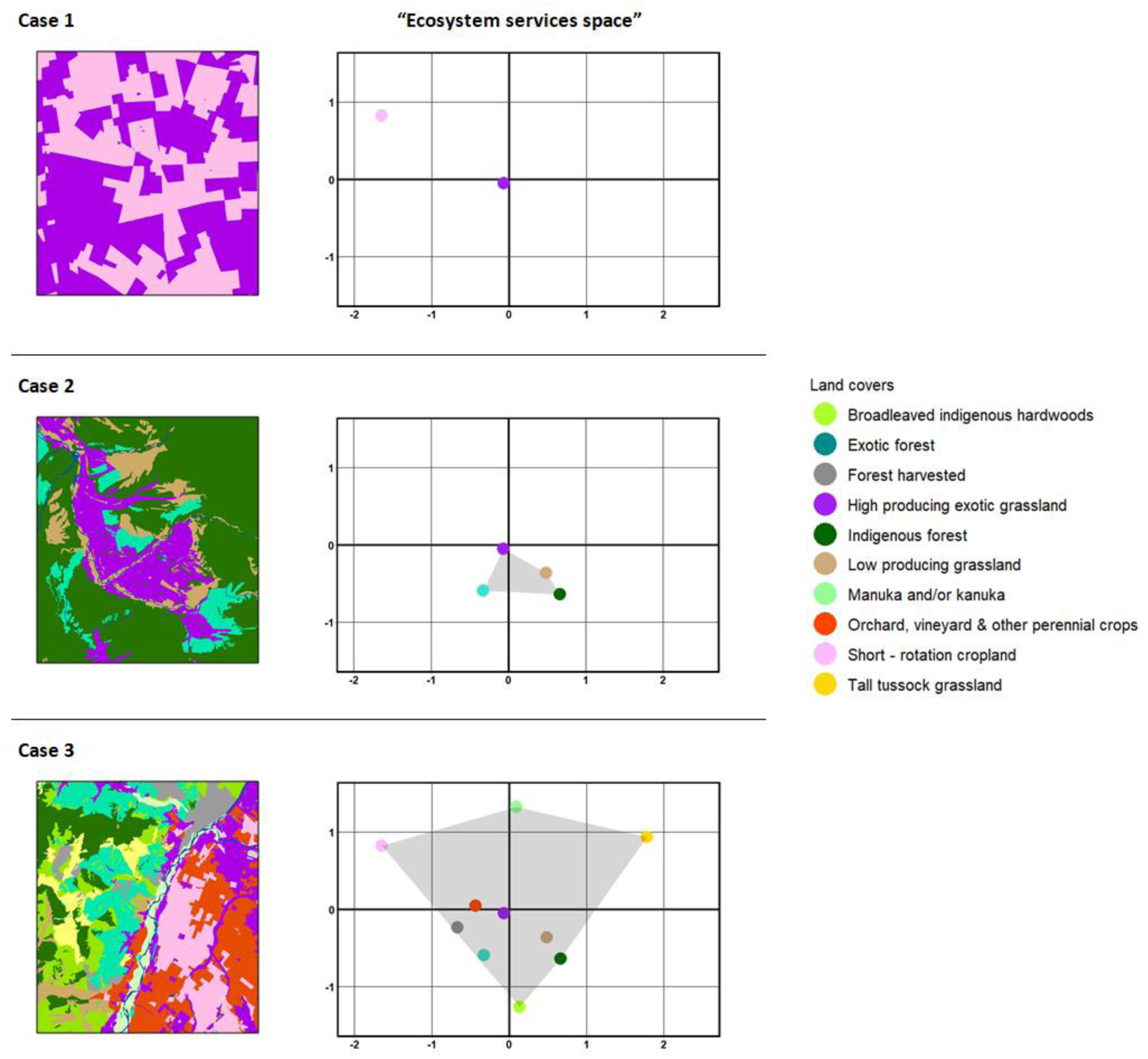
Example visualizations for exploring land cover trade-offs in the supply of multiple ecosystem services (ES) from entire landscapes. Quantitative measures of ES supply by different land uses or land covers (such as those obtained from our meta-analysis) can be used to generate ordinations that ‘map’ land covers or land uses into the multidimensional space of ES supply (ordination graphs). Distribution of land covers within that space can assist with identification of redundancies in ES supply (among land covers/uses that map close together) and trade-offs among land covers/uses that supply contrasting sets of ES and, consequently, occupy opposite extremes of the ordination space. Furthermore, the hypervolume enclosed by the total set of land covers/uses from a given landscape expresses the diversity of ES provided by that landscape. As an example, our data can be used to compare multi-service provision for: a landscape with few, undifferentiated production land covers (Case 1); a landscape with a combination of some production and non-production land covers (Case 2) and a landscape with a broad range of production and non-production land covers that supply a diverse range of services (Case 3).

For example, Case 3 in Fig. 3, has the greatest diversity of land covers and thus occupies the greatest hyper-volume in multidimensional ES-supply space. However, there are few land covers at the edge of this volume, such that the full array of services has low redundancy compared with Case 2 where land covers cluster around one location in ES-supply space. Because the entire ES-supply space may include areas that do not correspond to any configuration of ES, this approach is best applied for comparing landscapes rather than as an absolute measure of ES in one location.

Finally, mapping ES in multidimensional land-cover or land-use space (e.g., Fig. 1) allows the identification of ES bundles that respond similarly to land cover / land use. These bundles can then be used to identify management decisions that minimize disruption of service flows. Our approach opens the way for actively incorporating existing sources of information into ES research and informing practitioners to shape land systems that sustainably support human well-being.

## V. CONCLUSIONS

1. Our synthesis of land cover supply of ES in New Zealand revealed a consistent trade-off in the services supplied by high-value production land covers vs. those with low or no production and native elements in their vegetation cover. While production land covers specialized in the supply of primary production, low or no production land covers supplied a broad array of supporting and regulating ES. We did not find any evidence that forest cover was associated with any distinct patters of ES supply.
2. We show that the trade-off between ES supplied by production and non-production land covers is not solved with a single land cover. In contrast to earlier suggestions that a single natural ecosystem can support multiple ES at high levels (Foley *et al*., 2005), our analyses reveal that a mosaic of different land covers will be required to supply multiple ES within a landscape.
3. We show that exploring how different land covers map on to multidimensional ES space allows for an assessment of how diverse and resilient different combinations of land covers can be in their supply of ES. Such assessments can effectively support land use planning decisions beyond considerations of the specific identity of each land cover and the ES it supplies.
4. Our work suggests that there is great potential in using existing data for assessing ES bundles and interactions more cost-efficiently than through direct field observation. However, we also find that effective landscape management of ES will require further research on how environmental and land management factors can mediate the effects of land use on ES supply. We anticipate that these effects will differ across ES and will be more pronounced for ES where there is high variability in the supply by individual land covers (e.g., erosion control in our dataset).

## Supporting information

Supplementary Methods, Supplementary Results

## VI. ACKNOWLEDGEMENTS

We thank Melanie Hamzah for her assistance in the abstract screening and Sol Heber, Sophie Hunt, Jessica Furlong & Matthew Scott for their help with the full-text assessment and data extraction to collate our dataset. We also thank thematic experts Karen L. Adair (soil microbiology), Catherine M. Febria (freshwater ecology), Leo Condron (soil biogeochemistry), Matthew Turnbull (plant physiological ecology) & Angus McIntosh (freshwater ecology) for help with interpreting potential ecosystem service indicators. We are grateful to Daniel Stouffer and Melissa Ann Broussard for their valuable technical advice and members of the Stouffer and Tylianakis lab for their comments and useful discussions. This work was funded by the Ministry of Business, Innovation and Employment, NZ programme BEST: Building biodiversity into an ecosystem service-based approach for resource management (C09×1307). We thank Suzie Greenhalgh for coordinating this programme and providing helpful discussions.

## VII. AUTHOR CONTRIBUTIONS

JMT, EGB and CGC conceived and developed the original project concept; JMT and EGB secured the funding for the project; all authors contributed to the study design; data was acquired by CGC, with input from JMT and EGB; CGC, JMT, SN and ML contributed to the data analysis; CGC drafted the manuscript; JMT, SN, ML and EGB made revisions and critical appraisals of the manuscript.

*References marked with asterisk have been cited within the supporting information*.

## IX. SUPPORTING INFORMATION

Additional supporting information may be found online inthe Supporting Information section at the end of the article.

**Supplementary Methods 1.** Detailed data collection and processing methods.

**Supplementary Methods 2.** Full search phrase for pilot and formal searches.

**Supplementary Methods 3.** Decision tree for full-text assessment.

**Supplementary Methods 4.** Conversion of confidence intervals to variance and imputation of missing values.

**Supplementary Methods 5.** Scree plots and land cover classification for multivariate analyses.

**Supplementary Results 1.** Overview of research effort for New Zealand.

**Supplementary Results 2.** Evidence base and network meta-analysis for individual ES.

**Supplementary Results 3.** Summary of log response ratios per land cover and ecosystem service combination.

**Supplementary Results 4.** Detailed results from PERMANOVA analyses.

**Supplementary Results 5.** Data analysis with allocation of a single ES to each indicator.

**Supplementary Dataset 1**. Overview of Ecosystem Services (ES).

**Supplementary Dataset 2.** Overview of land cover classes as defined in New Zealand’s Land Cover Database (LCDB).

**Supplementary Dataset 3.** Quantitative indicators used to quantify supply of each ecosystem service.

**Supplementary Dataset 4.** Reference list for the studies included in our meta-analysis.

**Supplementary Dataset 5** Final log Response Ratios on ecosystem service supply for pairwise comparison of land covers in each study used in our analysis.

