## Supplementary Methods, Supplementary Results for "Consistent trade-offs in ecosystem services between land covers with different production intensities"

#### Table of Contents

**A. Supplementary Methods** 3

*Supplementary Methods 1. Detailed data collection and processing methods* 3
- Literature search 3
- Document screening and assessment 3
- Data aggregation and calculation of effect sizes 5

*Supplementary Methods 2. Full search phrase for pilot and formal searches* 8
- Phrase used in the formal search 8
- Phrases used in the pilot search 10

*Supplementary Methods 3. Decision tree for full-text assessment* 12

*Supplementary Methods 4. Conversion of confidence intervals to variance and imputation of missing values* 13
- Conversion of 95% confidence intervals to variance 13
- Regression model for applying Taylor's Law 13

*Supplementary Methods 5. Scree plots and land cover classification for multivariate analyses* 14

**B. Supplementary Results** 16

*Supplementary Results 1. Overview of research effort for New Zealand* 16

*Supplementary Results 2. Evidence base and network meta-analysis for individual ES* 19
- Regulation of water timing and flows 19
- Nutrient cycling 21
- Habitat provision 25
- Soil formation 26
- Freshwater provision 29
- Water purification 31
- Global climate regulation 33
- Primary production 34
- Water cycling 36
- Erosion control 39
- Pest regulation 40
- Waste treatment 44
- Capture fisheries 45
- Ethical and spiritual values 47
- Disease mitigation 48
- Pollination 50
- Regional & local climate regulation 52
- A note on confidence intervals and the size of the evidence base 53

*Supplementary Results 3. Summary of log response ratios per land cover and ecosystem service combination.* *55*

*Supplementary Results 4. Detailed results from PERMANOVA analyses* *56*

*Supplementary Results 5. Data analysis with allocation of a single ES to each indicator* *57*

**Captions for datasets S1 to S5** **65**

**Supplementary references** **66**

### A. Supplementary Methods

#### Supplementary Methods 1. Detailed data collection and processing methods

##### Literature search

To assess the viability of our project, an initial scoping search was conducted using Google Scholar and some general terms to capture our review subject (essentially, "New Zealand", "land use" or variations of this, and keywords on some of the ecosystem services, see Supplementary Methods 2). After manually screening the results of the scoping search, 201 potentially relevant documents were identified, suggesting that the project would not be limited by insufficient evidence/data.

The multidisciplinary database Scopus was selected for our formal search since it provided uniform access (i.e. independent of institutional subscription categories) to a comprehensive collection of abstracts and citations from international, peer-reviewed journals and serial books. We searched for titles, abstracts and keywords that contained at least one match in each of the 3 components that structured our search: 1) "New Zealand", 2) land cover and land use terms, and 3) ecosystem service terms (see Supplementary Methods 2 for the full search phrase). Land cover terms included all possible variations of "land use" and "land cover" as well as terms on specific land covers (both generic and specific to New Zealand. The ecosystem services component drew upon the names of each service (and possible variations of these) but also included vocabulary describing processes and conditions that could reflect their supply at the site scale akin to an individual land cover unit. To distill the final set of land-use and ecosystem service terms, 69 trial searches were conducted. This ensured that the final phrase, with approximately 840 terms, was sufficiently comprehensive. The search was finalized in December 2014, and was constrained to include documents published from 1970 onward, to be comparable with current land use regimes in New Zealand (MacLeod & Moller, 2006).

##### Document screening and assessment

In total, 9,741 citations matched our search criteria. The titles, abstracts and keywords of these citations were subjected to an automatic filtering that removed any duplicates and selected only those that mentioned at least two different land cover terms. We conducted the latter step because our search returned studies with at least one land cover term whereas we required studies that compared two or more land covers. The abstracts of the 4,373 citations marked as relevant after this filtering were then screened to check whether they pertained to research that could potentially allow for the quantitative comparison of two or more different terrestrial land covers, in New Zealand, for the supply of any of the 35 ecosystem services we defined for the project.

Abstracts were screened using an interactive machine learning system for semi-automated abstract screening, Abstrackr, which is often used in medical meta-analyses (Wallace *et al.*, 2012). By using an active learning approach and a dual supervision classification algorithm, Abstrackr draws from the selection decisions and relevant/irrelevant words reviewers find in a sample of their abstracts to estimate the likelihood of the unscreened abstracts being relevant. The screening order of the remaining abstracts is subsequently prioritized according to this likelihood. Abstracts in our study were screened by two reviewers who, after checking for agreement in their decisions for a common pilot set of 500 abstracts, independently reviewed the remaining abstracts until a stopping point of 50 consecutive non-relevant citations was reached. This stopping point was reached after 2,957 abstracts were screened, leaving 1,416 unscreened ones, which the machine-learning algorithm deemed to be less relevant than the 50 that comprised our stopping point.

The abstract screening and the initial pilot search yielded 914 relevant abstracts, which were passed on to a team of 4 reviewers for full-text assessment and data extraction. The full-text assessment included the following inclusion criteria:

1. Selected studies had to present quantitative data derived from original research conducted in 1970 or later, and which did not duplicate data that had been published in other studies already in our analysis.

2. Only quantitative measures that could be taken as indicators of the supply of one (or more) ecosystem services (according to the service definitions in Supplementary Dataset 2) were extracted from each study.

3. The data for each land cover had to come from at least two replicate observations. For any given study, a replicate observation was defined as one taken from a land cover unit that could be identified as a distinct spatial feature and which had sufficient separation from other units of the same land cover included in the same study so as to ascertain spatial independence of the observations. For the cases where the spatial separation of two units of the same land cover could not be readily ascertained, we applied a distance criterion. Instead of defining a fixed minimum distance between replicate land cover units (which could vary depending on the scale of a service), we defined as separate replicates any two units of the same land cover that were separated such that the distance between them was larger than the distance between any of them and any neighboring units of a different land cover.

Given the diversity of ecosystem services in our review, the documents we assessed comprised a very heterogeneous set of sampling and experimental designs, which meant that a land cover unit could range in size from whole forests and catchments to forest fragments, fields and crop plots; all of which were included as long as we could verify that the unit was spatially independent from other units of the same land cover and dominated in at least 80% by the same land cover type. Studies that did not have replicated observations (as defined above) for any land covers were discarded, whereas studies that contained replication on some, but not all, of the land covers were kept and only data on the replicated land covers were extracted. If, in the original studies, data were corrected for the potential effect of confounding covariates (i.e. slope), the corrected values were extracted instead of the uncorrected ones. Finally, within our review, the units of interest comprised only terrestrial land covers such that any comparisons between water bodies and any terrestrial land covers were not extracted. However, information on how different terrestrial land covers affected ecosystem services linked to a water body was included in our analysis. Full details of how the full-text selection criteria were applied can be found in Supplementary Methods 3.

Weekly meetings of the reviewers were held throughout the assessment and data extraction processes to ensure consistent implementation of these criteria. Authors were contacted whenever the data presented in a study were not readily extractable (due to legibility or formatting issues) or when further clarification was needed on the types of land covers involved in the study or the methods used to sample them. Authors of 96 studies were contacted with a 50% success response rate. In total, data from 260 studies were extracted.

#### Data aggregation and calculation of effect sizes

Extracted data from all studies were recorded in a database. As described in the main text, we assigned the quantitative measures of ecosystem service supply (indicators) reported by each study to one or more ecosystem services (Supplementary Dataset 1) and defined whether service supply would generally increase or decrease with larger values of the indicator. Thematic experts (see Acknowledgements) were consulted when there was uncertainty in the allocation of an indicator to a service or the direction of the relation between them.

Our database included some cases where either a single document contained the results of multiple experiments (each with a unique method or indicator) or, conversely, different results of a single experiment were published in separate documents. The latter case included studies with partial duplication of the results in different publications and a case where the results for different land covers for the same experiment were published in separate documents. To bring these studies into comparable terms with those that had the results of experiments published in only one article, we generated new unique study identifiers such that, for all effects in our review, these cases would either be treated as separate studies (i.e. when a single publication presented the outcomes of multiple, separate experiments) or merged into a single study (i.e. where the outcomes of a single experiment was published in multiple studies with no or partial overlap). For cases where two or more publications contained duplicated results from a single experiment, only data from the publication with the most comprehensive set of results was kept in our dataset.

In addition, several studies in our database contained multiple or repeated measures of the same indicator within a single land cover replicate. To allow for a standardized comparison across all studies, these were summarized to a single value per land cover replicate. In cases where the multiple measures were taken at different soil or water depths, the measurement of the topmost layer that occurred across all the land covers in the study was taken as the summarized value. For all other cases, repeated or multiple measures were summarized into a single mean value of the service indicator per replicate. This is equivalent to aggregating data to one response value per individual patient in medical meta-analyses. Studies that did not provide enough information to allow for the aggregation of multiple/repeated measurements to the standard replication level had to be excluded from the analysis.

Some studies reported a summary for all replicates of the same land cover; for the remaining studies, the mean and variance across replicates of the same land cover (and ecosystem service indicator) were calculated. For studies where data were already summarized across land cover replicates, but were presented as medians with either absolute and/or interquartile ranges, conversions to means and variances were made following the methods defined by Wan and colleagues(2014). Similarly, conversions from standard deviation, standard error and 95% confidence intervals to variance were also applied for summarized data that presented these measures of variation (see Supplementary Methods 6 for the equations to convert 95% confidence intervals). For studies that reported a mean value for the indicator per land cover, but no measure of variance, Taylor's power law (Taylor, 1961) was used to impute variances with estimates from a linear regression model of all reported means and variances in our dataset in log-log space (the full regression model can be found in Supplementary Methods 6). Imputed variances accounted for 11% of the records in the final dataset found in Supplementary Dataset 5. Cases where data could not be converted into means with variances across replicate land covers (including data on only the maximum and minimum values, geometric means with no variation and medians with standard errors or lacking variation or sample size) were not included in the dataset.

We adopted a log response ratio as the standard effect measure for comparing pairs of land covers within each study. For each pair of land covers (A and B) in a study, this measure was estimated as ln(A) - ln(B) for the indicators in which larger values corresponded to an increase in service supply. For the indicators in which larger values reflected a decrease in service supply (e.g., when the amount of soil eroded was a measure of erosion control) the inverse of the log response ratio was used i.e. - (ln(A) - ln(B)). For all cases the variance of the log response ratio ($v_{RR}$) was estimated as (Rosenberg, Rothstein, & Gurevitch, 2013):

$$v_{RR} = \frac{s_A^2}{n_1 \overline{Y}_A^2} + \frac{s_B^2}{n_2 \overline{Y}_B^2}$$

Where $\overline{Y}$ denotes the mean value of the ecosystem service indicator for land covers A and B, with variance $s$ and sampling size $n$. For cases where both land covers had negative numbers in the indicator, we based the ratio and variance calculations on the absolute indicator values and took the inverse of the ratio calculated with absolute values as the final value. Similarly, in cases where at least one of the land covers had a value of zero for the ecosystem service indicator, we added a small value (3 orders of magnitude smaller than the smallest value in the dataset) to the zero to allow for the ratio and variance calculations.

Studies that presented multiple indicators for a single ecosystem service from each site (e.g., measures of the cycling rates of different nutrients), or which presented data on the same indicator expressed in two different units, required further aggregation of their log response ratios to avoid giving these studies disproportionate weight in the analysis. For some studies, not all the indicators of a service (or different units for the same indicator) had information from all of the land covers for the study. Thus, for each study, we only aggregated the indicators of the same ecosystem service (or sets of data with different units on the same indicator) that were measured from all of the land covers present in the study for that service, and excluded from the analysis any indicators that were not presented for the full range of land covers. If only a single indicator (or unit) contained data on all of the land covers then the other indicators of that service were excluded for that study. If two or more indicators (or units) contained data for the full set of land covers, then data were aggregated by taking a mean of the log response ratios and corresponding variances across the different indicators for each pairwise combination of land covers of that service in that study. The final dataset with aggregated log response ratios for each study and ecosystem service can be found in Supplementary Dataset 5.

#### Supplementary Methods 2. Full search phrase for pilot and formal searches

##### Phrase used in the formal search

Database used: Scopus
*Please consult https://dev.elsevier.com/tips/ScopusSearchTips.htm for an explanation of Boolean operators and wildcards used*
Search was conducted on the 26th January, 2015.

Phrase:

TOPIC: (“New Zealand”)
AND
TOPIC: (“land use” OR “land cover” OR habitat OR “vegetation type” OR ecosystem* OR forest* OR plantation OR scrub* OR shrub* OR pasture* OR grass* OR crop* OR “tussock grassland” OR “grey scrub” OR bush OR herbfield OR catchment OR “drainage basin” OR watershed* OR wetland* OR river OR lake OR peatland OR marsh OR bog OR fernland OR flaxland OR matagouri OR mangrove OR orchard OR estuary OR urban OR mine OR town OR city OR residential OR park OR garden)
AND
TOPIC: (“ecosystem service*” OR “habitat quality” OR “habitat provision” OR “nursery provision” OR “habitat diversity” OR “habitat complexity” OR “habitat feature” OR “habitat character*” OR ((feeding OR resting OR roosting OR nesting OR brood OR foraging OR mating) NEAR (site* OR cover)) OR “vegetation cover” OR “food availability” OR nurser* OR nitrogen OR phosphorus OR potassium OR calcium OR magnesium OR sulphur OR nitrification OR “soil organic matter” OR nitrification OR fixation OR “nutrient cycl*” OR “chemical cycl*” OR “decomposition” OR “nutrient uptake” OR “nutrient export” OR detritus OR bacteria OR microorganism OR “biogeochemical cycl*” OR microbial OR decomposition OR “soil formation” OR weathering OR “humification” OR “mineralization” OR pedogen* OR “soil quality” OR “soil fertility” OR “soil nutrients” OR “nutrient leaching” OR “microbial biomass” OR “nutrient storage” OR “soil structure” OR “nutrient assimilation” OR biomass OR “primary production” OR “primary productivity” OR litter* accumulation OR aboveground OR belowground OR NPP OR “carbon allocation” OR “productivity allocation” OR “air quality” OR pollut* OR “nitrogen oxide*” OR “sulphur and oxide*” OR aerosol* OR “atmospheric cleansing capacity” OR “tropospheric oxidizing capacity” OR “acid rain” OR particulate OR “volatile organic compounds” OR “carbon stock” OR “total carbon” OR “total C” OR “carbon storage” OR emission* OR “carbon loss” OR ((carbon OR methane OR “tropospheric ozone” OR aerosol* OR “greenhouse gas*” OR “nitrous oxide”) SAME (emission* OR sink OR sequestration)) OR albedo OR “heat flux” OR evapotranspiration OR precipitation OR rainfall OR temperature OR wind OR humidity OR “climate regulation” OR “climatic variability” OR runoff OR interception OR infiltration OR “water flow” OR discharge OR “water retention” OR “lag time” OR “water storage” OR “aquifer recharge” OR “stream*flow” OR “water yield” OR “water balance” OR “base*flow” OR percolation OR “flow regime” OR “flow regulation” OR erosion OR gullying OR gully OR “soil cover” OR “vegetation cover” OR rill OR “soil loss” OR “sediment yield” OR “sediment retention” OR “soil stability” OR “soil compaction” OR “aquatic or pollution” OR “water quality” OR “water purification” OR “water filtration” OR “filtration” OR “dissolved organic carbon” OR “heavy

metals” OR “dissolved oxygen” OR “nutrient retention” OR “microbial degradation” OR “benthic indicators” OR “nutrient removal” OR “maximum daily loads” OR “load” OR (waste SAME (regulat* or treat* or assimilat* or decompos* or process* or degrad*)) OR pollut* OR toxic* OR contaminant* OR “detoxification” OR “soil pollut*” OR “nutrient retention” OR remineralisation OR “human AND pathogen*” OR " human AND disease*" OR “infectious disease*” OR (propagation SAME (disease OR vector OR pathogen)) OR “disease vector” OR “pathogen infect*” OR “disease risk” OR “disease incidence” OR “ecology of disease” OR “disease ecology” OR “vector control” OR “invasion resistance” OR “pest control” OR " pest management" OR biocontrol OR “biological control” OR “biological pest control” OR “natural pest control” OR weed* OR “pest predat*” OR (“natural enemy” SAME (conservation OR augmentation)) OR “seed set” OR pollinat* OR “flower visit*” OR zoophilus OR ornithophilous OR melittophilous OR entomophilous OR fruit OR “crop plants” OR “hazard mitigation” OR “disaster reduction” OR “disaster risk reduction” OR buffer* OR ((storm OR flood OR drought OR fire OR landslide OR avalanche OR “mass movement” OR hurricane OR windstorm) SAME (protect* OR buffer* OR mitigate* OR attenuat* OR defen*)) OR “flood storage” OR “storm flow” OR “peak flow” OR “extreme event*” OR “storm peak” OR ((timber OR round*wood OR pulp*wood OR wood) NEAR (harvest* OR yield OR extraction OR production)) OR “forest product” OR “Non-timber forest product” OR “non-wood forest product” OR ((fiber OR leather OR hemp OR hide OR “merino wool” OR yarn OR “alpaca wool” OR “merino wool” OR “possum wool” OR “possum fur” OR “harakeke flax” OR “flax fibre” OR wool OR fur) SAME (production OR supply OR provision OR yield OR extraction)) OR ((“fuel wood” OR “wood fuel” OR firewood) SAME (production OR extraction))OR biofuel OR “biomass energy” OR biogas OR biodiesel OR “woody biomass” OR “cultural identity” OR “maori” OR “livelihood” OR ((sacred OR spiritual) SAME (site OR landscape OR place OR plant OR animal* OR ecosystem)) OR “spiritual inspiration” OR “ritual site” OR “sacred grove” OR “sense of belonging” OR aesthetics OR “scenic value” OR “environmental attribute*” OR “site attribute*” OR “aesthetic enjoyment” OR “aesthetic preference” OR “environmental aesthetics” OR “scenic beauty” OR “aesthetic pleasure” OR “environmental perception” OR wilderness OR “landscape preference*” OR “visual landscape” OR hedonic OR “cultural importance” OR “archaeological site*” OR “historic site” OR “heritage site” OR “ancestral site” OR “cultural landscape” OR “cultural heritage” OR “cultural site” OR “cultural attribute* OR”traditional landscape" OR landmark* OR “ritual site” OR “burial site” OR “tribal landmark” OR “natural heritage place*” OR “Maori site*” OR “intellectual development” OR “traditional knowledge system” OR ethnobotan* OR ethnobiolog* OR “maatauranga maori” OR “experimental farm” OR “educational forest” OR “educational farm” OR “distribution research project” OR “distribution research locations” OR “distribution research site*” OR “didactic farm” OR “didactic forest” OR “educational visit*” OR “school visit*” OR “field trip*” OR “field station” OR “research site*” OR “research location*” OR “research project location*” OR inspirational site OR ((movie OR film OR photograph* OR painting) SAME (setting OR location)) OR “maori art” OR craft* OR “inspiration from nature” OR “nature in art” OR “nature in film” OR “nature in literature” OR biomimicry OR bionics OR biomimet* OR recreation OR visit* OR tourism OR “nature tourism” OR ecotourism OR “adventure tourism” OR “rural tourism” OR “agri*tourism” OR “cultural tourism” OR “nature*based tourism” OR “nature*based recreation” OR angling OR hiking OR tramping OR birding OR hunting OR fishing OR mountaineering OR alpinism OR walking OR kayaking OR rowing OR surfing OR sailing OR rafting OR canoeing OR skiing OR “snow sport*” OR “winter sport*” OR windsurfing OR “kites surfing” OR horse riding" OR caving OR “outdoor sport*” OR rappelling OR abseiling)

##### Phrases used in the pilot search

Database used: Google Scholar
Searches were conducted between February - April, 2014.

1. “New Zealand” AND “land use” AND diversity
2. “New Zealand” AND “land use” AND biodiversity
3. “New Zealand” AND “land use” AND H’
4. “New Zealand” AND “land use” AND evenness
5. “New Zealand” AND “land use” AND “species richness”
6. “New Zealand” AND “land use” AND “species abundance”
7. “New Zealand” AND “land use” AND insect
8. “New Zealand” AND “land use” AND arthropod
9. “New Zealand” AND “land use” AND invertebrate
10. “New Zealand” AND “land use” AND Collembola
11. “New Zealand” AND “land use” AND Diptera
12. “New Zealand” AND “land use” AND beetle
13. “New Zealand” AND “land use” AND Coleoptera
14. “New Zealand” AND “land use” AND arachnid
15. “New Zealand” AND “land use” AND spider
16. “New Zealand” AND “land use” AND mite
17. “New Zealand” AND “land use” AND Acari
18. “New Zealand” AND “land use” AND bird
19. “New Zealand” AND “land use” AND avifauna
20. “New Zealand” AND “land use” AND plant
21. “New Zealand” AND “land use” AND vegetation
22. “New Zealand” AND “land use” AND aquatic
23. “New Zealand” AND “land use” AND stream
24. “New Zealand” AND “land use” AND “water yield”
25. “New Zealand” AND “land use” AND “water quality”
26. “New Zealand” AND “land use” AND soil
27. “New Zealand” AND “land use” AND “microbial biomass”
28. “New Zealand” AND “land use” AND microbes
29. “New Zealand” AND “land use” AND biota
30. “New Zealand” AND “land use” AND fungi
31. “New Zealand” AND “land use” AND mycorrhiza
32. “New Zealand” AND “land use” AND nutrient
33. “New Zealand” AND “land use” AND nitrogen

34. “New Zealand” AND “land use” AND phosphorus
35. “New Zealand” AND “land use” AND potassium
36. “New Zealand” AND “land use” AND carbon
37. “New Zealand” AND “land use” AND methane
38. “New Zealand” AND “land use” AND transpiration
39. “New Zealand” AND “land use” AND evapotranspiration
40. “New Zealand” AND “land use” AND photosynthesis

Searches also substituted “land use” with:

1. “catchment”
2. “paired catchment”
3. “vegetation type”
4. “site” and “comparison”
5. “forest” and “tussock”
6. “forest” and native bush"
7. “forest” and “pasture”
8. “plantation” and “native”
9. “plantation” and “pasture”
10. “plantation” and “tussock”
11. “native and pasture”
12. “native and tussock”
13. “tussock” and “bush”
14. “tussock” and “pasture”
15. “paired catchment” and “forest”
16. “paired catchment” and “tussock”
17. “paired catchment” and “pasture”
18. “paired catchment” and “bush”
19. “paired catchment” and “native”
20. Pinus
21. Podocarp
22. Broadlea*
23. Chionochloa
24. Nothofagus
25. Pasture
26. Hieracium
27. Scrub
28. Shrubland
29. “Grey shrub”

### Supplementary Methods 3. Decision tree for full-text assessment

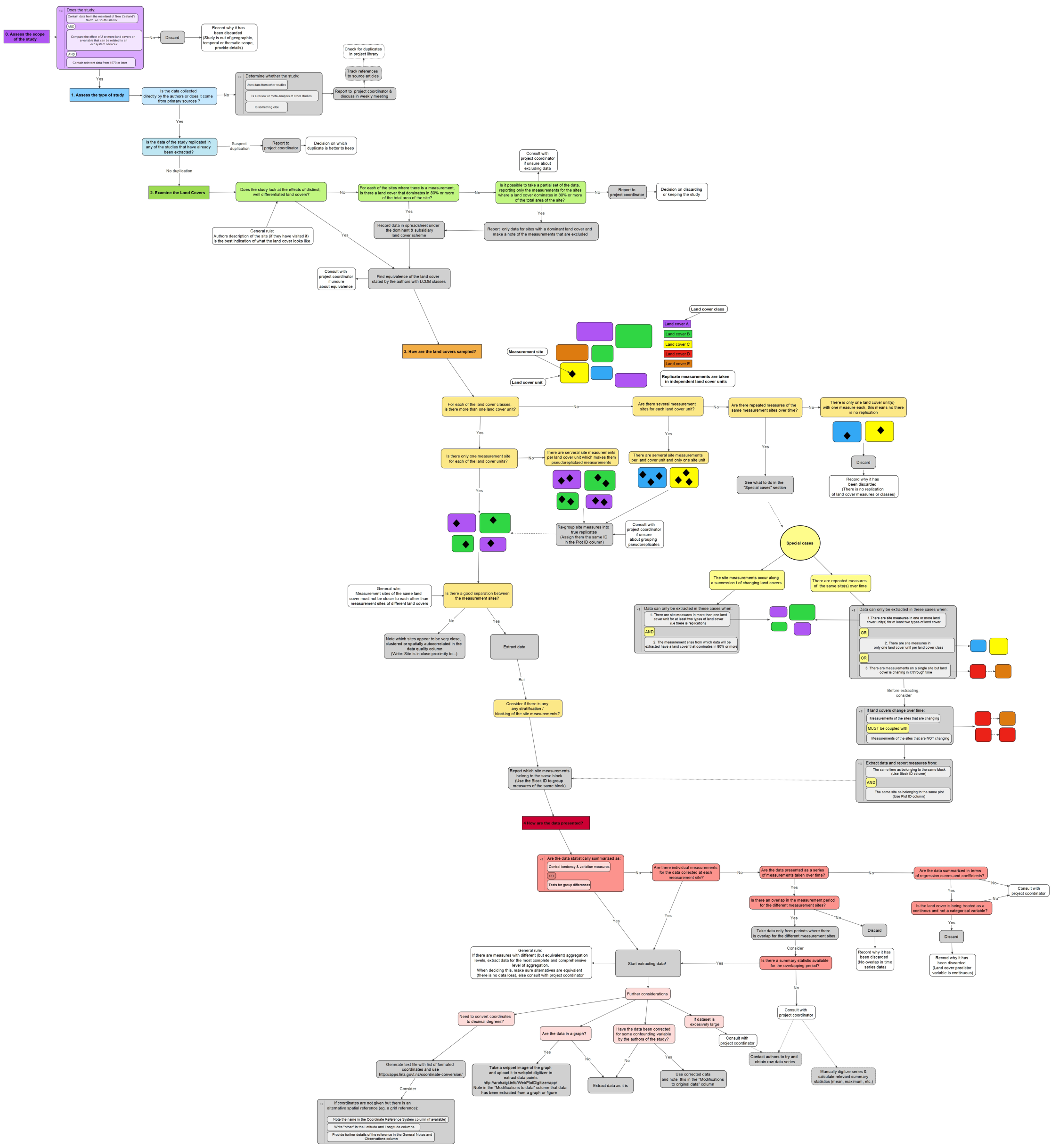

*Figure S1: Decision tree with the selection criteria used in the full-text assessment of publications with relevant abstracts for our review. Note: this figure is intended to be viewed in PDF format at magnification*

#### Supplementary Methods 4. Conversion of confidence intervals to variance and imputation of missing values

##### Conversion of 95% confidence intervals to variance

$$s^2 = \frac{95\%CI}{t - critical} \times \sqrt{n}$$

Where $s^2$ denotes the variance, 95%CI corresponds to the 95% confidence interval and $n$ is the sample size reported by the authors. The $t$-critical value is the value in the $t$-distribution for the corresponding alpha and degrees of freedom. For all studies in our dataset where we needed to apply this conversion, the degrees of freedom were not available so we approximated the $t$-critical value as two. This was done because the two-sided $t$-distribution values for an alpha of 0.05 start at 2.57 and asymptote at 1.96 for 5 or more degrees of freedom.

##### Regression model for applying Taylor's Law

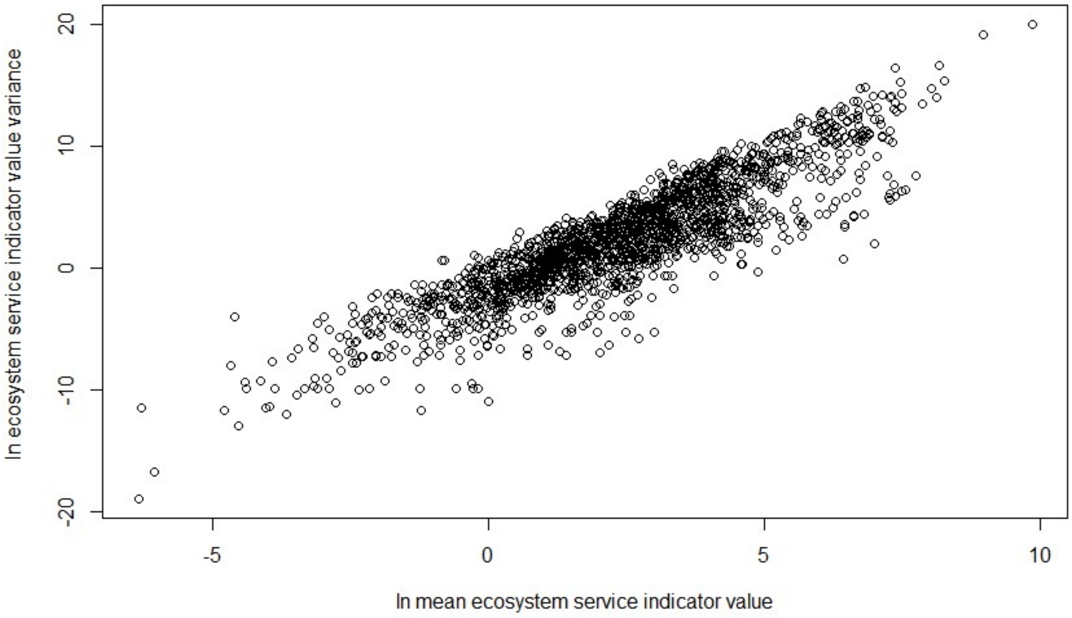


*Figure S2: Regression (in natural logarithm space) of the mean ecosystem service indicator values for all land covers reported in our dataset against their corresponding variances.*

The equation used to impute variances from means based on regression coefficients from their relation in natural logarithm space is shown below. Coefficients are given to three decimal places in the equation, however their full values were used in the actual calculation of the imputed values.

$$s_{imp}^2 = e^{[-2.147+(1.878\times\ln(|x|)]}$$

Where $s_{imp}^2$ is the imputed variance and $x$ the mean used for the imputation.

#### Supplementary Methods 5. Scree plots and land cover classification for multivariate analyses

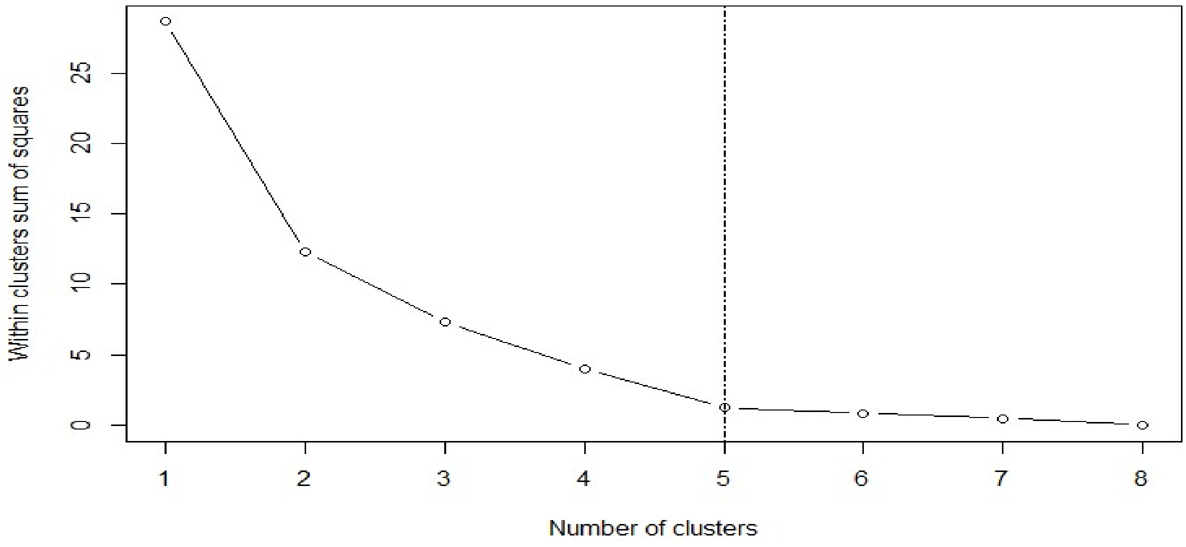


*Figure S3: Scree plot used to identify number of clusters for k-means cluster analysis of ES. Vertical dashed line shows the number of groups that minimizes within groups sum of squares and that was selected for the analysis.*

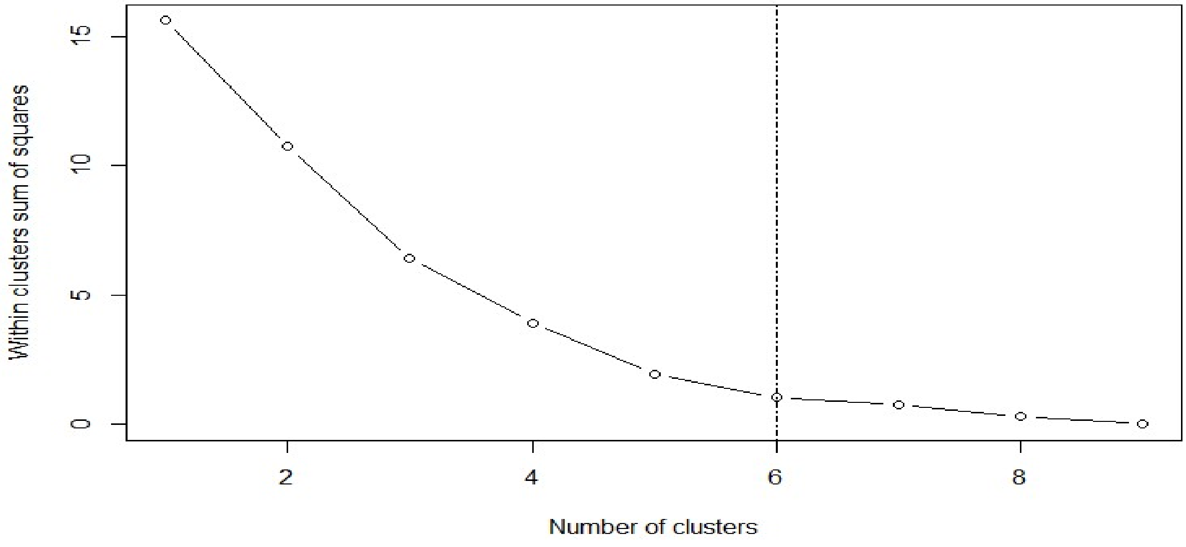


*Figure S4: Scree plot used to identify number of clusters for k-means cluster analysis of land covers. Vertical dashed line shows the number of groups that minimizes within groups sum of squares and that was selected for the analysis.*

Table S1: Delimitation of categorical variables used in PERMANOVA of land cover effects across ecosystem services

| Land cover | Forest Cover | Production | Type of vegetation cover |
|---|---|---|---|
| Indigenous forest | Present | No | Native |
| Exotic forest | Present | Yes | Exotic |
| Forest harvested | Present | Yes | Exotic |
| Manuka and/or kanuka | Present* | No | Native |
| Tall tussock grassland | Absent | No | Native |
| Low producing grassland | Absent | No** | Mixed |
| High producing exotic grassland | Absent | Yes | Exotic |
| Orchard vineyard & other perennial crops | Absent | Yes | Exotic |
| Short-rotation cropland | Absent | Yes | Exotic |

* Manuka and/or kanuka is a successional scrub often found on previously forested land (Thompson *et al.*, 2003). This land cover can be found as dense scrub or as forest and therefore has been classified as having a forest cover.
** Low producing grassland comprises a mix of native and exotic grasslands. It has been classified in the no production class because it has poor pastoral quality and extensive grazing or non-agricultural use (Thompson *et al.*, 2003).

Vegetation cover was not included in the analysis since all but one of the land cover groups in the native / exotic groups overlap with those in the production / non-production groups.

### B. Supplementary Results

#### Supplementary Results 1. Overview of research effort for New Zealand

Figure S5 shows that all of the supporting ecosystem services are represented in our database, whereas for the remaining categories our data only offer partial coverage of their corresponding services, with information on: nine out of the 11 regulating services, two of the 15 provisioning services and one of the three cultural services defined for the project in Table S1. For the categories that were represented by more than one ecosystem service in our database, the number of studies per service ranged from two to 45 for the regulating services; from five to 40 for the provisioning ones and from 21 to 57 in the supporting services. A total of four studies provide evidence for the single service in the cultural category in our database.

| Land cover | Habitat provision | Nutrient cycling | Soil formation | Primary production | Water cycling | Capture fisheries | Freshwater provision | Global climate regulation | Regional and local climate regulation | Regulation of water timing and flows | Erosion control | Water purification | Waste treatment | Disease mitigation | Pest regulation | Pollination | Ethical and spiritual values | Total |
|---|---|---|---|---|---|---|---|---|---|---|---|---|---|---|---|---|---|---|
| Indigenous forest | 31 | 20 | 11 | 19 | 11 | 3 | 24 | 7 | 1 | 23 | 12 | 21 | 2 | 4 | 5 | 0 | 2 | 55 |
| Deciduous hardwoods | 2 | 1 | 0 | 1 | 0 | 1 | 1 | 0 | 0 | 0 | 0 | 1 | 0 | 0 | 0 | 0 | 1 | 2 |
| Exotic forest | 24 | 25 | 28 | 13 | 11 | 2 | 18 | 21 | 2 | 26 | 6 | 17 | 4 | 2 | 2 | 0 | 2 | 59 |
| Forest harvested | 6 | 3 | 2 | 3 | 3 | 0 | 5 | 1 | 0 | 6 | 4 | 5 | 0 | 0 | 2 | 0 | 0 | 11 |
| Mixed exotic shrubland | 1 | 0 | 0 | 0 | 0 | 0 | 0 | 0 | 0 | 0 | 0 | 0 | 0 | 0 | 1 | 0 | 0 | 1 |
| Sub-alpine shrubland | 1 | 1 | 0 | 1 | 0 | 0 | 0 | 0 | 0 | 1 | 0 | 0 | 0 | 0 | 1 | 0 | 0 | 3 |
| Broadleaved indigenous hardwoods | 4 | 3 | 2 | 4 | 0 | 0 | 2 | 0 | 0 | 1 | 0 | 2 | 0 | 0 | 1 | 1 | 0 | 8 |
| Matagouri or grey scrub | 1 | 1 | 1 | 0 | 0 | 0 | 0 | 1 | 0 | 0 | 0 | 0 | 1 | 0 | 1 | 0 | 0 | 3 |
| Manuka and/or kanuka | 2 | 4 | 5 | 2 | 0 | 0 | 1 | 3 | 0 | 1 | 2 | 1 | 1 | 0 | 2 | 0 | 0 | 10 |
| Gorse and/or broom | 0 | 1 | 1 | 0 | 0 | 0 | 0 | 1 | 0 | 0 | 0 | 0 | 0 | 0 | 1 | 0 | 0 | 2 |
| Flaxland | 0 | 0 | 0 | 0 | 0 | 0 | 0 | 0 | 0 | 0 | 0 | 0 | 0 | 0 | 0 | 1 | 0 | 1 |
| Herbaceous saline vegetation | 0 | 0 | 1 | 0 | 0 | 0 | 0 | 0 | 0 | 0 | 0 | 0 | 0 | 0 | 0 | 0 | 0 | 1 |
| Herbaceous freshwater vegetation | 0 | 0 | 1 | 0 | 0 | 0 | 0 | 0 | 0 | 0 | 0 | 0 | 0 | 0 | 0 | 0 | 0 | 1 |
| Depleted grassland | 1 | 0 | 0 | 0 | 0 | 0 | 0 | 0 | 0 | 0 | 0 | 0 | 0 | 0 | 1 | 0 | 0 | 2 |
| Tall tussock grassland | 9 | 11 | 6 | 9 | 3 | 0 | 10 | 1 | 0 | 5 | 4 | 10 | 2 | 0 | 2 | 0 | 0 | 19 |
| Low producing grassland | 7 | 4 | 9 | 3 | 2 | 2 | 4 | 3 | 0 | 5 | 2 | 4 | 1 | 0 | 1 | 0 | 2 | 18 |
| High producing exotic grassland | 38 | 43 | 37 | 22 | 17 | 5 | 29 | 29 | 2 | 34 | 15 | 28 | 4 | 4 | 2 | 2 | 4 | 90 |
| Orchard vineyard & other perennial crops | 2 | 2 | 3 | 1 | 1 | 0 | 2 | 3 | 1 | 1 | 0 | 1 | 0 | 0 | 0 | 1 | 0 | 5 |
| Short rotation cropland | 6 | 16 | 18 | 2 | 2 | 0 | 4 | 12 | 0 | 9 | 4 | 6 | 0 | 0 | 1 | 1 | 0 | 24 |
| Alpine grass/herbfield | 0 | 0 | 1 | 0 | 0 | 0 | 0 | 0 | 0 | 0 | 1 | 0 | 0 | 0 | 0 | 0 | 0 | 1 |
| Gravel and rock | 0 | 0 | 0 | 0 | 1 | 0 | 0 | 0 | 0 | 1 | 0 | 0 | 0 | 0 | 0 | 0 | 0 | 1 |
| Transport infrastructure | 0 | 0 | 0 | 0 | 0 | 0 | 0 | 0 | 0 | 0 | 0 | 0 | 0 | 0 | 1 | 0 | 0 | 1 |
| Surface mine & dump | 0 | 1 | 0 | 0 | 0 | 0 | 0 | 0 | 0 | 0 | 0 | 0 | 0 | 0 | 0 | 0 | 0 | 1 |
| Urban parkland/open space | 0 | 0 | 0 | 0 | 0 | 0 | 0 | 0 | 0 | 0 | 0 | 0 | 0 | 0 | 2 | 2 | 0 | 4 |
| Built-up area (settlement) | 1 | 1 | 0 | 0 | 1 | 0 | 2 | 0 | 0 | 1 | 0 | 1 | 0 | 1 | 1 | 0 | 0 | 3 |
| Total | 52 | 57 | 51 | 32 | 21 | 5 | 40 | 33 | 2 | 45 | 22 | 40 | 6 | 4 | 12 | 3 | 4 | 133 |

*Figure S5: Distribution of studies per ecosystem service and land cover. For most services, data are concentrated along a selection of land covers: high producing exotic grassland (with a total of 92 studies across all ecosystem services), exotic forest (64 studies in total), indigenous forest (58 studies), short-rotation cropland (24 studies) and tall tussock grassland (20 studies). In addition, eight of the 43 land cover classes in the LCDB classification were not present in our data base. These land cover classes were: "sand, gravel and rock" (i.e. the coastal strip separating land from sea), "mangrove", "fernland", "landslide", "permanent snow and ice", "lake or pond", "river" and "estuarine open water". The last three units correspond to aquatic land covers which were not included in our review, whereas the remainder were simply poorly represented within the literature used for our review. Note that the total row and column do not match the actual sum of column and row counts because our dataset includes studies with data on multiple ecosystems and land covers. Likewise, the row and column totals do not add up to the grand total in the lower right corner which, instead, corresponds to the total number of studies in our dataset.*

Since we aggregated data from studies with multiple indicators of the same service, the matrix in Fig. S5 effectively reflects the number of data points in our spreadsheet for each ecosystem service and land cover combination. The actual number of indicators for each ecosystem service - land cover combination are shown in Fig. S6 which indicates that, overall, the number of indicators follow a similar distribution to that of the number of studies. However, for the most common ecosystem service-land cover combinations in our dataset (e.g., soil formation or nutrient cycling in both exotic forest and high producing exotic grasslands) there were as many as four to five times more indicators than studies, suggesting that studies with multiple indicators were more frequent in the land covers and ES that were also more commonly studied.

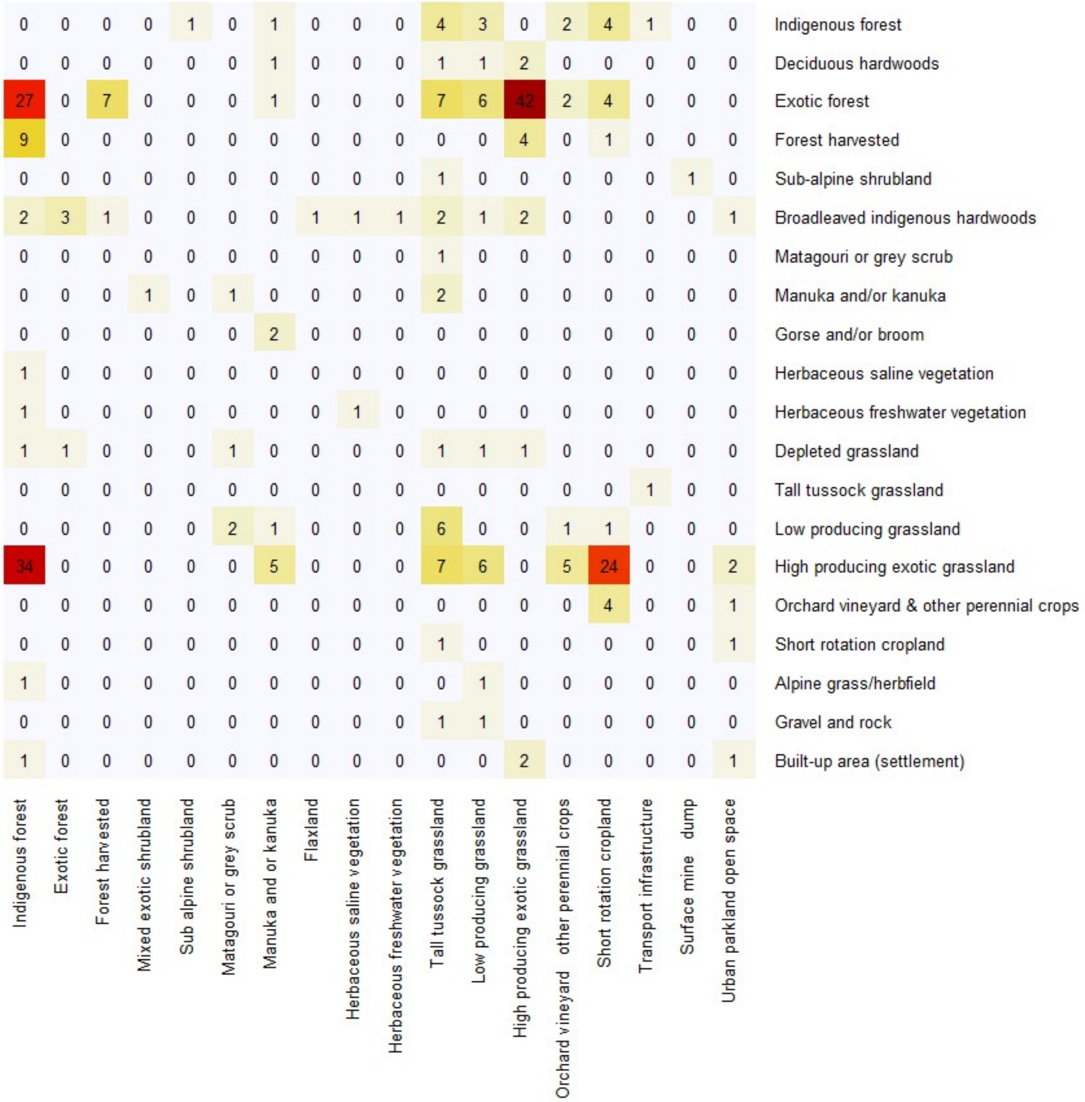


*Figure S6: Distribution of studies per land cover comparisons. Studies contributing data on multiple ecosystem services are only counted once in each pair of land covers where they contribute data.*

#### Supplementary Results 2. Evidence base and network meta-analysis for individual ES

Below we present the results of the individual network meta-analyses for each of the 17 ES in our dataset. For each ES we summarize details on the evidence base used in the analysis (including the types of indicators used, the number of studies and the network of land cover comparisons) and present the outputs of the analysis. The latter include a forest plot or blobbogram (sensu Lewis & Clarke, 2001) and two measures of heterogeneity: $\tau^2$ and $I^2$. In network meta-analysis, $\tau^2$ expresses the between-study variance, while $I^2$ expresses the extent of heterogeneity across the entire evidence network ($Q_{total}$) adjusted by degrees of freedom (which are defined by the number of treatment arms, i.e. land covers compared in each study – see Schwarzer, Carpenter, & Rücker, 2015 for further details). $I^2$ is expressed as a percentage with larger values indicating higher heterogeneity.

##### Regulation of water timing and flows

**Type of indicators for this ES:** Comparisons for this ES were drawn from 33 indicators. The main aspects quantified by these indicators pertain to soil characteristics that either supply greater regulation by enhancing soil water retention or have detrimental effects in the supply of this ES by promoting increased runoff. In addition, there are also some indicators on stream channel characteristics (such as its dimensions) that affect its ability to regulate water flow over time and that, to an extent, can be altered by land cover.

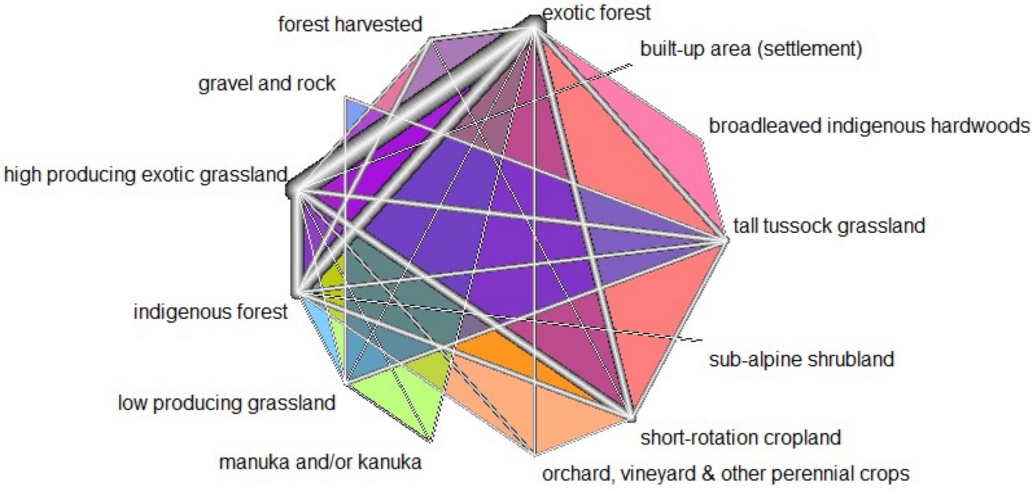


*Figure S7: Evidence network for land cover comparisons on regulation of water timing and flows. In all evidence network graphs, lines connect pairs of land covers that are compared in one or more studies and their thickness is inversely proportional to the standard error for the comparison, with thicker lines indicating smaller standard errors and, consequently, a greater evidence for the comparison. Shaded areas indicate the presence of multi-arm studies which compare three or more land covers.*

**Evidence base:** This evidence network is formed by 104 pairwise comparisons of 13 land covers. Data were obtained from 45 different studies, each contributing a minimum of one and a maximum of four pairwise land cover comparisons. As indicated by the thicker lines in Fig. S7, the land covers that are most commonly compared are:

- High producing exotic grassland (55 comparisons)
- Exotic forest (48 comparisons)
- Indigenous forest (41 comparisons)
- Short-rotation cropland (19 comparisons)

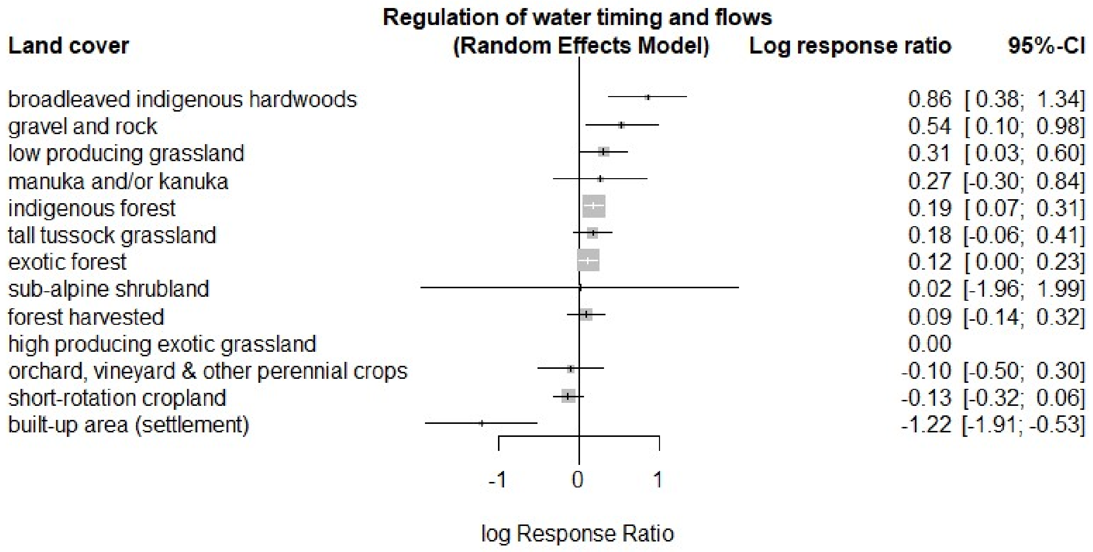


*Figure S8: Forest plot of land cover contrasts in the supply of regulation of water timing and flows. Random effects model using high producing exotic grassland as a reference. Log response ratios depicted here are the network meta-analysis model estimates of the overall ratios between each land cover and high producing exotic grassland. The model accounts for the direct and indirect comparisons in the evidence network, as well as the random effects from having comparisons on the same land covers drawn from different studies. Bars indicate the 95% confidence intervals for each estimate, while grey boxes reflect the relative weight of the comparison between each land cover and high producing exotic grassland in the overall model estimates. Comparisons that have greater weights are depicted with larger boxes. Land covers are presented in descending order of their P-scores which are calculated from the magnitude and precision of the log response ratio estimates for each land cover and provide a means to rank treatment effects (i.e. land covers) according to their comparative effectiveness (Rücker & Schwarzer, 2015).*

**Measures of heterogeneity/network inconsistency:**

$\tau^2$ = 0.195
$I^2$ = 65.573

**Main results:**

- Overall there is a gradient from native vegetation (broadleaved hardwoods and indigenous forest) to more artificial and production-oriented land covers (high producing exotic grassland, short-rotation cropland, built-up area).
- Exotic forest seems to perform similarly to indigenous forest, as do tall tussock grassland, manuka and/or kanuka, and low producing grassland.
- Built-up area stands out as the worst performing land cover in terms of regulation of water timing and flows, which is likely explained by the presence of impervious surfaces and channel morphologies that enhance runoff.
- The high infiltration capacity in gravel and rock probably accounts for its high ranking in the supply of this ES.

##### Nutrient cycling

**Type of indicators for this ES:** Comparisons for this ES were drawn from 143 indicators, most of which focus on the cycling and flow of nutrients within the soil system and characteristics of the soil environment that promote or hinder nutrient cycling. The latter were taken as negative indicators for the supply of nutrient cycling, as were the indicators on nutrient loss from the soil system. In addition, the data also include indicators on how land cover conditions plant uptake and the processing of nutrients both in the soil and freshwater systems. A large number of the indicators pertain to nitrogen and phosphorus, however, there is also information on other nutrients including: calcium, carbon, copper, magnesium, potassium, sulfur and zinc (we have followed the Millennium Ecosystem Assessment in their delimitation of the nutrients for this ES as those relevant for plant growth; Lavelle *et al.*, 2005).

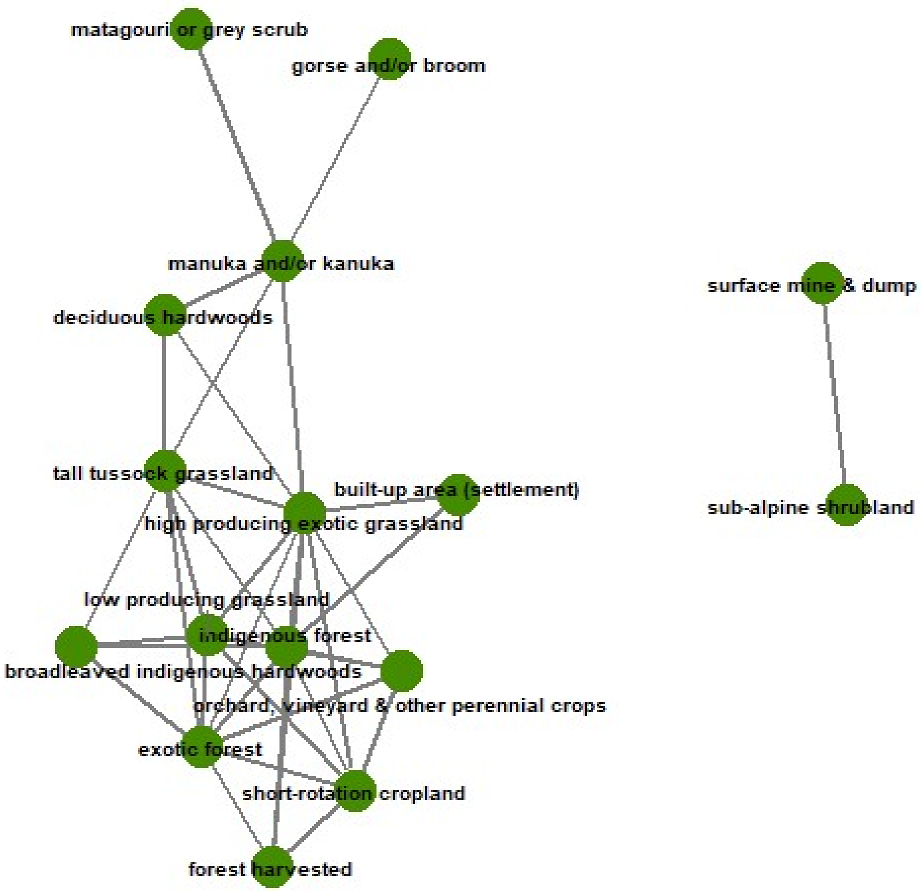


*Figure S9: Land-cover comparison networks for nutrient cycling. For this service, land cover comparisons were split into the two networks depicted here, however, only the comparisons in the larger network were used as evidence base for the network meta-analysis. Figure S10 presents the detailed configuration of this evidence base.*

**Evidence base:** The evidence base for this ES is split into two networks of land cover comparisons depicted in Fig. S9. The smaller of these networks holds the comparison between sub-alpine shrubland and surface mine & dump, for which there is only evidence from a single study and, consequently, is not connected to any of the land covers in the larger network. In the smaller network, the single study evidence defines a log response ratio of approximately 1.435 in favor of the sub-alpine shrubland over the surface mine & dump (the standard error of this estimate is approximately 0.054). In what follows we focus exclusively on the evidence base and network meta-analysis for the larger network of land covers in this ES.

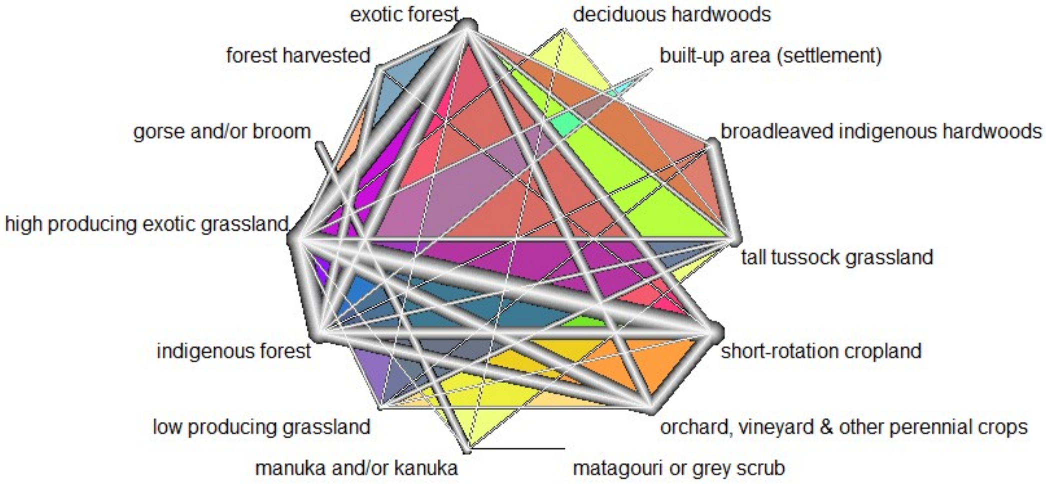


*Figure S10: Evidence network for land cover comparisons on nutrient cycling. Refer to Fig. S7 for an explanation of symbols and their interpretation.*

This evidence network is formed by 111 pairwise comparisons of 14 land covers. Data were obtained from 56 different studies, each contributing a minimum of one and a maximum of four pairwise land cover comparisons. As indicated by the thicker lines in Fig. S10, the land covers that are most commonly compared are:

- High producing exotic grassland (62 comparisons)
- Exotic forest (41 comparisons)
- Indigenous forest (34 comparisons)
- Short-rotation cropland (26 comparisons)

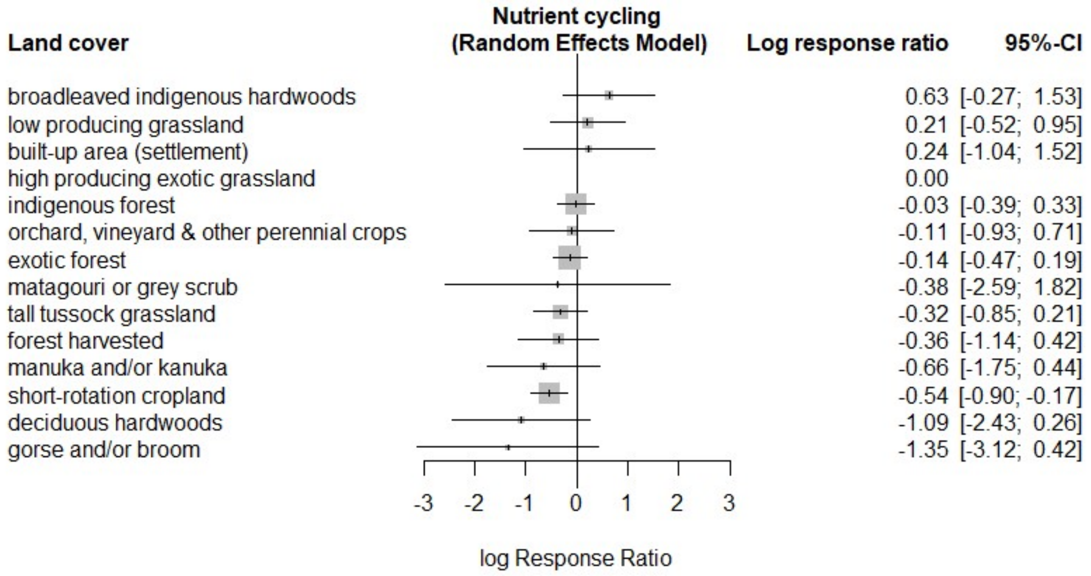


*Figure S11: Forest plot of land cover contrasts in the supply of nutrient cycling. Refer to Fig. S8 for details on how to interpret the symbols.*

**Measures of heterogeneity/network inconsistency:**

$\tau^2 = 0.702$
$I^2 = 90.341$

**Main results:**

- With the exception of short-rotation cropland, the confidence intervals for most land covers overlap the high producing exotic grassland reference. Moreover, exotic and indigenous forests also exhibit very small effect estimates, suggesting they may share similar nutrient cycling dynamics to those found in high producing exotic grasslands, where nutrient enrichment induces more dynamic processing rates in the soil system (Graaff *et al.*, 2006).
- Short-rotation croplands and other land covers dominated by exotic species and lacking nutrient enrichment (forest harvested, deciduous hardwoods) perform worse in the supply of this ES than the reference cover.
- The wider confidence intervals in the forest plot shown in Fig. S11 correspond to the land covers that had the fewest direct comparisons within the evidence network, while the land covers with narrower intervals are the ones that were informed by the greatest number of comparisons.

#### Habitat provision

**Type of indicators for this ES:** Comparisons for this ES were drawn from 83 indicators which, for the most part, expressed aspects relating to: the availability of resources and/or conditions favorable to wildlife within a land cover, habitat occupation or use by native fauna and the health of native animal species within a given land cover.

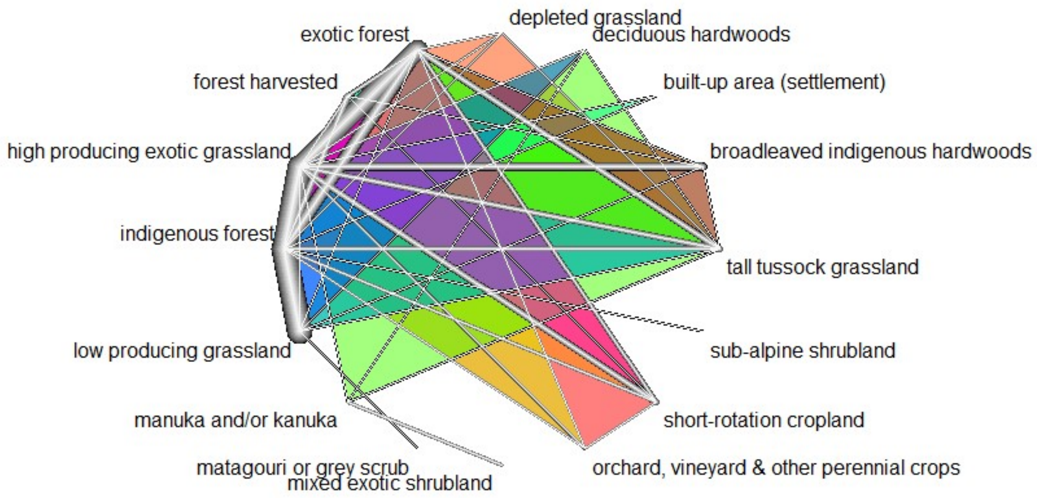


*Figure S12: Evidence network for land cover comparisons on habitat provision. Refer to Fig. S7 for an explanation of symbols and their interpretation.*

**Evidence base:** This evidence network is formed by 130 pairwise comparisons of 16 land covers. Data were obtained from 52 different studies, each contributing a minimum of one and a maximum of four pairwise land cover comparisons. As indicated by the thicker lines in Fig. S12, the land covers that are most commonly compared are:

- High producing exotic grassland (65 comparisons)
- Indigenous forest (53 comparisons)
- Exotic forest (50 comparisons)
- Tall tussock grassland (18 comparisons)

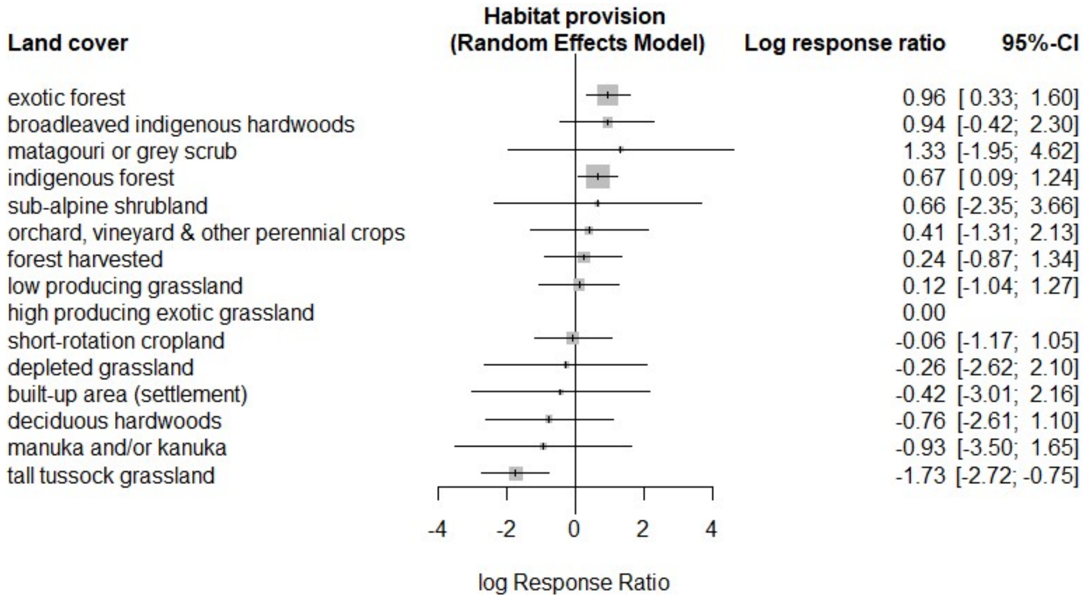


*Figure S13: Forest plot of land cover contrasts in habitat provision. Random effects model with high producing exotic grassland as a reference. Refer to Fig. S8 for details on how to interpret the symbols.*

**Measures of heterogeneity/network inconsistency:**

$\tau^2 = 1.493$
$I^2 = 99.535$

**Main results:**

- Exotic and indigenous forests are both significantly better than high producing exotic grassland in providing habitat and, although non-significant, the exotic forest ranks slightly higher than the indigenous one in supplying this ES.
- Tall tussock grasslands rank poorly and are significantly worse than both exotic and indigenous forests and high producing exotic grasslands in the provision of habitat.
- All croplands and grasslands (low, high and depleted) perform similarly in the supply of this ES and, overall, rank below the forest and native shrublands.

#### Soil formation

**Type of indicators for this ES:** Comparisons for this ES were drawn from 121 different indicators that cover aspects such as: soil aggradation and degradation processes (the latter having a negative effect on soil formation), soil structure and stability, the availability of nutrients and favorable conditions for plant growth in the soil.

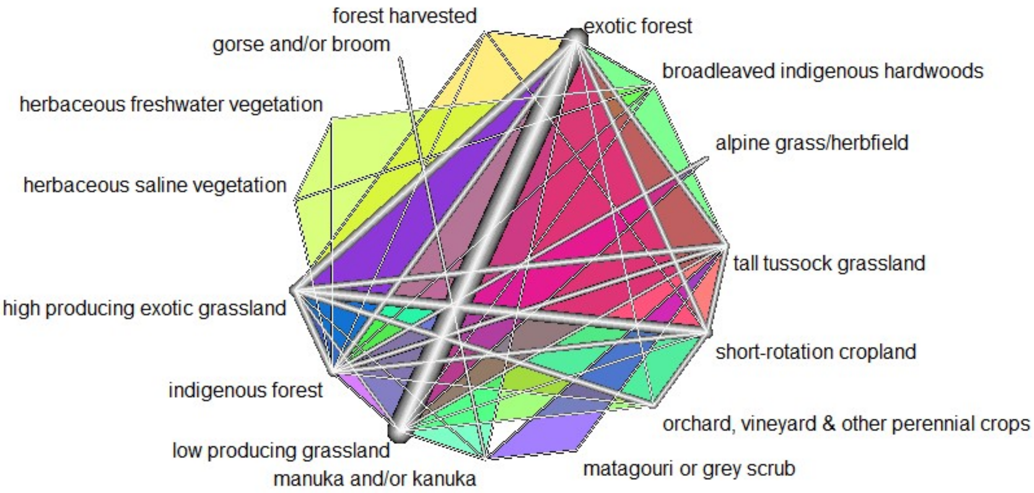


*Figure S14: Evidence network for land cover comparisons on soil formation. Refer to Fig. S7 for an explanation of symbols and their interpretation.*

**Evidence base:** This evidence network is formed by 112 pairwise comparisons of 15 land covers. Data were obtained from 51 different studies, each contributing a minimum of one and a maximum of four pairwise land cover comparisons. As indicated by the thicker lines in Fig. S14, the land covers that are most commonly compared are:

- High producing exotic grassland (54 comparisons)
- Exotic forest (45 comparisons)
- Short-rotation cropland (31 comparisons)
- Indigenous forest (25 comparisons)

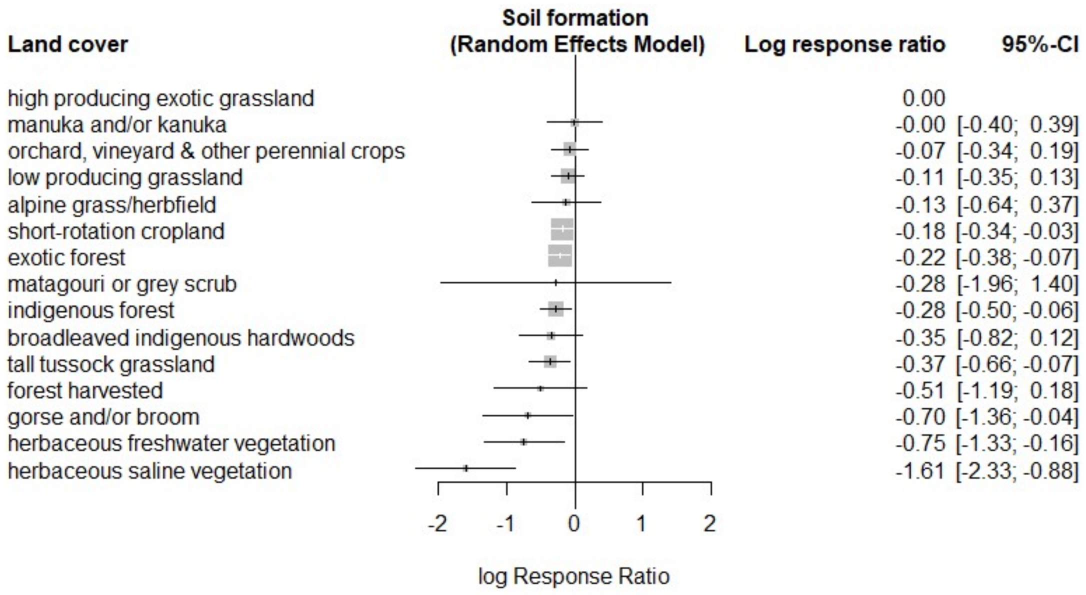


*Figure S15: Forest plot of land cover contrasts in the supply of soil formation. Random effects model with high producing exotic grassland as a reference. Refer to Fig. S8 for details on how to interpret the symbols.*

**Measures of heterogeneity/network inconsistency:**

$\tau^2$ = 0.244
$I^2$ = 65.874

**Main results:**

- No land cover is significantly better than the high producing exotic grassland in promoting soil formation. This is likely a result of both the high artificial nutrient inputs (that result in a greater nutrient availability for plants and, by our accounting, increased soil formation) and the fact that this land cover tends to be found in areas where the soils are well developed and very favorable to plant growth. The small (and statistically non-significant) differences between high producing exotic grassland and both cropland covers (short rotation and orchard, vineyard & other perennial crops) could be explained by similar factors.

- Except for manuka and/or kanuka, all native land covers rank similarly in the supply of this ES. Indigenous forests and tall tussock grasslands are significantly worse than the reference land cover, while broadleaved indigenous hardwoods and matagouri also do worse than the reference, but not significantly so.

- Short-rotation cropland, exotic forest and gorse and/or broom are significantly worse than the reference land cover in supplying this ES. Similarly, harvested forest also ranks below the reference land cover, but its wide confidence intervals make this difference statistically non-significant. Wide confidence intervals also apply to herbaceous freshwater and saline vegetation which, nevertheless, still perform significantly worse than the reference. For the freshwater and saline vegetation this makes sense given how they are prone to influxes of water that prevent soil forming processes.

##### Freshwater provision

**Type of indicators for this ES:** Comparisons for this ES were drawn from 66 indicators , all of which expressed a measure of land cover effects on the quantity or quality of freshwater supplied by streams. Indicators on the quality of water draw mostly on measures from concentrations of nutrients commonly linked to eutrophication (namely nitrogen and phosphorus), sediments and fecal contamination.

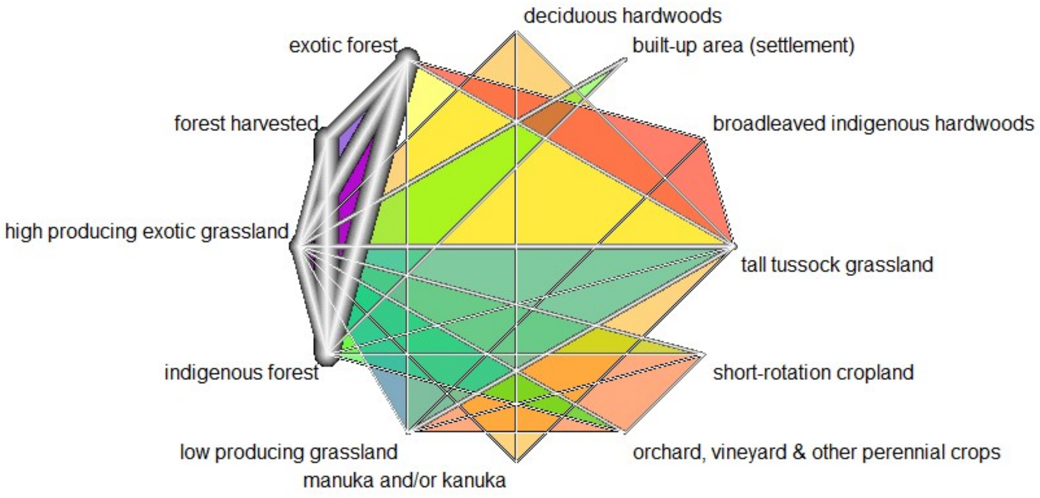


*Figure S16: Evidence network for land cover comparisons on freshwater provision. Refer to Fig. S7 for an explanation of symbols and their interpretation.*

**Evidence base:** This evidence network is formed by 88 pairwise comparisons of 12 land covers. Data were obtained from 40 different studies, each contributing a minimum of one and a maximum of three pairwise land cover comparisons. As indicated by the thicker lines in Fig. S16, the land covers that are most commonly compared are:

- High producing exotic grassland (46 comparisons)
- Indigenous forest (37 comparisons)
- Exotic forest (32 comparisons)
- Tall tussock grassland (16 comparisons)

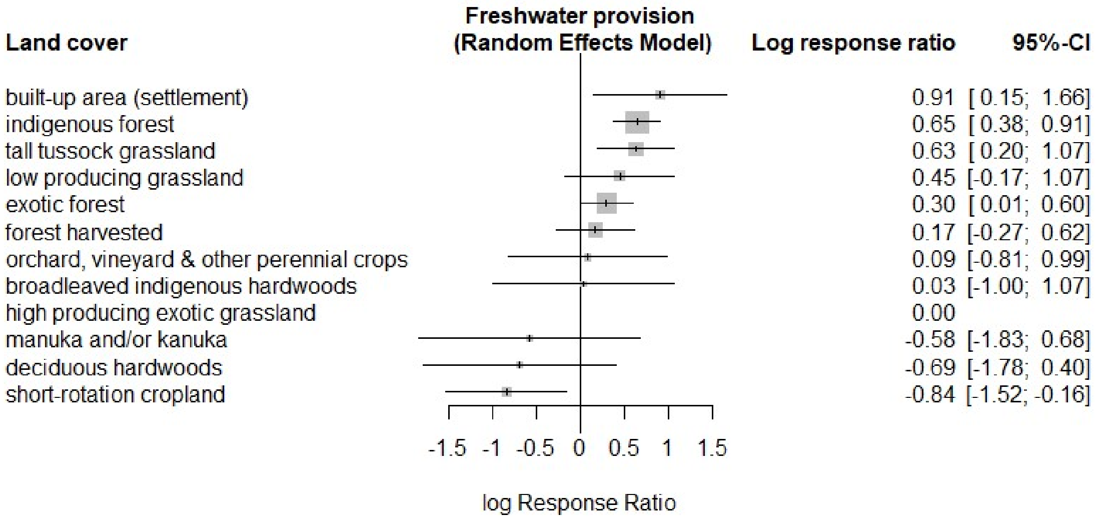


*Figure S17: Forest plot of land cover contrasts on freshwater provision. Random effects model with high producing exotic grassland as a reference. Refer to Fig. S8 for details on how to interpret the symbols.*

**Measures of heterogeneity/network inconsistency:**

$\tau^2$ = 0.505
$I^2$ = 85.984

**Main results:**

- It is striking to find built-up area as the highest ranking land cover in supplying this ES. However the log Response estimate is bounded by wide confidence intervals. For this specific land cover we only had information on how it compares to high producing exotic grassland in terms of specific stream flow, where it exceeds the grassland by 4 times. The effect we see here is, therefore, likely an artifact of this single data point.
- Tussock grasslands and indigenous forests both perform significantly better than the high producing exotic grassland in providing freshwater. While low producing grasslands also tended to do better than the reference, but not significantly so.
- Exotic and harvested forests have very similar rankings, performing slightly better than the reference, but not in a statistically significant way.
- Short-rotation cropland performs poorly in the supply of this ES and has significant differences not only with the reference land cover, but also with some of the ones that rank high in its supply (built-up area, tall tussock grassland, exotic and indigenous forests).
- For the remainder of the land covers, the confidence intervals are wide and intersect those of all the other land covers.

#### Water purification

**Type of indicators for this ES:** Comparisons for this ES were drawn from 78 indicators that provide a measure of either: the filtering of pollutants and excess nutrients from the water (either in the soil or in aquatic systems, for processes that are affected by land cover), or the accumulation of pollutants and toxic substances in freshwater.

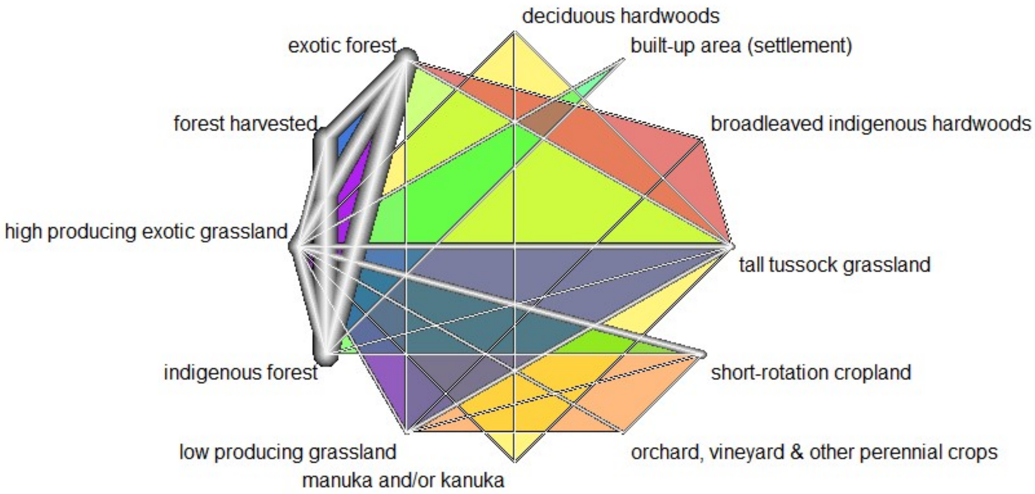


*Figure S18: Evidence network for land cover comparisons on water purification. Refer to Fig. S7 for an explanation of symbols and their interpretation.*

**Evidence base:** This evidence network is formed by 78 pairwise comparisons of 12 land covers. Data were obtained from 40 different studies, each contributing a minimum of one and a maximum of three pairwise land cover comparisons. As indicated by the thicker lines in Fig. S18, the land covers that are most commonly compared are:

- High producing exotic grassland (40 comparisons)
- Indigenous forest (30 comparisons)
- Exotic forest (27 comparisons)
- Tall tussock grassland (16 comparisons)

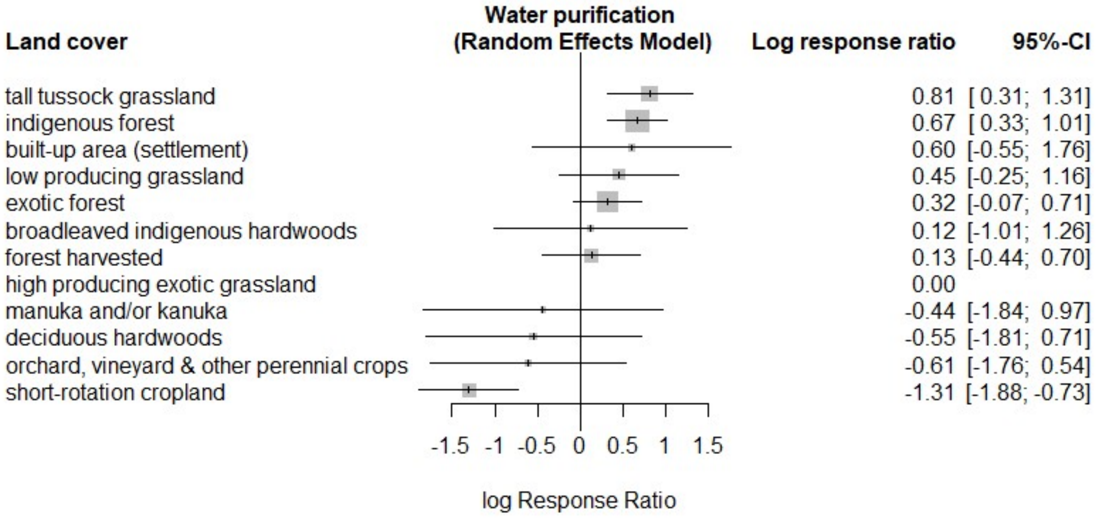


*Figure S19: Forest plot of land cover contrasts in the supply of water purification. Random effects model with high producing exotic grassland as a reference. Refer to Fig. S8 for details on how to interpret the symbols.*

**Measures of heterogeneity/network inconsistency:**

$\tau^2$ = 0.62
$I^2$ = 90.627

**Main results:**

- Both tall tussock grasslands and indigenous forests stand out as land covers that do significantly better than the reference in the supply of this ES, and overall, rank above all other land covers. Exotic and harvested forests also tend to perform slightly better than the reference land cover in this ES, although not statistically significantly so.
- Croplands & high producing grasslands rank poorly, with short rotation croplands performing significantly worse than many of the land covers, including the high producing grasslands.
- Broadleaved indigenous hardwoods, manuka and/or kanuka, deciduous hardwoods and orchard, vineyard & other perennial crops all have wide confidence intervals and, as a result, are not significantly different from the other land covers.
- The relatively high ranking estimate for built areas is quite surprising here. However, the estimate is bounded by large confidence intervals and supported by direct comparisons with only two other land covers (high producing exotic grassland and indigenous forest), both of which stem from a single study.

#### Global climate regulation

**Type of indicators for this ES:** Comparisons for this ES were drawn from 31 indicators, most of which are based on measures of greenhouse gas emission and sequestration processes in the soil. The majority of the indicators focus on carbon dioxide, however, the data also include a few on methane and nitrous oxide.

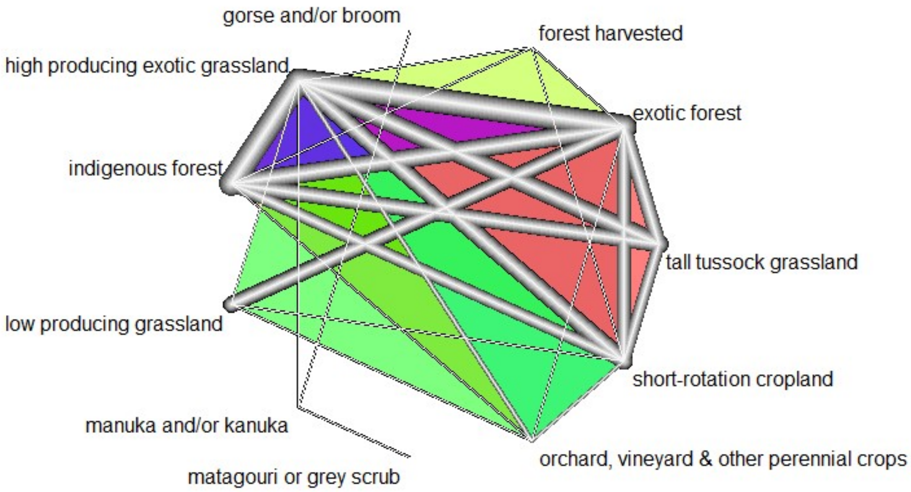


*Figure S20: Evidence network for land cover comparisons on global climate regulation. Refer to Fig. S7 for an explanation of symbols and their interpretation.*

**Evidence base:** This evidence network is formed by 75 pairwise comparisons of 11 land covers. Data were obtained from 33 different studies, each contributing a minimum of one and a maximum of four pairwise land cover comparisons. As indicated by the thicker lines in Fig. S20, the land covers that are most commonly compared are:

- High producing exotic grassland (45 comparisons)
- Exotic forest (34 comparisons)
- Short-rotation cropland (25 comparisons)
- Indigenous forest (19 comparisons)

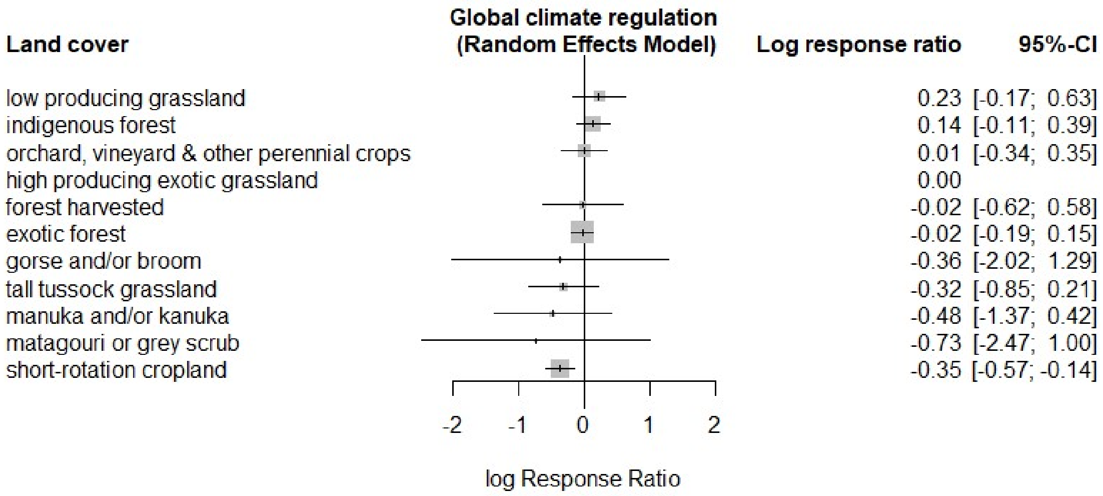


*Figure S21: Forest plot of land cover contrasts in the supply of global climate regulation. Random effects model with high producing exotic grassland as a reference. Refer to Fig. S8 for details on how to interpret the symbols.*

**Measures of heterogeneity/network inconsistency:**

$\tau^2$ = 0.335
$I^2$ = 94.579

**Main results:**

- There are no significant differences between the high producing exotic grassland reference and all other land covers, except for short-rotation croplands which performs significantly worse than the reference and indigenous forest in supplying this ES. The fact that short-rotation cropland does worse than the high producing grassland, and that the latter is not significantly different from the native land covers could, in part, be explained by the selection of indicators available for this ES (which focus mainly on processes at the soil level instead of the entire land system which would include the effects of livestock).

#### Primary production

**Type of indicators for this ES:** Comparisons for this ES were drawn from 22 indicators. The larger proportion of these indicators express primary productivity as the amount of biomass within a given land cover, however, there are also some indicators on the effects land covers have over primary productivity in streams (e.g., by providing more or less shade cover).

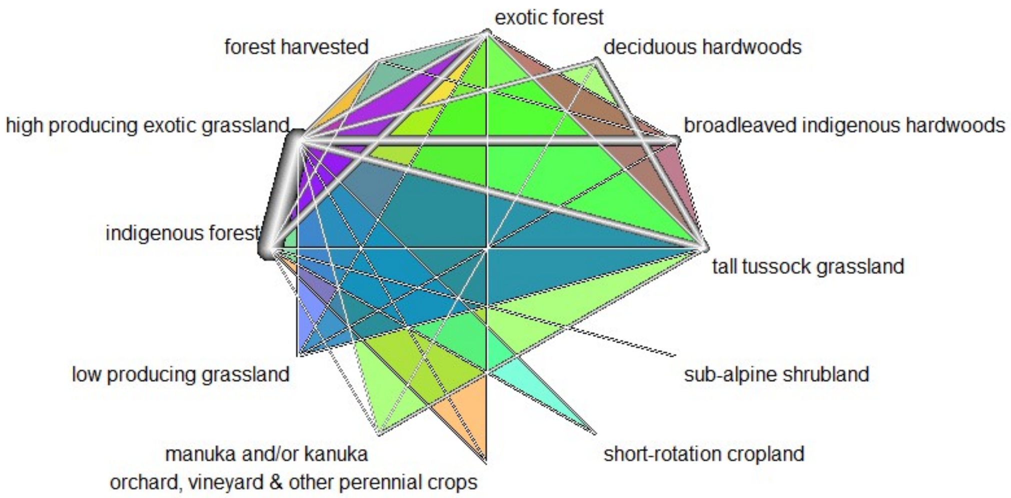


*Figure S22: Evidence network for land cover comparisons on primary production. Refer to Fig. S7 for an explanation of symbols and their interpretation.*

**Evidence base:** This evidence network is formed by 69 pairwise comparisons of 12 land covers. Data were obtained from 32 different studies, each contributing a minimum of one and a maximum of three pairwise land cover comparisons. As indicated by the thicker lines in Fig. S22, the land covers that are most commonly compared are:

- High producing exotic grassland (34 comparisons)
- Indigenous forest (27 comparisons)
- Exotic forest (25 comparisons)
- Tall tussock grassland (15 comparisons)

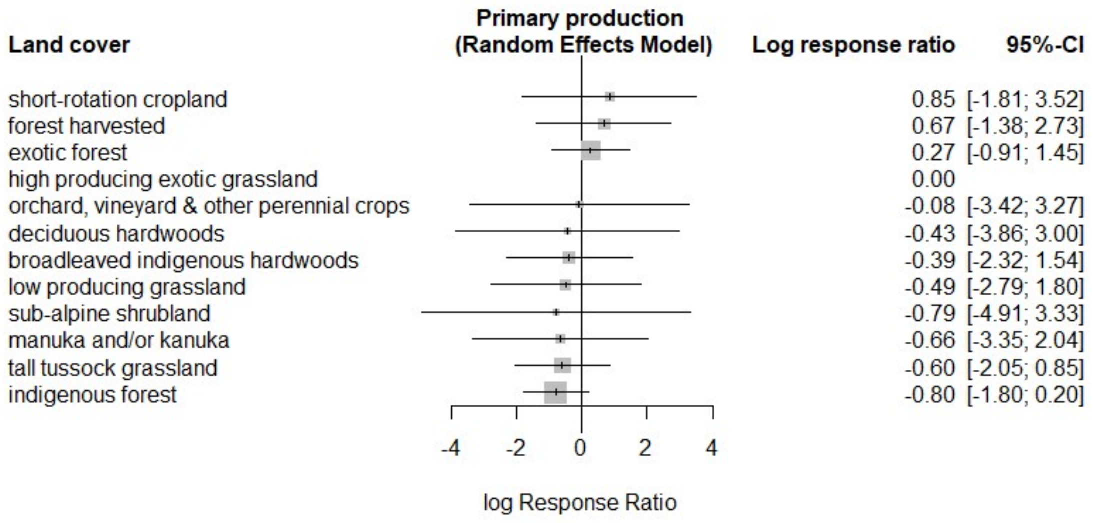


*Figure S23: Forest plot of land cover contrasts in the supply of primary production. Random effects model with high producing exotic grassland as a reference. Refer to Fig. S8 for details on how to interpret the symbols.*

**Measures of heterogeneity/network inconsistency:**

$\tau^2$ = 2.028
$I^2$ = 99.618

**Main results:**

- All of the land cover effect estimates for this ES have wide confidence intervals that overlap each other and intersect the reference mark for the high producing exotic grassland. However, the production land covers (exotic and harvested forests, croplands, deciduous hardwoods and high producing exotic grasslands) tend to be better at supplying this ES than the native ones (broadleaved indigenous hardwoods, tall tussock grassland, sub-alpine shrubland, manuka and/or kanuka and indigenous forest). This is likely due to the high biomass turnover found in production systems and reflected in the measures of biomass accumulation we have for this ES. Under the LCDB definition, forest harvested includes areas where indigenous and exotic forests have been cleared and within those, areas where the forest has been replanted and is up to 5 years old. The fast growth rate of young forests could thus account for the high rank of this land cover in the supply of primary production.

##### Water cycling

**Type of indicators for this ES:** Comparisons for this ES were drawn from 13 indicators, all of which quantify the amount of water flowing through the different components of the terrestrial segment of the cycle. For this ES we defined a positive relationship between all measures of water flow and the supply of the ES, such that the greater the flow at a given land cover, the larger the contribution of that land cover to the supply of the ES. This follows the MEA definition of water cycling as a supporting ES that benefits all living organisms by allowing the movement of water through ecosystems (MEA, 2005).

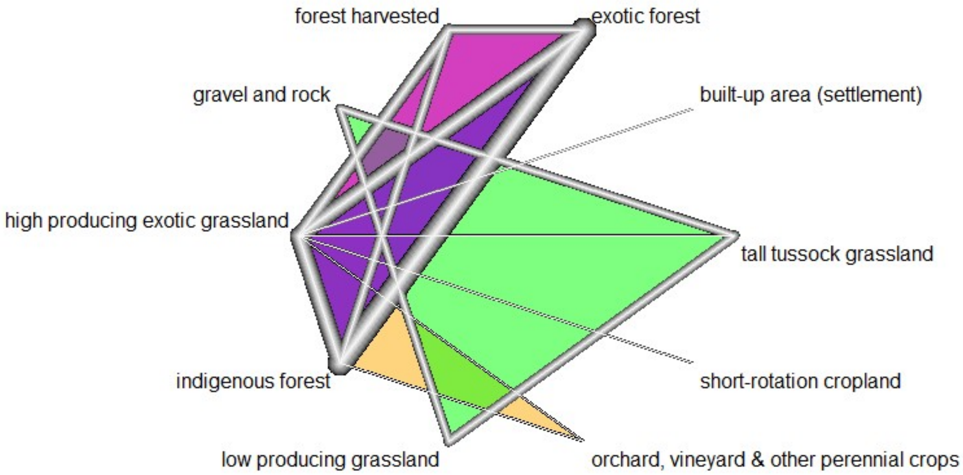


*Figure S24: Evidence network for land cover comparisons on water cycling. Refer to Fig. S7 for an explanation of symbols and their interpretation.*

**Evidence base:** This evidence network is formed by 42 pairwise comparisons of 10 land covers. Data were obtained from 21 different studies, each contributing a minimum of one and a maximum of three pairwise land cover comparisons. As indicated by the thicker lines in Fig. S24, the land covers that are most commonly compared are:

- High producing exotic grassland (26 comparisons)
- Exotic forest (19 comparisons)
- Indigenous forest (19 comparisons)
- Forest harvested (6 comparisons)

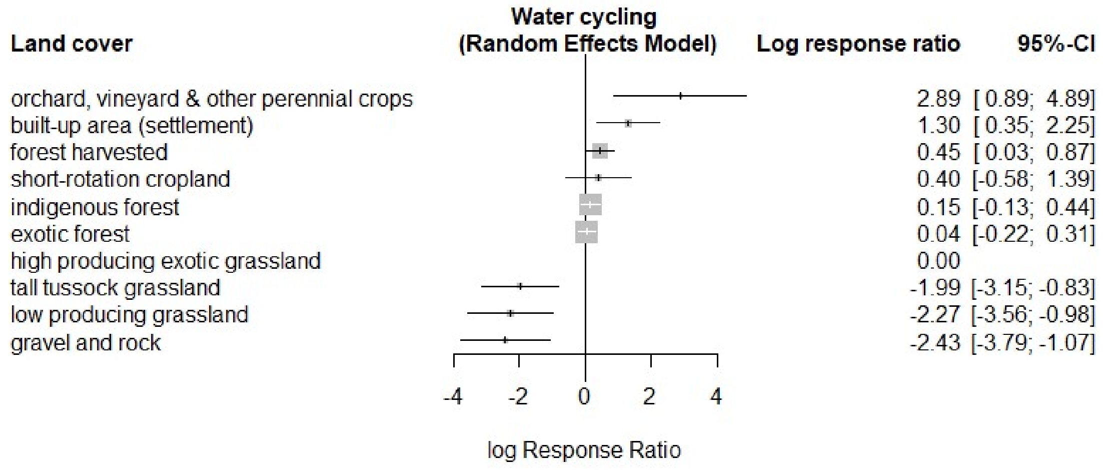


*Figure S25: Forest plot of land cover contrasts in the supply of water cycling. Random effects model with high producing exotic grassland as a reference. Refer to Fig. S8 for details on how to interpret the symbols.*

**Measures of heterogeneity/network inconsistency:**

$\tau^2 = 0.368$
$I^2 = 82.434$

**Main results:**

- Orchard, vineyard & other perennial crops, built-up area and forest harvested stand out as land covers that do very well in supplying this ES. Their effect estimates stand out as significantly better than the high producing exotic grassland reference and most of the other land covers (except harvested forest and short-rotation cropland). For built-up (settlement) areas, this could be an effect of increased flow speeds due to the presence of impervious surfaces.

- Tall tussock and low producing grasslands tend to rank worse in this ES than they do in the regulation of water timing and flows, suggesting a trade-off between water cycling and the regulation of flows for these systems. However, gravel and rock, which would be a system with faster cycling rates, also follows this trend. This could be driven by the direct evidence we have for gravel and rock under water cycling which involves water yield comparisons to tall tussock and low producing grasslands, and in which gravel and rock always performs worse (given its high infiltration capacity). In contrast, all other land covers with faster cycling rates (orchard, vineyard and other perennials and short-rotation cropland) are better at water cycling than they are at the timing and regulation of flows.

#### Erosion control

**Type of indicators for this ES:** Comparisons for this ES were drawn from 27 indicators on the magnitude of soil loss and sediment export to waterways as well as soil and stream channel characteristics that provide increased resistance to erosion.

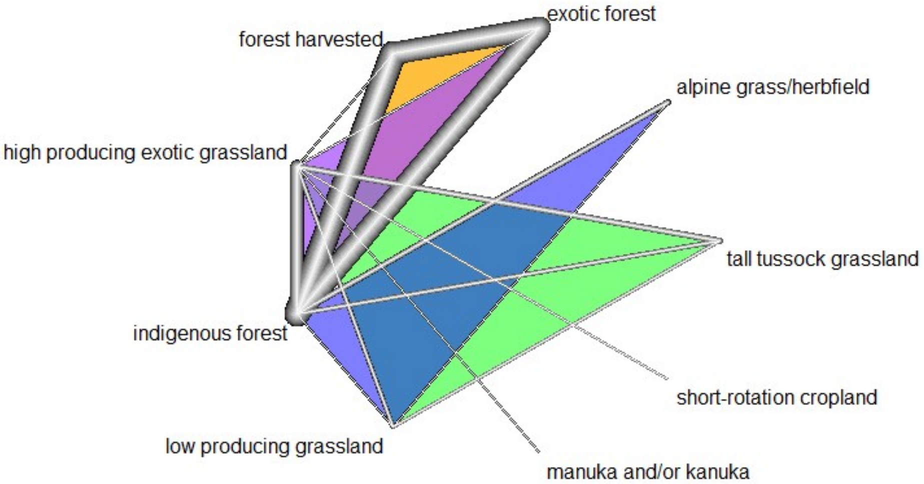


*Figure S26: Evidence network for land cover comparisons on erosion control. Refer to Fig. S7 for an explanation of symbols and their interpretation.*

**Evidence base:** This evidence network is formed by 34 pairwise comparisons of nine land covers. Data were obtained from 22 different studies, each contributing one to two pairwise land cover comparisons. As indicated by the thicker lines in Fig. S26, the land covers that are most commonly compared are:

- High producing exotic grassland (18 comparisons)
- Indigenous forest (17 comparisons)
- Exotic forest (10 comparisons)
- Forest harvested (six comparisons)

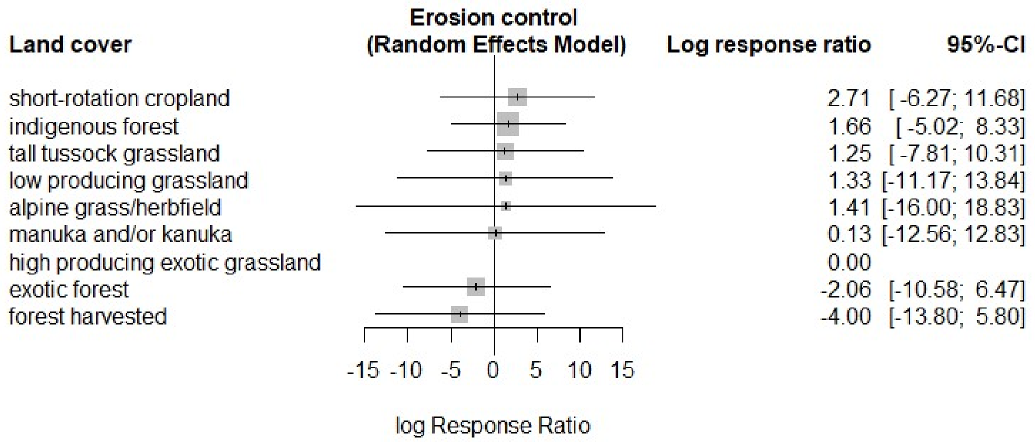


*Figure S27: Forest plot of land cover contrasts in the supply of erosion control. Random effects model with high producing exotic grassland as a reference. Refer to Fig. S8 for details on how to interpret the symbols.*

**Measures of heterogeneity/network inconsistency:**

$\tau^2$ = 9.155
$I^2$ = 99.969

**Main results:**

- Effect estimates are all bounded by wide confidence intervals, which yield no significant differences between any of the land covers. This could be due to the effect of environmental variables such as slope and parent material of the soil, which introduce additional variation within each land cover.
- Except for short-rotation cropland, the production land covers tend to perform poorly in the supply of this ES. In contrast, those with native vegetation covers (in the form of grass, forest or shrublands) have higher ranking estimates for their control over erosion processes.

#### Pest regulation

**Type of indicators for this ES:** Comparisons for this ES were drawn from 41 indicators, most of which focus on the abundance of invasive species in different land covers. However, there are also some indicators that quantify habitat occupation and use by invader species, and their response to biological controls.

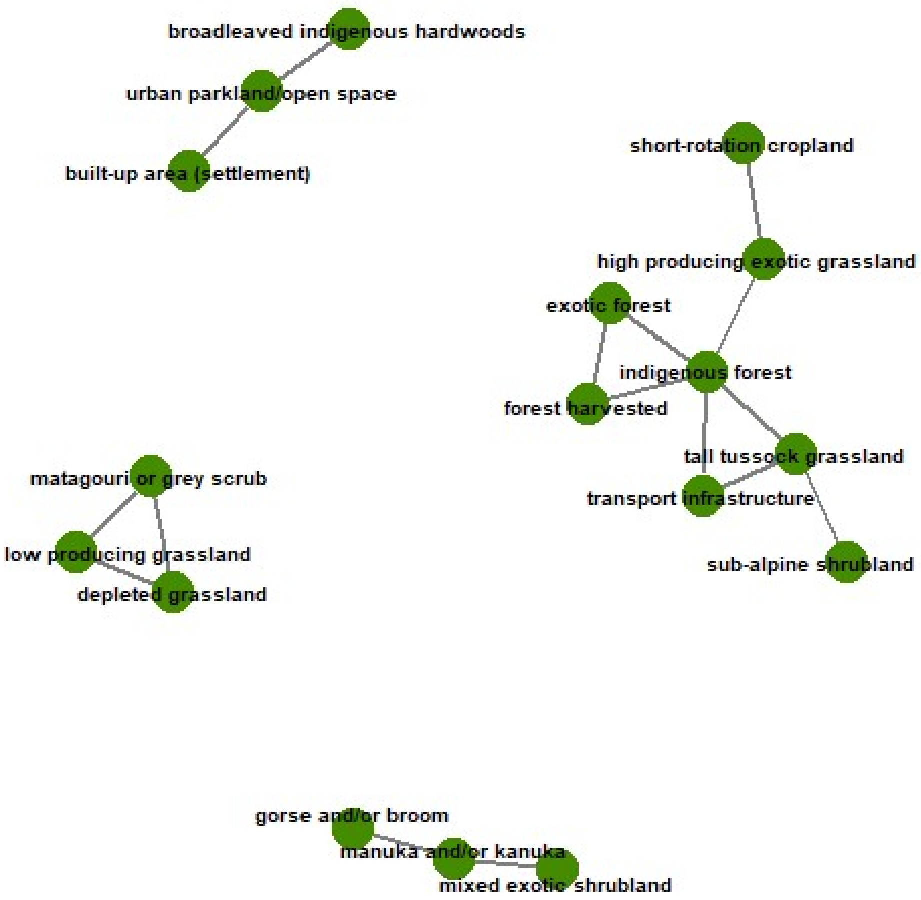


*Figure S28: Land-cover comparison networks for pest regulation. For this service, land cover comparisons were split into the four networks depicted here, however, only the comparisons in the larger network were used as evidence base for the network meta-analysis. Figure S29 presents the detailed configuration of this evidence base.*

**Evidence base:** As shown in Fig. S28, there are four networks connecting the land cover comparisons for this ES: three smaller ones and a larger one. The evidence for the smaller networks is summarized in Table S5, whereas the remainder of this section describes the data and analysis for the larger network of land covers in this ES.

Table S2: Reported response ratios for evidence subnetworks in pest regulation. Ratios are based on the natural logarithm of the quotient of land cover 1 over land cover 2

| Sub-network | Land Cover 1 | Land Cover 2 | Log Response Ratio | Standard Error | Study ID |
|---|---|---|---|---|---|
| 1 | broadleaved indigenous hardwoods | urban parkland/open space | -0.045 | 0.206 | S14626 |
| 1 | built-up area (settlement) | urban parkland/open space | 8.760 | 0.126 | S16437 |
| 2 | gorse and/or broom | manuka and/or kanuka | -0.572 | 0.304 | S14266 |
| 2 | manuka and/or kanuka | mixed exotic shrubland | 0.117 | 0.065 | S18250 |
| 3 | depleted grassland | low producing grassland | 0.105 | 0.593 | S13628 |
| 3 | depleted grassland | matagouri or grey scrub | 0.063 | 0.550 | S13628 |
| 3 | low producing grassland | matagouri or grey scrub | -0.042 | 0.424 | S13628 |

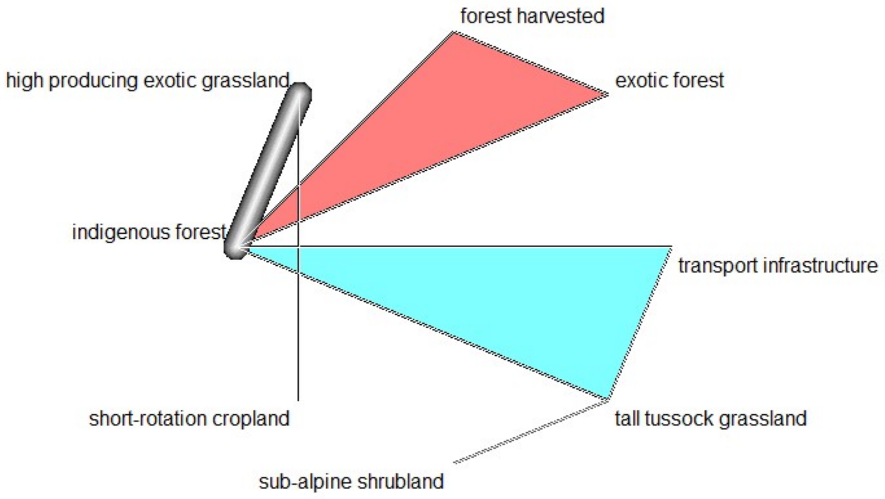


*Figure S29: Evidence network for land cover comparisons on pest regulation. Refer to Fig. S7 for an explanation of symbols and their interpretation.*

The largest evidence network for pest regulation is formed by 11 pairwise comparisons of eight land covers. Data were obtained from seven different studies, each contributing a minimum of one and a maximum of two pairwise land cover comparisons. As indicated by the thicker lines in Fig. S29, the land covers that are most commonly compared are:

- Indigenous forest (seven comparisons)
- Tall tussock grassland (three comparisons)
- Forest harvested (three comparisons)
- Exotic forest (three comparisons)

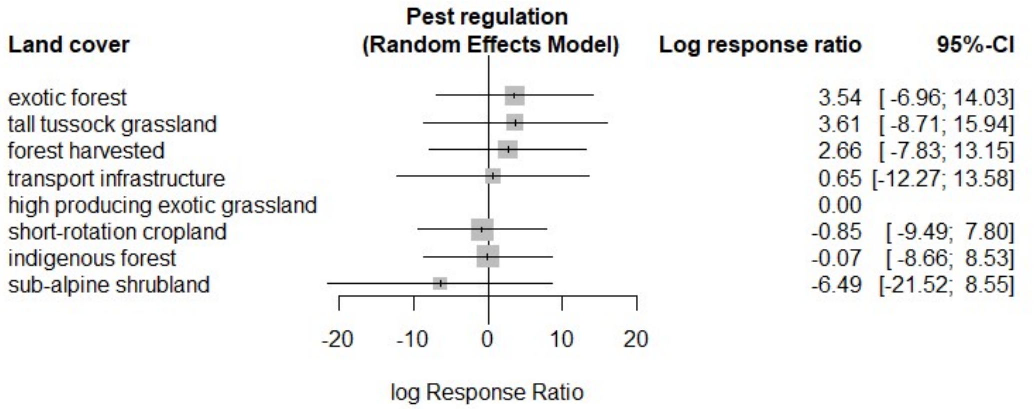


*Figure S30: Forest plot of land cover contrasts in the supply of pest regulation. Random effects model with high producing exotic grassland as a reference. Refer to Fig. S8 for details on how to interpret the symbols.*

**Measures of heterogeneity/network inconsistency:**

$\tau^2$ = 4.384
$I^2$ = 95.105

**Main results:**

- Effect estimates are all bounded by wide confidence intervals which yield no significant differences between the ratio estimates for any of the land covers. This could be explained by the fact that the evidence network has many comparisons converging around indigenous forest (Fig. S29) and that five out of the seven studies in this ES provide evidence only on a single pair of land covers.
- Tall tussock grasslands and exotic forests (harvested and unharvested) have the highest log response ratio estimates, ranking above the high producing grassland reference in the supply of this ES. In contrast, indigenous forests rank similarly to short-rotation croplands with small differences from the reference while the remaining native land cover, sub-alpine shrubland, performs worse than all other land covers in this ES.

##### Waste treatment

**Type of indicators for this ES:** Comparisons for this ES were drawn from 18 indicators most of which provide a measure of the concentration and export of toxic compounds in the soil.

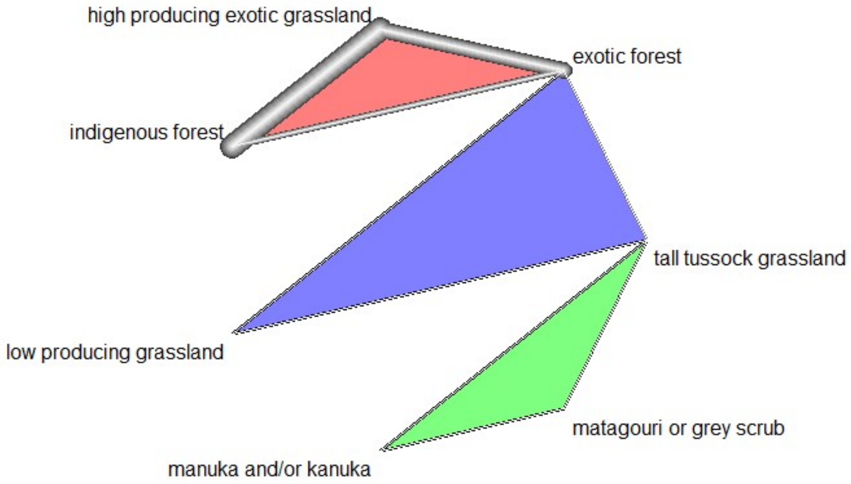


*Figure S31: Evidence network for land cover comparisons on waste treatment. Refer to Fig. S7 for an explanation of symbols and their interpretation.*

**Evidence base:** This evidence network is formed by 12 pairwise comparisons of seven land covers. Data were obtained from six different studies, each contributing one to two pairwise land cover comparisons. As indicated by the thicker lines in Fig. S31, the land covers that are most commonly compared are:

- Exotic forest (six comparisons)
- High producing exotic grassland (five comparisons)
- Tall tussock grassland (four comparisons)
- Indigenous forest (three comparisons)

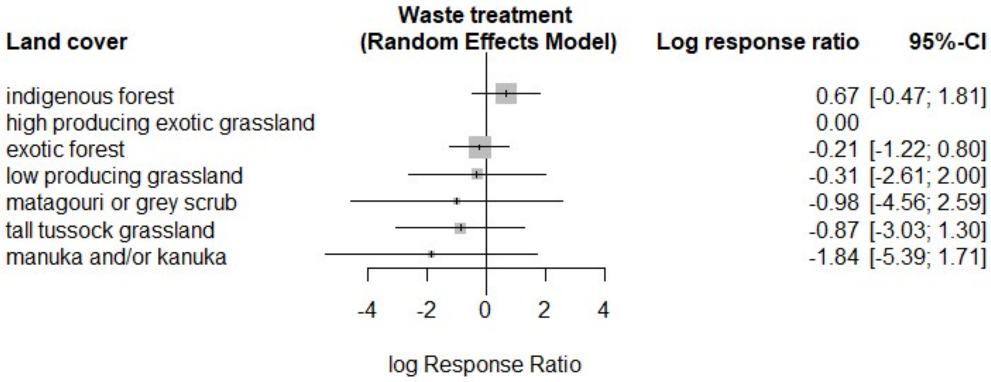


*Figure S32: Forest plot of land cover contrasts in the supply of waste treatment. Random effects model with high producing exotic grassland as a reference. Refer to Fig. S8 for details on how to interpret the symbols.*

**Measures of heterogeneity/network inconsistency:**

$\tau^2$ = 0.841
$I^2$ = 95.858

**Main results:**

- No significant differences can be found between the land covers in this ES ,since the confidence intervals all overlap each other and extend over both sides of the reference mark. Furthermore, the estimated log response ratio between each land cover and the high producing exotic grassland reference is always between one and negative one.
- Indigenous forest and high producing exotic grassland rank the highest in the supply of this ES, while exotic forest and the remaining native land covers (tall tussocks, manuka and/or kanuka, matagouri or grey scrub) all rank poorly and perform comparatively worse than the reference land cover.

#### Capture fisheries

**Type of indicators for this ES:** Comparisons for this ES were drawn from 22 indicators on the abundance, biomass, size and growth of freshwater fish.

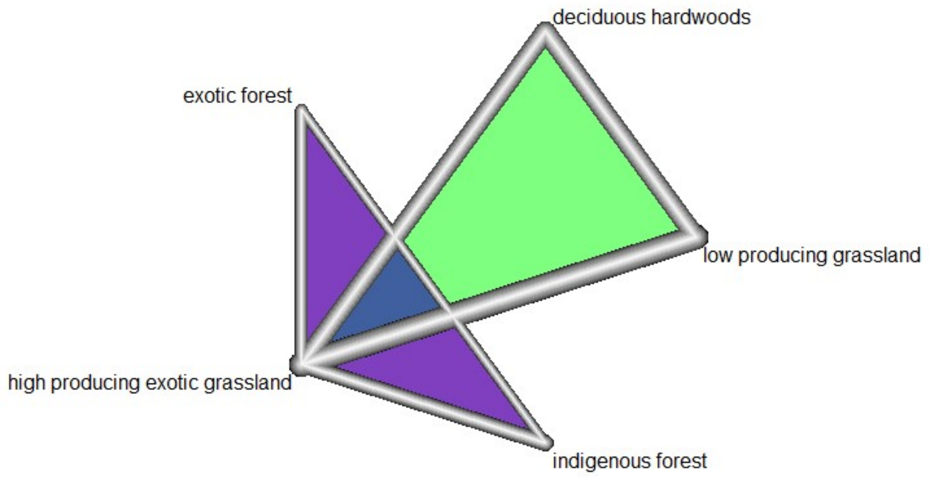


*Figure S33: Evidence network for land cover comparisons on capture fisheries. Refer to Fig. S7 for an explanation of symbols and their interpretation.*

**Evidence base:** This evidence network is formed by 11 pairwise comparisons of five land covers. Data were obtained from five different studies, each contributing one to two pairwise land cover comparisons. The land covers that are most commonly compared are:

- High producing exotic grassland (eight comparisons)
- Indigenous forest (five comparisons)
- Exotic forest (four comparisons)
- Low producing grassland (three comparisons)

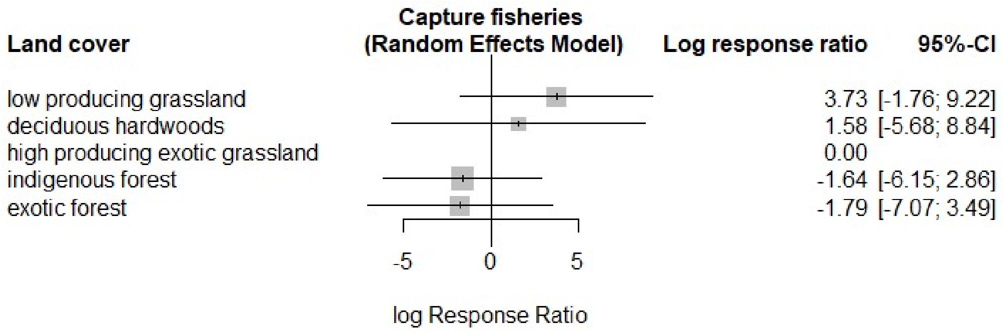


*Figure S34: Forest plot of land cover contrasts in the supply of capture fisheries. Random effects model with high producing exotic grassland as a reference. Refer to Fig. S8 for details on how to interpret the symbols.*

**Measures of heterogeneity/network inconsistency:**

$\tau^2$ = 3.955
$I^2$ = 99.399

**Main results:**

- The wide confidence intervals for all land covers preclude any significant differences between any of the land covers. However, in terms of the log response ratio estimates, both indigenous and exotic forests tend to supply less of the capture fisheries ES than the low and high producing grasslands and the deciduous hardwoods.

#### Ethical and spiritual values

**Type of indicators for this ES:** Comparisons for this ES were drawn from 13 indicators, all of which express the abundance, biomass, and growth of culturally valuable fauna, namely, long- and short-finned eels. This includes most of the indicators that were also used for the capture fisheries ES.

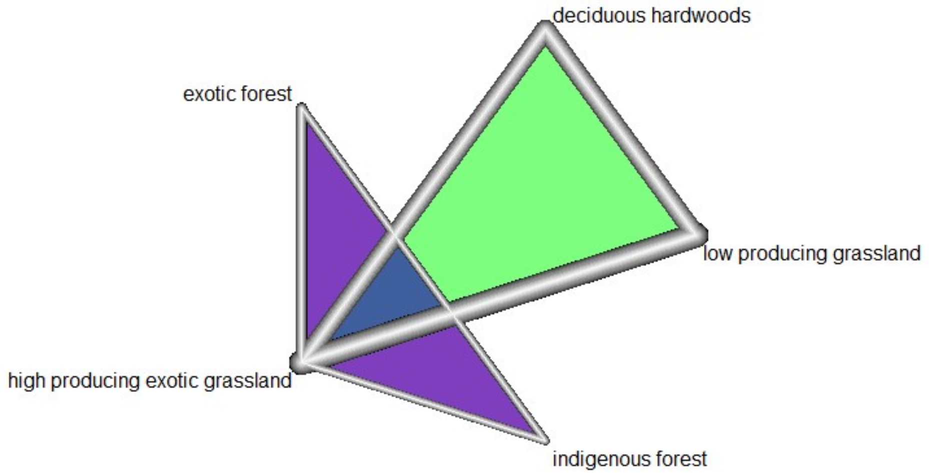


*Figure S35: Evidence network for land cover comparisons on ethical and spiritual values. Refer to Fig. S7 for an explanation of symbols and their interpretation.*

**Evidence base:** This evidence network is formed by 10 pairwise comparisons of five land covers. Data were obtained from four different studies, each contributing one to two pairwise land cover comparisons. The land covers that are most commonly compared are:

- High producing exotic grassland (seven comparisons)
- Indigenous forest (four comparisons)
- Exotic forest (four comparisons)
- Low producing grassland (three comparisons)

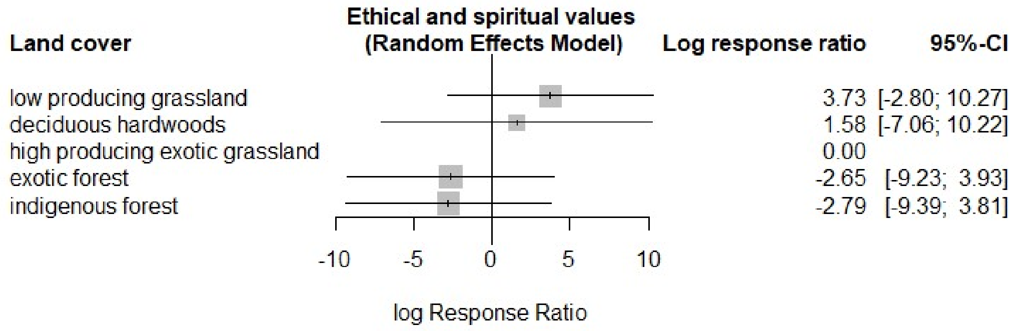


*Figure S36: Forest plot of land cover contrasts in the supply of ethical and spiritual values. Random effects model with high producing exotic grassland as a reference. Refer to Fig. S8 for details on how to interpret the symbols.*

**Measures of heterogeneity/network inconsistency:**

$\tau^2$ = 4.712
$I^2$ = 99.531

**Main results:**

- Land covers in this ES appear to share the same trends as those for the capture fisheries ES with wide confidence intervals and the estimated ratios for exotic and indigenous forests at the opposite end of the spectrum to those for the grasslands and deciduous hardwoods. This is an effect of the overlap between the indicators that support the evidence on this ES and that of capture fisheries.

#### Disease mitigation

**Type of indicators for this ES:** Comparisons for this ES were drawn from nine indicators on the abundance of mosquitoes and their predators, and on the presence of fecal coliforms, *Escherichia coli* or enteric coliforms in streams and freshwater sources.

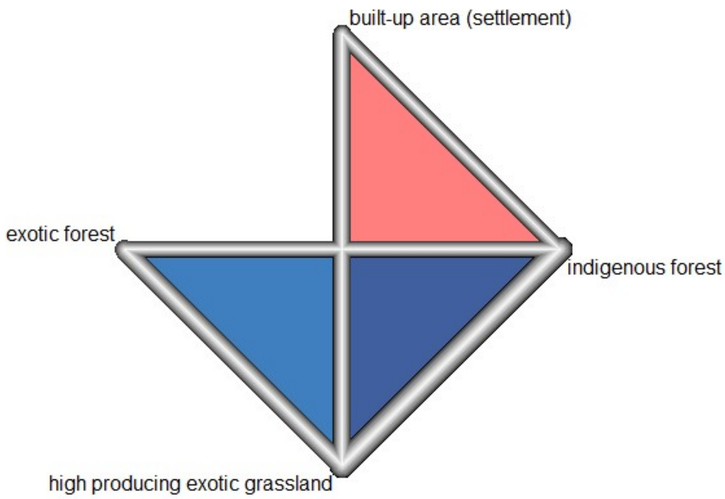


*Figure S37: Evidence network for land cover comparisons on disease mitigation. Refer to Fig. S7 for an explanation of symbols and their interpretation.*

**Evidence base:** This evidence network is formed by ten pairwise comparisons of four land covers. Data were obtained from four different studies, each contributing one to two pairwise land cover comparisons. The number of comparisons per land cover is as follows:

- Indigenous forest (seven comparisons)
- High producing exotic grassland (seven comparisons)
- Exotic forest (four comparisons)
- Built-up area (settlement) (two comparisons)

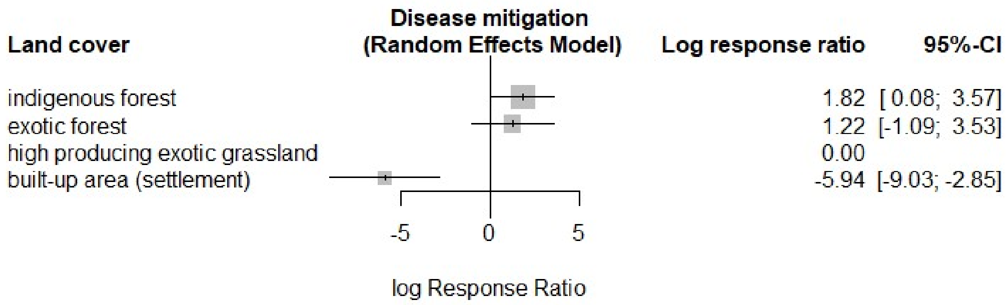


*Figure S38: Forest plot of land cover contrasts in the supply of disease mitigation. Random effects model with high producing exotic grassland as a reference. Refer to Fig. S8 for details on how to interpret the symbols.*

**Measures of heterogeneity/network inconsistency:**

$\tau^2$ = 1.729
$I^2$ = 95.505

**Main results:**

- Built-up area stands out as the land cover that performs significantly worse than any of the others in supplying this ES.
- Production land covers (exotic forest and high producing exotic grassland) perform slightly worse than the indigenous forest with respect to disease mitigation. The difference with the indigenous forest is significant for the high producing exotic grasslands but not for the exotic forest.

##### Pollination

**Type of indicators for this ES:** Comparisons for this ES were drawn from three indicators that quantify the potential for pollination in a land cover based on either the abundance of pollinators or on the flower visitation rates and duration.

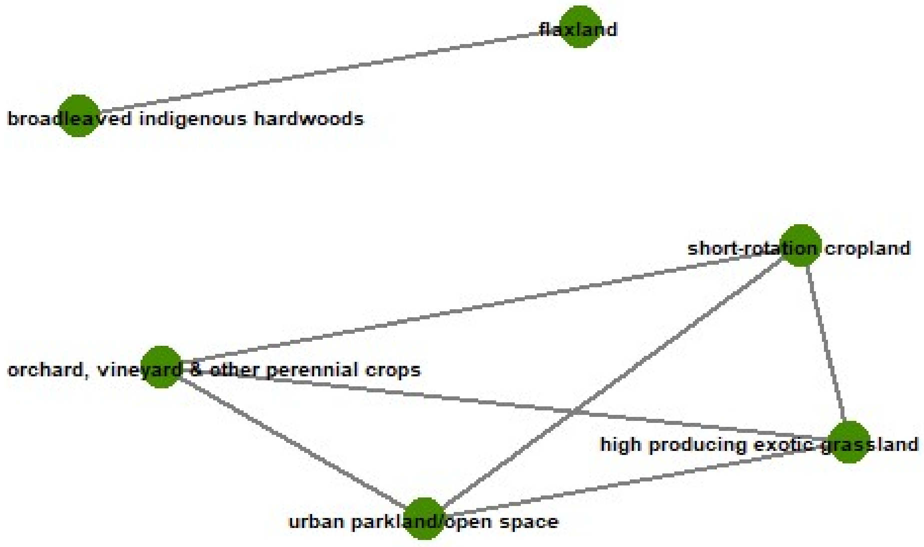


*Figure S39: Land-cover comparison networks for pollination. For this service, land cover comparisons were split into the two networks depicted here, however, only the comparisons in the larger network were used as evidence base for the network meta-analysis. Figure S40 presents the detailed configuration of this evidence base.*

**Evidence base:** As shown in Fig. S39, the evidence for this ES is split into two networks: a smaller one connecting broadleaved indigenous hardwoods and flaxland, and a larger one with the four remaining land covers. For the land covers in the smaller sub-network, the evidence available comes from a single study in which the log response ratio of broadleaved indigenous hardwoods to flaxland is approximately -0.823 and has a standard error of 0.625. Details for the larger network are presented below.

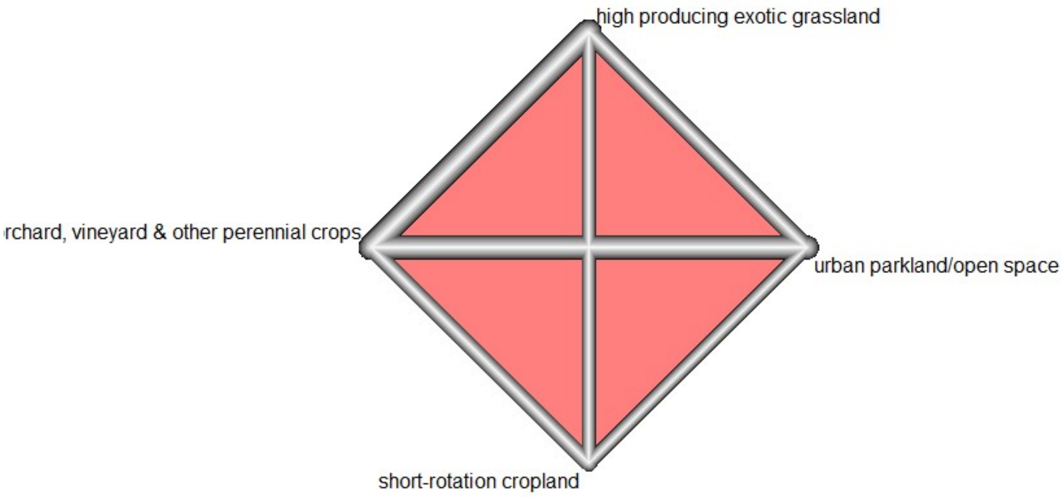


*Figure S40: Evidence network for land cover comparisons on pollination. Refer to Fig. S7 for an explanation of symbols and their interpretation.*

This evidence network is formed by seven pairwise comparisons of four land covers. Data were obtained from two different studies, one contributing one and the other three pairwise land cover comparisons. As indicated by the thicker lines in Fig. S40, the land covers that are most commonly compared are:

- Urban parkland/open space (four comparisons)
- High producing exotic grassland (four comparisons)
- Short-rotation cropland (three comparisons)
- Orchard, vineyard & other perennial crops (three comparisons)

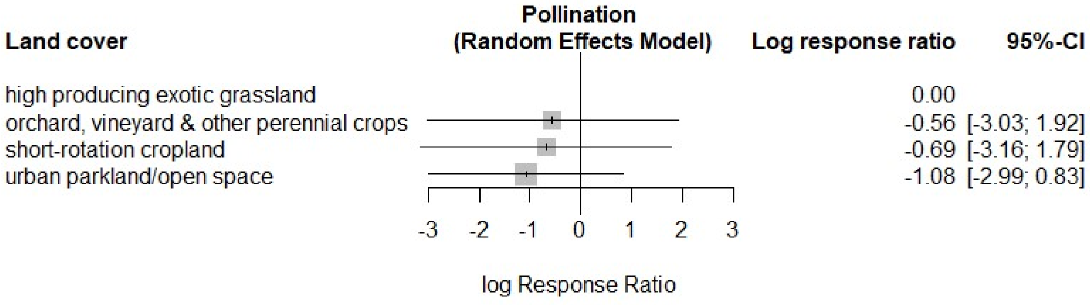


*Figure S41: Forest plot of land cover contrasts in the supply of pollination. Random effects model with high producing exotic grassland as a reference. Refer to Fig. S8 for details on how to interpret the symbols.*

**Measures of heterogeneity/network inconsistency:**

$\tau^2$ = 1.343
$I^2$ = 94.525

**Main results:**

- There are no significant differences between the different land covers in the supply of this ES, since all confidence intervals for the ratio estimates overlap each other and extend across the baseline reference. This is likely a result of having only 2 studies informing this analysis.
- The log response ratio estimates suggest that urban parkland/open spaces and, to a lesser extent, croplands perform worse than high producing exotic grasslands in supplying pollination ES.

#### Regional & local climate regulation

**Type of indicators for this ES:** Comparisons for this ES were drawn from three indicators that quantify the regulation of temperatures either in stream water or near the land surface (as expressed by evapotranspiration).

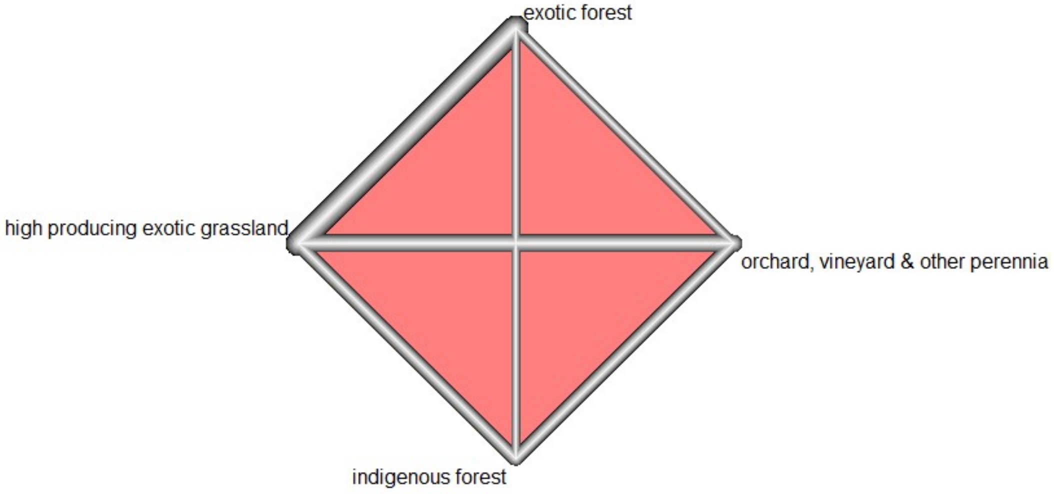


*Figure S42: Evidence network for land cover comparisons on regional and local climate regulation. Refer to Fig. S7 for an explanation of symbols and their interpretation.*

**Evidence base:** This evidence network is formed by seven pairwise comparisons of four land covers. Data were obtained from two different studies, one contributing one and the other three pairwise land cover comparisons. The number of comparisons per land cover is as follows:

- High producing exotic grassland (four comparisons)
- Exotic forest (four comparisons)

- Orchard, vineyard & other perennial crops (three comparisons)
- Indigenous forest (three comparisons)

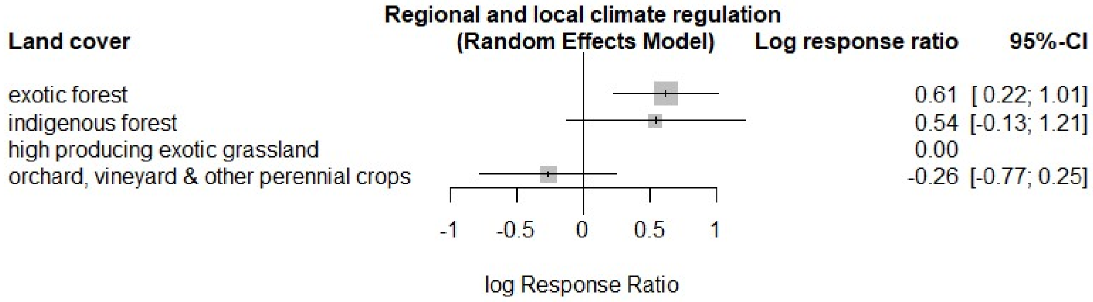


*Figure S43: Forest plot of land cover contrasts in the supply of regional and local climate regulation. Random effects model with high producing exotic grassland as a reference. Refer to Fig. S8 for details on how to interpret the symbols.*

**Measures of heterogeneity/network inconsistency:**

$\tau^2$ = Not available
$I^2$ = Not available

**Main results:**

- Besides exotic forest (which is significantly better than the reference land cover at supplying this ES), all other land covers have overlapping confidence intervals and, consequently, are not significantly different from each other in the supply of this ES. This is probably due to the limited number of studies with evidence on this ES.
- The log response ratio estimates suggest that forested land covers tend to supply greater climate regulation at the local and regional level than high producing exotic grassland and orchard, vineyard and other perennial crops.

#### A note on confidence intervals and the size of the evidence base

The forest plots above show that primary production, erosion control, pest regulation, waste treatment, capture fisheries, ethical & spiritual values, pollination and regional & local climate regulation all present wide, overlapping confidence intervals for all or most of their estimates. This suggests that differences in the supply of these ES across land covers were not significant. For some of these ES, this is due to smaller evidence bases, as in the case of pollination and regional & local climate regulation where the each network meta-analyses was informed by only 7 comparisons taken from 2 different studies. For ES with a slightly larger evidence base (e.g., capture fisheries, ethical and spiritual values, pest regulation and waste treatment) there is an asymmetry in the number of comparisons available for the different land covers, since one or two pairs of land covers harness most of the comparisons and leave limited evidence for the other pairs of comparisons (note the large differences in link weights in the corresponding evidence networks presented above for these ES). With over 60 pairwise comparisons across all land covers, the evidence base for primary production has a similar problem since most of the comparisons involve high producing exotic grasslands, indigenous and exotic forests. A similar trend can be observed with some (but not all) pairs of land covers for other well-informed ES, such as regulation of water timing and flows and soil formation. For waste treatment, the low sample size results from having few comparisons (12 in total) spread over a large number of land covers (7 in all), which results in an evidence network formed by several, poorly informed links.

#### Supplementary Results 3. Summary of log response ratios per land cover and ecosystem service combination

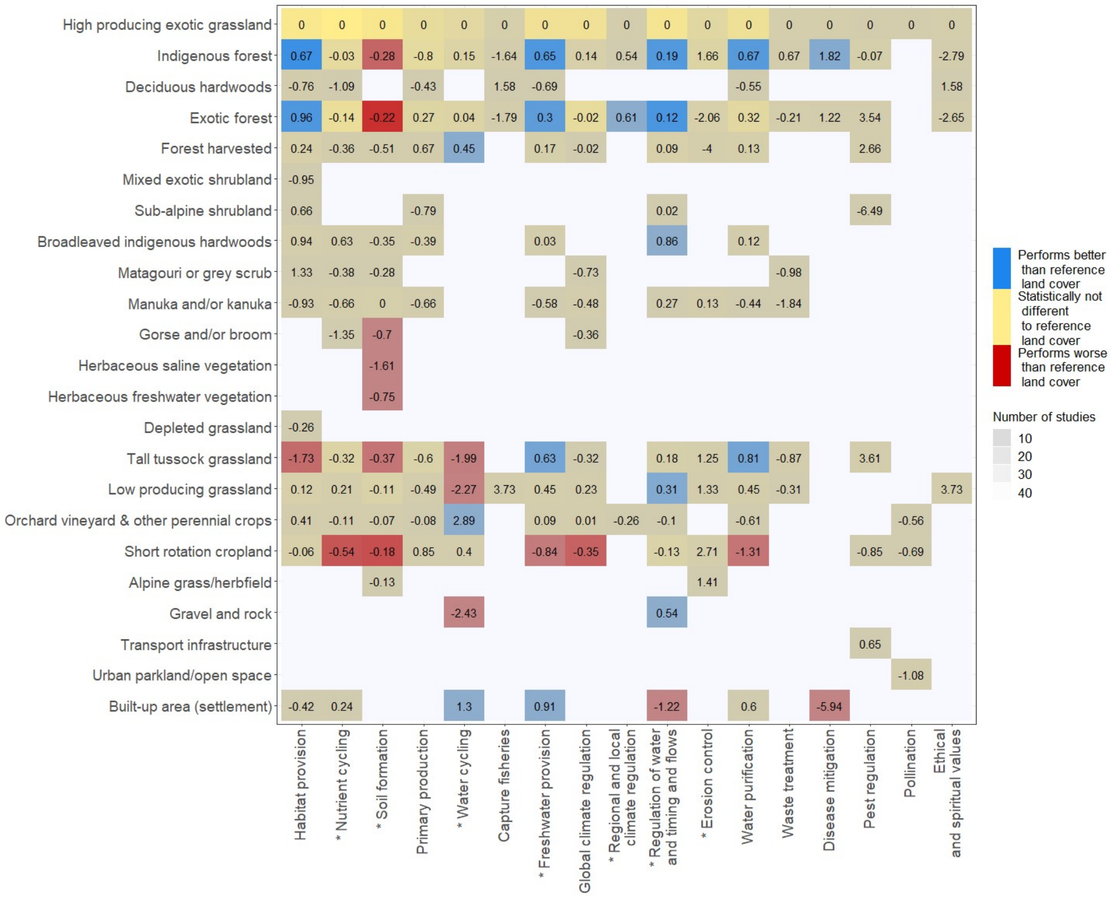


*Figure S44: Aggregated log response ratios of ES supply across land covers. Values are given relative to the high producing exotic grassland reference. Yellow tones highlight cases where the 95% confidence intervals for a land cover's log response ratio estimate intersect the reference value of 0, indicating there is no statistical difference in ES supply between that land cover and the reference one. In contrast, blue and red tones indicate cases where the 95% confidence interval for the log response ratio estimate does not intersect the reference value and where, consequently, the ES supply from the corresponding land cover is significantly better (blue) or worse (red) than the reference. The grey shading over each cell represents the number of studies contributing direct evidence on the corresponding land cover - ES combination, with darker tones reflecting fewer studies. For services marked with an asterisk (*), the effect of some land covers are likely to be masked by biophysical factors that have not been directly accounted for in our analysis. For example precipitation patterns will affect the supply of services related to water flow (water cycling, freshwater provision); soil type will affect nutrient cycling, primary productivity and soil formation, while precipitation intensity and slope (which also determines the location of some land covers) will affect erosion control.*

#### Supplementary Results 4. Detailed results from PERMANOVA analyses

Table S3: Detailed output of the permutational analysis of variance (PERMANOVA, *adonis* function with 200 permutations) on the effects of land cover characteristics on the supply of multiple ecosystem services. Model specified with production (production vs. non-production land covers) as the first variable term.

| Variable | Degrees of freedom | $F$ | Partial $R^2$ | $p$ - value |
|---|---|---|---|---|
| Production | 1 | 3.064 | 0.312 | **0.015** |
| Forest cover | 1 | 0.592 | 0.060 | 0.652 |
| Interaction (Production : Forest cover) | 1 | 1.159 | 0.118 | 0.318 |
| Residuals | 6 | | 0.509 | |
| Total | 9 | | 1.000 | |

Table S4: Detailed output of the permutational analysis of variance (PERMANOVA, *adonis* function with 200 permutations) on the effects of land cover characteristics on the supply of multiple ecosystem services. Model specified with forest cover as the first variable term.

| Variable | Degrees of freedom | $F$ | Partial $R^2$ | $p$ - value |
|---|---|---|---|---|
| Forest cover | 1 | 0.536 | 0.055 | 0.692 |
| Production | 1 | 3.119 | 0.318 | **0.025** |
| Interaction (Production : Forest cover) | 1 | 1.159 | 0.118 | 0.358 |
| Residuals | 6 | | 0.509 | |
| Total | 9 | | 1.000 | |

Table S5: Comparisons of group mean dispersions for variables on land cover characteristics. Separate tests were conducted for each variable (permdisp, *betadisper* and *permutest* functions with 999 permutations).

| Variable | $F_{(1,8)}$ | $p$- value |
|---|---|---|
| Production | 0.718 | 0.447 |
| Forest cover | 0.013 | 0.904 |

#### Supplementary Results 5. Data analysis with allocation of a single ES to each indicator

For our main analysis, we allocated each of the indicators in our dataset to as many ES as they were relevant, because some biophysical measures can have multiple values to humans. However, to understand how the sharing of indicators between different services affected our outcomes, we also conducted our analysis with each indicator allocated only to a single ES. In this section we present the results of this analysis - which are not to be confused with the results of our main analysis, presented in the main text and the preceding Supplementary Results sections.

Allocating each indicator to a single ES resulted in a dataset with information on the same 25 land covers used in our main analysis. Except for ethical and spiritual values (which uses indicators on culturally-significant food sources that are all shared with capture fisheries in our main analysis), the 17 ES present in the main text analysis were also present in the dataset for this analysis. However, restricting the allocation of indicators to a single service effectively decreased the total number of land cover comparisons: from 920 in the dataset used for our main analysis to 638 in the dataset used for the analysis we describe here.

Fig. S45 summarizes the estimated log response ratios for the land covers and ES that were part of this analysis. As in the main text analysis, these estimates were generated from individual random-effects network meta-analyses for each ES. The ES from the main analysis that are not included in this figure are ethical and spiritual values (which, as mentioned earlier, had no unique indicators) and water cycling. In this analysis, the evidence base for water cycling was formed by two separate subnetworks Fig. S46, the largest one of which did not include high producing exotic grassland. The results for this ES are therefore shown separately in Fig. S47, where low producing grassland is used as the reference land cover.

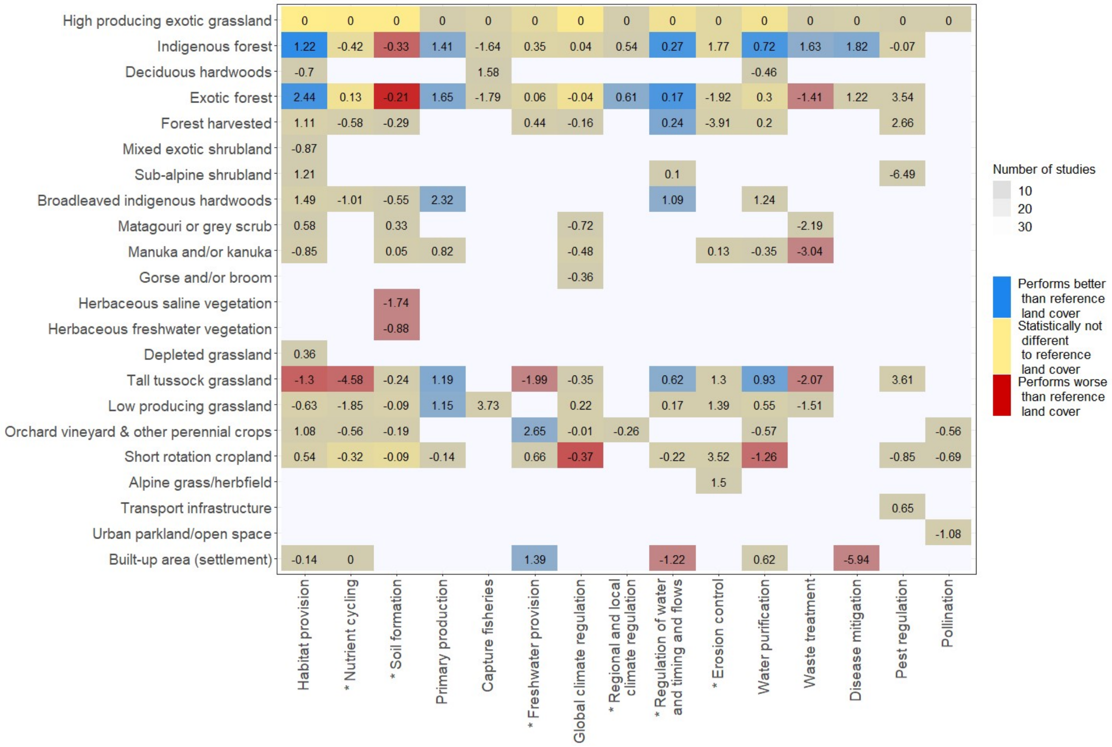


*Figure S45: Aggregated log response ratios of ES supply across land covers when indicators are allocated to a single ES. Values are given relative to the high producing exotic grassland reference. Yellow tones highlight cases where the 95% confidence intervals for a land cover's log response ratio estimate intersect the reference value of 0 indicating there is no statistical difference in ES supply between that land cover and the reference one. In contrast, blue and red tones indicate cases where the 95 confidence interval for the log response ratio estimate does not intersect the reference value and where, consequently, the ES supply from the corresponding land cover is significantly better (blue) or worse (red) than the reference. The grey shading over each cell represents the number of studies contributing direct evidence on the corresponding land cover - ES combination, with darker tones reflecting fewer studies. For services marked with an asterisk (*) the effect of some land covers can be masked by biophysical factors that have not been directly accounted for in our analysis.*

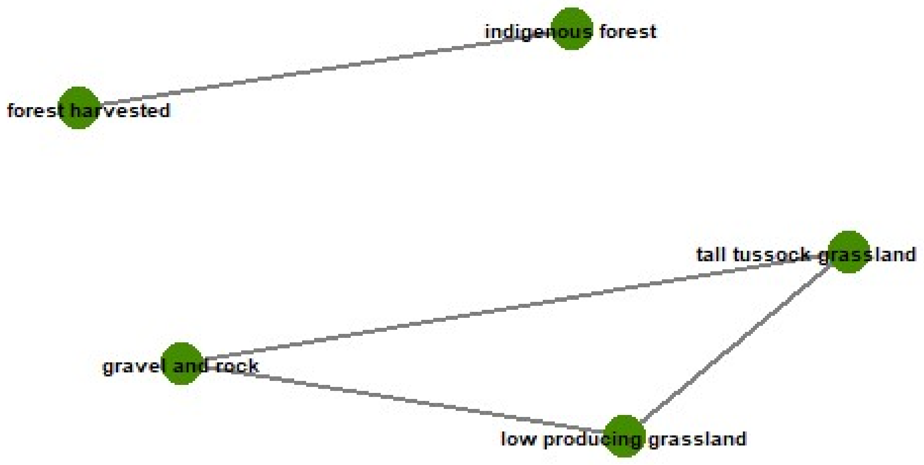


*Figure S46: Land-cover comparison networks for water cycling when indicators are allocated to a single ES.*

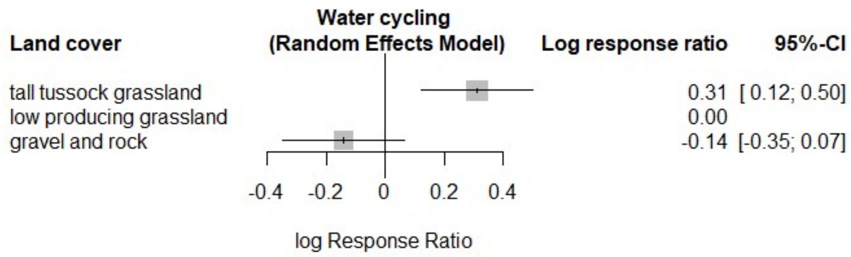


*Figure S47: Forest plot of land cover contrasts in the supply of water cycling when indicators are allocated to a single ES.*

Overall, the general trends depicted in Fig. S45 are consistent with the ones we find in the corresponding results for our main analysis (Fig. S44). In both cases, we find that no single land cover supplies high levels of all ES and that several production land covers (e.g., orchard, vineyard & other perennial crops, short-rotation cropland and forest harvested) tend to perform similarly or worse than the high producing exotic grassland reference in the supply of most services. However, for primary production, freshwater provision and waste treatment we do find results here that contrast with those shown in the main text analysis (Fig. S44), and which can be seen in Fig. S45. For each of these services, there are fewer studies informing the network meta-analyses (from which the log response ratio estimates in Figs. S44 and S45 are generated) in this analysis than there are in the main text analysis. For primary production, the number of studies is reduced from 32 in the main text analysis to only 8 in this analysis, for freshwater provision this changes from 40 to 13 and for waste treatment it is only from 6 to 4 studies. In all three cases, the reduction in the number of studies is accompanied by rearrangements of the networks of land cover comparisons that support the network meta-analysis for each ES. With these rearrangements, many of the links we observe in Figs. S22, S16 and S31 for the main text analysis are lost for the analysis in this service (figures not shown). The reconfiguration in the evidence bases suggest that, for these 3 ES, the results for this analysis and the main text ones are likely not comparable.

The results for pollination, pest regulation, capture fisheries and disease mitigation are the same for this analysis and the main text analysis, since the studies in the evidence base for each of these services are the same in both cases (see Figs. S40, S29, S33 and S37 for the corresponding evidence networks). Other services, such as water purification, erosion control, global climate regulation and soil formation, show only slight differences between the main text analysis (Fig. S44) and this one (Fig. S45). In these cases, the number of studies and the configuration of the evidence base is also similar in both analyses.

Following the same approach used in the main text analysis, here we also used clustering methods to examine how land covers compared against each other in their supply of ES and how ES compared against each other in terms of land covers that supplied them. For this, we selected a subset of the matrix in Fig. S45 that had no gaps and contained the largest possible number of total cells. This was a matrix of seven ES and seven land covers. We used the distances between ES in this subsetted matrix as inputs for the hierarchical clustering and the k-means cluster analysis comparing ES. For the analysis comparing land covers, we transposed the subsetted matrix and used this to calculate the distances required for the clustering analysis.

Figure S48 shows the results of the ES clustering analysis. Similar to the main text results, in this case we also observe that several services each form a cluster of their own and are, therefore, likely supplied in a distinct way by the seven land covers that were used in this analysis. Erosion control (which did not have enough data to be included in the subsetted matrices used in the main analysis), forms a cluster of its own that is separated from all other ES at the highest point in the height axis in Fig. S48, suggesting that it is supplied best by land covers that differ the most from those that best supply the other services in this analysis. The remaining services in Fig. S48 were all included in the main analysis and, with the exception of nutrient cycling and freshwater provision, they all form similar clusters to the ones shown in the results for the main analysis (Fig. 1, main text).

In the dendrogram for the main analysis (Fig. 1, main text), both nutrient cycling and freshwater provision are situated very close to services with which they share a large proportion of indicators: global climate regulation and water purification, respectively (see Supplementary Dataset 1 for shared indicators). In this analysis nutrient cycling forms a cluster of its own, while freshwater provision maps next to regulation of water and timing of flows (Fig. S48), suggesting that the clustering we observe for both of these services in the main analysis likely reflects their shared indicators with other services, as mentioned in the main text discussion.

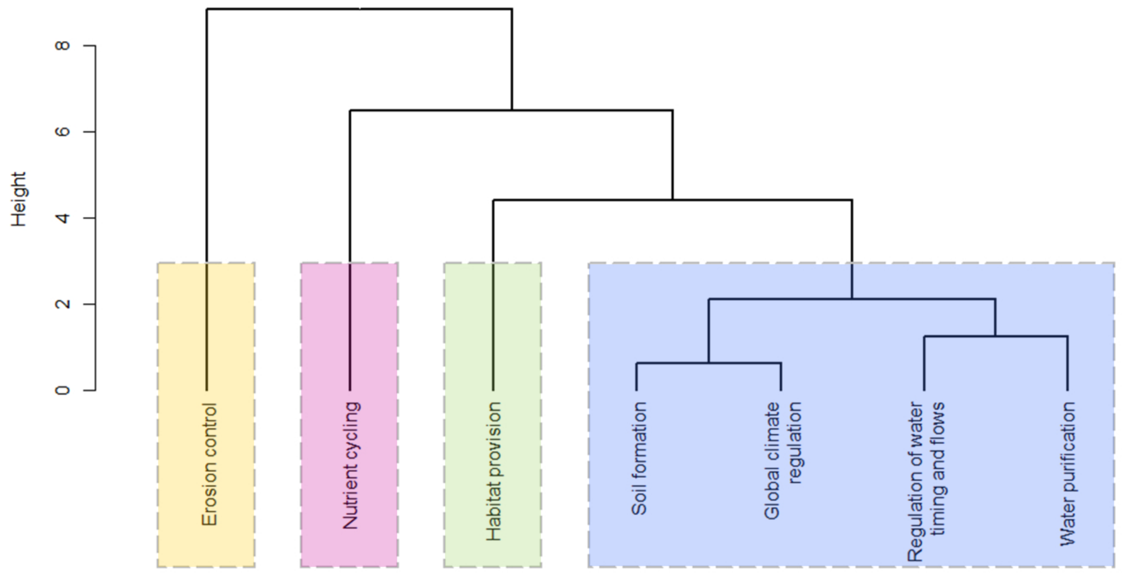


*Figure S48: Hierarchical clustering of ES when indicators are allocated to a single ES. Services that cluster within the same box were found (by k-means cluster analysis) to be supplied similarly across seven land covers (low producing grassland, tall tussock grassland, high producing exotic grassland, short - rotation cropland, indigenous forest, exotic forest and harvested forest). A greater separation between the branching points for clusters along the height axis indicates greater dissimilarity among clusters in the extent to which they are supplied by the seven land covers included in the analysis.*

Overall comparisons of land covers in terms of the ES they supply yielded similar results in this analysis as they did in the main text. In both cases, tall tussock grassland and short-rotation grassland each form separate clusters from the other services (although in the main text analysis tall tussock is paired with manuka and/or kanuka which is not included in this analysis). Similarly, exotic forest and harvested forest cluster together in both analyses, as do indigenous forest and low producing grassland (Fig. S49 and Fig. 2, main text). For this analysis, however, we did not include orchard, vineyard and other perennial crops which, in the main analysis, formed a cluster with high producing exotic grassland (Fig. 2, main text). In Fig. S49 all production and non-production land covers are segregated into different clusters, except for high producing exotic grassland, which clusters with indigenous forest and low producing grassland. However, this association is lost when erosion control (the ES with the most dissimilarities in its supply across multiple land covers for this analysis, Fig. S48) is removed from the analysis (Fig. S50).

The wide confidence intervals we observe in the forest plots (sensu Lewis & Clarke, 2001) for erosion control in both the main text analysis (Fig. S27) and this one (Fig. S51) suggest that there is high variability in the individual estimates for the supply of this service for each land cover. Since there are 22 studies supporting the erosion control network meta-analysis in the main text and 18 in this analysis, it is likely that this variability is not due to low replication, particularly for the land covers that are best represented within each analysis (high producing exotic grassland, and indigenous, exotic and harvested forest, Fig. S26). Instead, we expect this variability to reflect the effect that factors other than land cover (e.g., slope, soil type, weather and management practices) have on the supply of erosion control within each land cover type. These effects were not accounted for in our analysis and could potentially explain why including erosion control in the hierarchical clustering for this analysis results in the clustering of high producing exotic grassland and indigenous forest that we observe in Fig. S48.

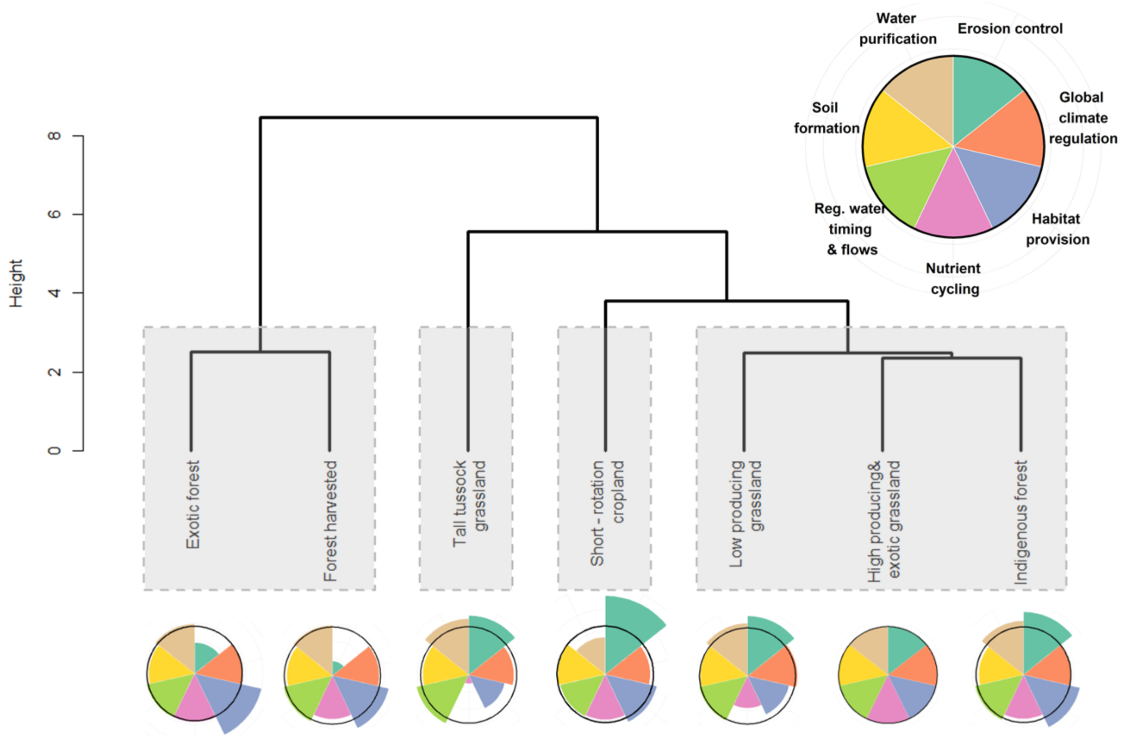


*Figure S49: Hierarchical clustering of land covers when indicators are allocated to a single ES. Boxes enclose land covers that exhibit greater similarity in their supply of seven ecosystem services (erosion control habitat provision, soil formation, nutrient cycling, water purification, global climate regulation and regulation of water timing and flows). The flower diagrams at the bottom illustrate how each land cover supplies each of the seven ES, with longer petals indicating a greater supply of an ES. For comparison, the black ring around each flower diagram marks the supply from high producing exotic grassland, the land cover used as a reference in our meta-analysis.*

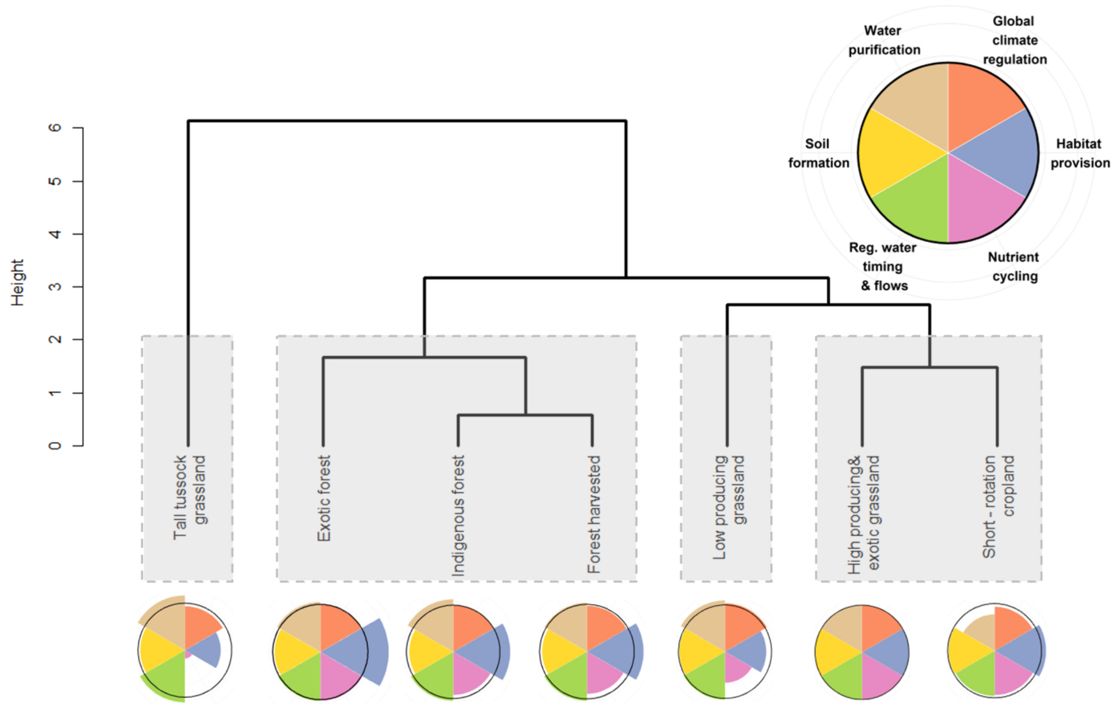


*Figure S50: Hierarchical clustering of land covers when indicators are allocated to a single ES and erosion control is excluded from the analysis.*

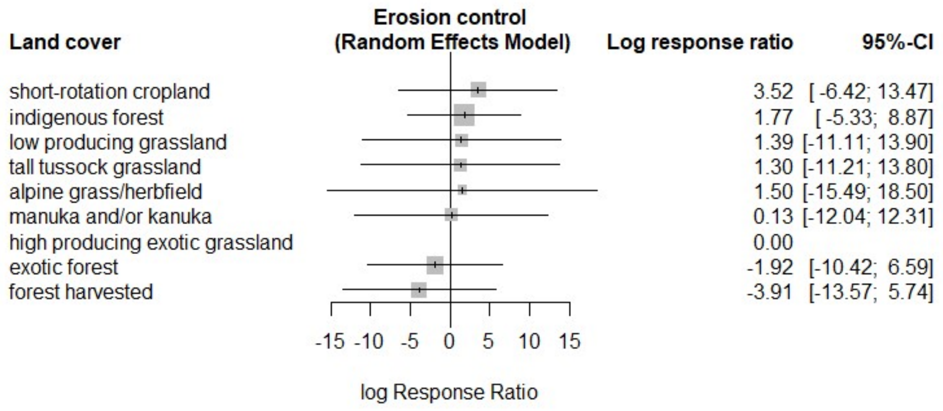


*Figure S51: Forest plot of land cover contrasts in the supply of erosion control when indicators are allocated to a single ES.*

Like in the main text analysis, here we conducted a PERMANOVA to test whether forest cover and the production explained the trends observed on Fig. S48. For this, we used the distances between land covers in the matrix with seven ES and seven land covers. Land covers were assigned to the same groups (forest / no forest and production / no production) they had in the PERMANOVA for the main analysis (Table S4). Unlike the main analysis, here we did not use a restricted number of permutations to calculate the significance values, since the matrix was small enough to allow for the calculation of all permutations (5,039 in total). Table S9 and Table S10 show that regardless of the order in which it was included in the model, the variable separating production and non-production land covers was not statistically significant in explaining the differences in ES supply between land covers. In contrast, the trade-off in ES supply between forest and non-forest land covers was statistically significant, but only when this term was added first in the model. In both cases, the assumption of homogeneous dispersion between both groups was met (Table S9, Table S10).

These results differ from the ones in our main analysis where we found the production vs. non-production dichotomy to be statistically significant regardless of the order it occupied in the model. Relative to the main analysis, here we retain the same number of groups but use a smaller matrix, with fewer land covers per group to test our hypothesis. The resulting decrease in power could be limiting our ability to detect the effect of production. Moreover, the two land covers that were not included in this analysis (manuka and/or kanuka and orchard, vineyard and other perennial crops) occupy extreme positions in the hierarchical clustering in the main analysis (Fig. 2, main text). Since each of these land covers occupied different groups in the production/non production classification, their exclusion from this analysis could mean that the differences between these groups are also reduced in this analysis relative to the main one. Lastly, these results could also signal that some of the indicators that were allocated to multiple ES in the main analysis presented large differences in ES supply between production and non-production land covers and that, when allocated to a single indicator, these differences were represented in fewer ES in this analysis. This would include several indicators on the leaching of nitrogen- and phosphorus-based compounds from terrestrial to aquatic systems. In the main analysis, these indicators were allocated to freshwater provision, water purification and nutrient cycling in but only water purification in this analysis. Likewise, indicators on coliform contamination in streams were allocated to freshwater provision, water purification and disease mitigation in the main analysis but only to disease mitigation in this analysis.

Table S6: Detailed output of the permutational analysis of variance (PERMANOVA, *adonis* function with 5,039 permutations) on the effects of land cover characteristics on the supply of multiple ecosystem services when indicators are allocated to a single service. Model specified with production (production vs. non-production land cover) as the first variable term.

| Variable | Degrees of freedom | *F* | Partial $R^2$ | *p* - value |
|---|---|---|---|---|
| Production | 1 | 3.898 | 0.284 | 0.067 |
| Forest cover | 1 | 3.935 | 0.287 | 0.068 |
| Interaction (Production : Forest cover) | 1 | 2.900 | 0.211 | 0.108 |
| Residuals | 3 | | 0.218 | |
| Total | 6 | | 1.000 | |

Table S7: Detailed output of the permutational analysis of variance (PERMANOVA, *adonis* function with 5,039 permutations) on the effects of land cover characteristics on the supply of multiple ecosystem services when indicators are allocated to a single service. Model specified with forest cover as the first variable term.

| Variable | Degrees of freedom | $F$ | Partial $R^2$ | $p$ - value |
|---|---|---|---|---|
| Forest cover | 1 | 4.994 | 0.364 | **0.029** |
| Production | 1 | 2.838 | 0.207 | 0.122 |
| Interaction (Production : Forest cover) | 1 | 2.900 | 0.211 | 0.108 |
| Residuals | 3 | | 0.218 | |
| Total | 6 | | 1.000 | |

Table S8: Comparisons of group mean dispersions for variables on land cover characteristics. Separate tests were conducted for each variable (permdisp, *betadisper* and *permutest* functions with 999 permutations).

| Variable | $F_{(1,5)}$ | $p$- value |
|---|---|---|
| Production | 0.684 | 0.459 |
| Forest cover | 0.003 | 0.915 |

#### Captions for datasets S1 to S5

**Supplementary Dataset 1:** Overview of Ecosystem Services (ES). Service categories and descriptions follow the classification presented in the Millennium Ecosystem assessment - MEA. Ecosystems and human well-being: Synthesis. (Island Press, 2005).

**Supplementary Dataset 2:** Overview of land cover classes as defined in New Zealand's Land Cover Database (LCDB) - Thompson, S., Grüner, I. & Gapare, N. Illustrated Guide to Target Classes of the New Zealand Land Cover Database Version 2. (2003).

**Supplementary Dataset 3:** Quantitative indicators used to quantify supply of each ecosystem service. For each indicator the units used by different studies are given. Indicators that lack units are reported as index or ratio in the units column. If the indicator is a variable that was logged, units of the variable before applying the logarithm are generally given. The "Ecosytem Service" 1 to 4 columns show the different services to which each indicator was allocated in our main analysis. "Ecosystem Service unique allocation" states the unique service to which each indicator was allocated for the supplementary analysis where indicators were matched to only one service. We present our reasoning for this single allocation in the "Rationale" column and provide references that support it on the "Supporting evidence" column. To support file sharing, special characters have been omitted from the letters in the last column.

**Supplementary Dataset 4:** Reference list for the studies included in our meta-analysis. The Study ID values are not sequential because they correspond to the unique identifier that each study received at the start of the literature screening process. The Study ID can be used to link the values presented in Supplementary Dataset 5 to the bibliographical reference of each study.

**Supplementary Dataset 5:** Final log Response Ratios on ecosystem service supply for pairwise comparison of land covers in each study used in our analysis. Within each study, log Response Ratios of multiple indicators for the same ecosystem service have been aggregated to a single value per service for each pairwise land cover comparison of land covers. Column heading abbreviations: Ecosystem service (ES), Unique study identifier (S.ID), Log Response Ratio (LRR), Variance (Var) and Standard Errror (SE). Bibliographical details for the studies linked to each unique identifier are provided in Supplementary Dataset 4**.**
